# The latent simplicity of microbial ecological interactions

**DOI:** 10.64898/2026.07.30.741683

**Authors:** José Camacho-Mateu, Isabel Quirós-Rodriguez, Giulio Burgio, Andrea Arrabal, Alfonso Mendaña, Miguel D. Fernandez-de-Bobadilla, Belén Benítez-Domínguez, Álvaro Sanchez

## Abstract

Microbial communities carry out functions of ecological, clinical and industrial importance. These functions arise from individual species contributions and from pairwise and higher-order interactions, whose number grows exponentially with community size. Yet community function is often surprisingly predictable. Here we show that this predictability does not require weak interactions. Instead, interactions across orders are strongly coordinated: those involving a given species are approximately proportional to related interactions one order below. This coordination causes species effects to vary together across community backgrounds, allowing function to be described by only one or two collective variables. We demonstrate this organization across dozens of combinatorial bacterial experiments and multiple environments. For total biomass, consumer–resource models and experiments identify the dominant collective variable with species yield, a trait that remains stable as abundances change through community dynamics. Microbial communities are therefore predictable not because their interactions are simple, but because their complexity is coherently organized. Our work provides a framework for understanding and ultimately designing community function from species traits.

## Introduction

Microbial communities perform functions that are essential in natural and engineered ecosystems, from nutrient cycling in soils^1^ to the production of metabolites^2,3^, pharmaceuticals and biofuels^4,5^. These functions are often collective: the contribution of any one species depends on which other species are present. Species compete for shared resources, exchange metabolites and alter the environment they inhabit^6–9,10–13^, and these effects are themselves modified when further species are added, producing the higher-order interactions widely documented in microbial consortia^14–17^. Because the number of such interactions grows exponentially with community size, predicting function from composition would seem to require measuring an exponentially large space of community states. Thus, rationally designing a community to optimize a desired function would appear to be intractable^18,19^.

Yet this combinatorial barrier may be less prohibitive than it appears. Accumulating evidence indicates that microbial communities often require only a small number of parameters to describe and predict their function^10,20–23^. One such regularity is that the effect of adding a species on the function of the community (its “functional effect”) often depends linearly on the function of the background community it is added to (Fig. 1A,B), a relationship known as a Functional Effect Equation (FEE)^21,24,25^. If, as we argue above, the functional effect of a species is itself the result of many pairwise and higher-order interactions with other members of the community, how can such simplicity in the effect of a species coexist with such complexity in the interactions that produce it?

**Figure 1.**
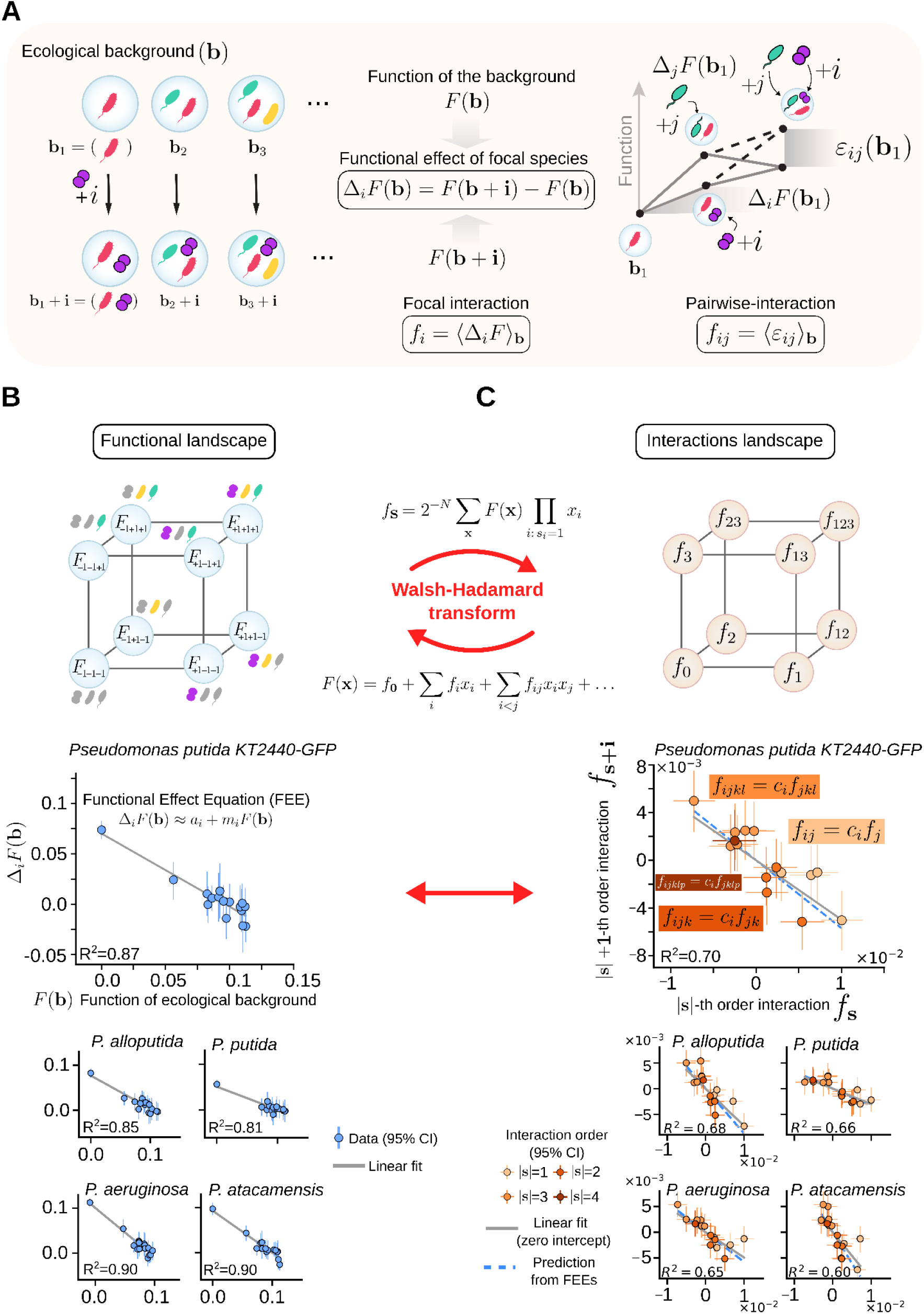
Functional Effect Equations imply linear relationships among ecological interaction coefficients. **(A)** For a focal species *i*, the functional effect in an ecological background ecological background ***b*** is defined as the change in community function upon addition of that species, Δ_*i*_ *F*(***b***) = *F*(***b*** + **i**) − *F*(***b***). Averaging this effect across backgrounds gives the first-order interaction coefficient of the focal species, *f*_*i*_ = ⟨Δ_*i*_ *F*⟩_***b***_. A pairwise interaction *ε* quantifies how the functional effect of species *i* changes in the presence of another species *j*, and averaging this context-dependent change across backgrounds gives the pairwise interaction coefficient *f*_*ij*_ = ⟨*ε* _*ij*_ ⟩_**b**_. Higher-order interaction coefficients are defined analogously (Supplementary Text S2). **(B, C; Top)** The same community-function landscape can be represented either in ecological-function space or, after the Walsh-Hadamard transform, Eq. (2), in interaction space. Here, **s** is a binary vector indicating the species involved in an interaction, and |**s**| gives its interaction order. In this interactions representation, an exact FEE predicts that interaction coefficients involving a focal species *i, f*_**s**+**i**_, are proportional to the corresponding lower-order coefficients not containing that species, *f* _**s**_, with slope *c*_*i*_ = *m*_*i*_ */*(2 + *m* ) (Supplementary Text S3). **(B; Bottom)** FEEs measured in the biomass landscape of five bacterial species grown in citrate + acetate medium. Points show functional effects across community backgrounds with *95%* confidence intervals, gray lines show linear fits. All focal species show strong FEE predictability, with *R*^2^ = 0. 81 to 0. 90. **(C; Bottom)** Interaction-coefficient relationships in the same biomass landscape. WH coefficients involving each focal species collapse against the corresponding lower-order coefficients not containing that species. Dashed blue lines show the slopes predicted from the FEEs, and gray lines show empirical linear fits. The interaction-coefficient relationships are strongly linear across focal species, with *R* = 0. 60 to 0. 70. The corresponding analyses for biomass in the remaining resource environments and for pyoverdine production are shown in Supplementary Figs. S4–S9.

In this paper we address this question and show that the two are reconcilable only if species interactions are not independent of one another: a strong FEE for a focal species implies that its higher-order interactions are proportional to the corresponding lower-order ones, with the constant of proportionality set by the slope of the FEE. We test this prediction experimentally and show that it holds across functions and environments. Furthermore, we show that this latent simplicity among interactions gives rise to an emergent simplicity in community function. If the interactions involving each species are proportional to one another, different species cannot affect the same community in idiosyncratic ways: their effects must rise and fall together as the background community changes. Community function is then governed by just one or two ecological collective modes: weighted combinations of species whose functional effects change together across backgrounds, even when the underlying interactions are strong and context-dependent.

The collective modes are inferred from the landscape itself, which leaves open what they represent biologically. Using consumer-resource models and biomass as a model function, we show that the dominant mode typically corresponds to a vector of species traits: the growth yield, or biomass produced per unit of resource consumed. We confirm this experimentally across 31 resource environments. Because yield is a property of a species’ metabolism rather than of its abundance, the dominant mode should not be affected by population dynamics: we find it to barely change over ten daily passages, even as species abundances diverge from the inoculum as communities move towards a stable state. The community-function landscape is therefore strikingly robust to population dynamics. Our results reveal a latent simplicity in microbial ecological interactions: community functions are not simple because interactions are weak, but rather because interactions are proportional to each other and contribute coherently through a small number of collective coordinates, which can be mechanistically interpretable.

## Results

### Pairwise and Higher-Order Ecological Interactions Follow Simple Statistical Patterns

To study how interactions shape community function, we represent the mapping between community composition and function as a *community-function landscape*^19,23,26^. Each community corresponds to a binary vector **x** ∈ {− 1, + 1} ^*N*^, where entries *x* _*i*_ =+ 1 and *x* _*i*_ =− 1 indicate species presence and absence, respectively. The community-function landscape is then a function *F*(**x**) assigning a quantitative output, such as biomass or a metabolite production, to each community composition (Fig. 1B, and Supplementary Text S2).

Previous work has shown that the functional effect of adding species *i* to a background community can often be predicted from the function of that background through a simple linear relationship (see Fig. 1B)^21^. Specifically, if Δ _*i*_ *F*(**x**) denotes the change in function caused by adding species *i* to background **x** (see Fig. 1A), then

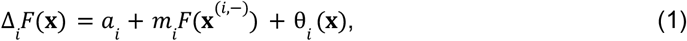

where *a* _*i*_and *m* _*i*_ are species-specific coefficients, and θ _*i*_ (**x**) captures deviations from the linear trend. We refer to these relationships as Functional Effect Equations (FEEs) (Fig. 1B, and Supplementary Text S3)^21,23^.

The functional effect of a species on a community emerges from pairwise and higher-order interactions with the resident community members^21^. Thus, we reasoned that the success of FEEs should reflect a latent, simple underlying organization of interactions at all orders. We tested this hypothesis by rewriting each community-function landscape in interaction space using the Walsh-Hadamard (WH) transform^18,27,28^ (see Supplementary Texts S2-S3 for a full description). The WH transform decomposes *F*(**x**) into additive, pairwise and higher-order coefficients,

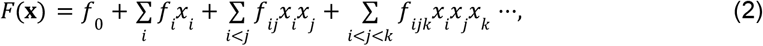

providing a one-to-one map between measured community functions for all 2^*N*^ communities in the landscape and the 2^*N*^ interaction coefficients at all orders (Fig. 1 B,C). As we show in the Supplementary Text S3, this representation yields a simple prediction. If the FEE for species *i* is exact, then WH coefficients involving *i* are recursively related to coefficients one order below (Fig. 1C, and Supplementary Texts S2 and S3)

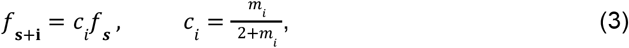

where ***θ*** is a binary vector denoting a subset of species not containing *i*, and **i** is the unit vector that adds species *i* to that subset. Thus, the slope *m* _*i*_ of the FEE predicts the slope *c* _*i*_ of a linear relationship in interaction space: pairwise coefficients involving *i* should scale with the corresponding additive coefficients, third-order coefficients involving *i* with pairwise coefficients not containing *i*, and so on, in all cases obeying the same linear law. Eq. (3) is exact only in the limit of an exact FEE. For empirical FEEs, this proportionality is approximate: deviations from the predicted line are determined by the residuals of the FEE (Supplementary Text S3).

To test this prediction we need a fully sampled community-function landscape. Because complete factorial landscapes are experimentally demanding, existing datasets are scarce and often have limited replication. This is a critical limitation for testing the predicted linear law amongst interaction coefficients, as high-order coefficients are especially sensitive to measurement noise^18^. We therefore constructed complete landscapes comprising all possible combinations (all monocultures, pairs, trios, four-member and the five member community) of five bacterial species (*Pseudomonas aeruginosa PA14 (PA), Pseudomonas putida KT2440-GFP (KT), Pseudomonas atacamensis (P1), Pseudomonas putida (P2)* and *Pseudomonas alloputida (P3)*). Each community was constructed in four independent biological replicates. Communities were inoculated by introducing all present species at an *OD* = 0. 001, and then grown for 24 *h* at *T* = 32ºC in 400 μ*L* of minimal carbon-limited M9 media supplemented with citrate and acetate as carbon sources, each at 0. 014 *Cmol/L* ( 0. 028 *Cmol/L* total). This medium is similar to those that have been previously used to construct synthetic community-function landscapes^21,22,24,29^. For each community, we measured two different community functions: optical density at 600*nm*, as a proxy for biomass, and absorbance of the filtered supernatant at 405*nm*, as a proxy for pyoverdine concentration^30,31^ (see Methods and Supplementary Text S1 for details). This experiment allowed us to determine the function *F*(**x**) for each community composition **x**.

We first analysed the biomass landscape. Consistent with similar findings in other community-function landscapes^21,24,25^, all species exhibited strong FEEs, with *R*^2^ values above ∼ 0. 8 (Fig. 1B). To test the theoretical prediction that a strongly predictive FEE for the focal species *i* would imply the existence of a strong linear association between interactions *f*_**s**+**i**_ and *f*_***s***_, we directly determined all interaction coefficients through the Walsh-Hadamard transform of *F*(**x**) (Fig. 1B). Consistent with our theoretical expectation (Fig. 1C), strong FEEs with *R*^2^∼0. 8-0. 9 do imply that all interaction coefficients are strongly related to one another, following approximate linear relationships, with *R*^2^∼0. 6 to ∼0. 7, and slopes consistent with those predicted by Eq. (3). Thus, interactions involving a focal species are not independent across orders, but are statistically constrained by lower-order coefficients. Repeating the same analysis for pyoverdine production yielded qualitatively similar results (Supplementary Fig. S4).

To test the generality of this comparison between theory and experiment, we repeated the entire experiment in a wide range of different growth conditions. Previous work has established that species interactions are sensible to the physico-chemical environment in which species grow^6–9^. Because the dominant form of species interactions in experiments like ours is resource competition and cross-feeding^32^, we systematically manipulated the resource composition of the environment, by creating all 31 combinations of 5 different carbon sources: *citrate, acetate, fructose, glucose and glycerol* (Supplementary Text S1). In each of these 31 different growth media we constructed our full factorial community-function landscape in four biological replicates exactly as described above, and measured the same two functions. The good match between theory and experiment was consistent across all environments (Fig. S5-S9), indicating that the existence of simple linear equations constraining interactions at all orders is a robust feature of our community-function landscapes.

To assess to what extent this finding generalizes beyond the interactions that take place in our specific experiments and functions, we re-analyzed six previously published experiments with publicly available data^15,18,24,29,33,34^ (Supplementary Table T2). These datasets included both functions similar to those measured in our experiments, such as biomass, and functions extending beyond them. The latter included the population sizes of individual community members in a recent full-factorial study of plant-associated communities^33^, which quantify how the presence of one species affects the abundance of another and therefore connect more directly to the population-dynamical interpretation of ecological interactions. We also included community-level functions as diverse as life-history traits of fruit flies grown axenically with every possible combination of five different members of their native microbiome^34^. Although these prior experiments lacked sufficient replication to estimate higher-order interactions reliably^18^, first-, second-, and third-order interaction coefficients frequently followed the same linear co-dependencies observed in our experiments (Supplementary Figs. S24-S26). This suggests that our empirical findings extend beyond our particular experimental system.

### Correlated interactions produce ecological collective modes and low-dimensional community-function landscapes

Complexity is often associated with systems that exhibit dense pairwise and higher-order interaction networks. Yet, the fact that interactions at all orders are strongly dependent on each other in community-function landscapes (and that the functional effects of species follow simple linear trends) suggests that such landscapes may be simpler than one might expect based on how many interactions they involve. To formalize this intuition, we construct the vector of functional effects,

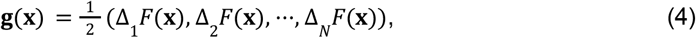

which captures the effect that every species has on function *F* on background **x**, e.g. Δ_1_ *F*(**x**) = *F*(1, *x*_2_, *x*_3_,…) − *F*(− 1, *x*_2_, *x*_3_,…). From Eq. (1), we see that Δ*i F*(**x**) is affected by the interactions at all orders involving the focal species *i* as well as every other member of the community (Supplementary Text S3). If these interactions were all independent, species would produce unrelated functional effects on the same background. By contrast, the effects of different species on the same background are coordinated rather than idiosyncratic when interactions are proportional to one another (Supplementary Text S4-S5). Whether the functional effects of different species vary independently or coherently across backgrounds can be captured by the following matrix (Fig. 2 A, and Supplementary Text S4)

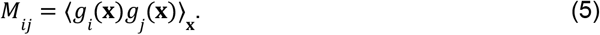

The eigenvectors of **M, *u*** _α_, α = 1, ···, *N*, define *ecological collective modes*: linear combinations of species along which functional effects are coherent. If the landscape is effectively low-dimensional, the spectrum of **M** should be dominated by a few leading eigenmodes (Fig. 2A). These modes also provide natural coordinates for compressing the original high-dimensional landscape. Projecting a community **x** onto an eigenmode ***u*** _α_ defines a latent coordinate,

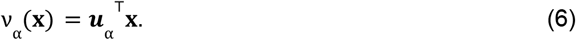

When one mode dominates, community function can be approximated by a one-dimensional non-linear map, *F*(**x**) ≈ *q*(*ν*_1_). Importantly, as we show on the Supplementary Text S5, this map is a latent projection that can be theoretically calculated from the leading eigenmode ***u***_1_, the leading eigenvalue λ_1_, and the first-order terms (*f*_*i*_) in the WH expansion of the landscape. In practice, we can evaluate this projection using the second-order truncation of the expansion, motivated by previous work showing that first and pairwise terms capture most of the variance in microbial community-function landscapes ^18,22^ (Supplementary Text S2 and Supplementary Figs. S2-S3).

**Figure 2.**
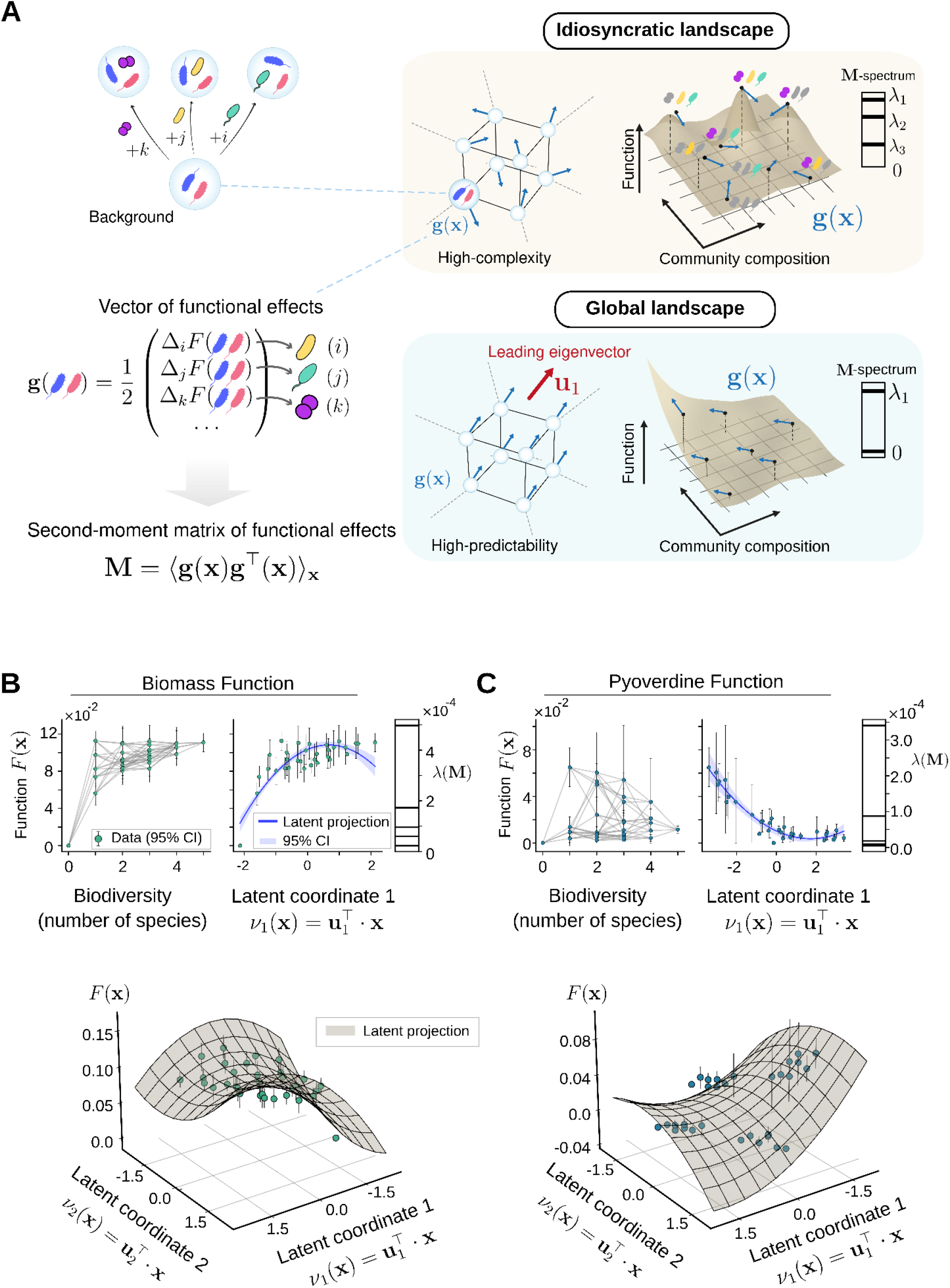
Correlated ecological interactions generate low-dimensional collective modes in community-function landscapes. **(A)** For each community background, the functional effects of adding each focal species define a functional-effect vector **g**(**x**), **Eq. (4)**. The second-moment matrix of these vectors,, quantifies how the functional effects are coordinated across **M** = ⟨***g***(**x**)***g***^⊤^ (**x**)⟩ _**x**_ backgrounds. Idiosyncratic landscapes produce functional-effect vectors pointing in many directions and a broadly distributed λ(**M**)-spectrum, whereas structured or global landscapes have aligned functional effects and a spectrum dominated by one or a few collective modes. **(B)** Biomass landscape measured in *citrate + acetate* medium. Projection onto community biodiversity (computed as the total number of species) captures the overall increase in biomass with the number of species, but leaves substantial variation among communities with the same biodiversity. Projection onto the leading eigenmode ***u*** reveals a latent coordinate, *ν*_1_(**x**) = ***u***_1_^⊤^ · **x**, that captures the dominant trend. The spectrum of **M** is strongly concentrated: the first eigenvalue, accounts for ∼77% of the total spectral weight and the first two for ∼96%. The second-order WH, **Eq. (2)**, projection onto *ν*_1_(**x**) yields a one-dimensional latent map, *q*(*ν*_1_), that captures the dominant trend with *RMSD* = 0. 013, or ∼11. 5% of the observed functional range (see Supplementary Text S5). Including the second collective coordinate, *ν*_2_(**x**), gives a two-dimensional latent surface, *Q* (*ν*_1_, *ν*_2_), that describes the structure of the landscape in the latent plane (see Supplementary Text S6). **(C)** Pyoverdine landscape measured in the same medium. Pyoverdine production is poorly explained by species biodiversity, whereas projection onto *ν*_1_ (**x**) reveals a clear functional collapse. Here, the first eigenvalue accounts for ∼80% of the total spectral weight and including the second eigenvalue for ∼98%. The latent map *q*(*ν*_1_) captures the empirical structure with *RMSD* = 0. 005, or ∼7. 6% of the observed functional range, and *Q*(*ν*_1_, *ν*_2_) provides a compact two-dimensional representation.

When a second mode (*ν*_2_ ) also contributes significantly, the approximation extends naturally to a response surface *F*(**x**) ≈ *Q*(*ν*_1_, *ν*_2_ ) (Supplementary Text S6). This surface can likewise be evaluated from the two leading eigenmodes of **M**, their eigenvalues, and the first and second-order interaction coefficients. Importantly, *Q*(*ν*_1_, *ν*_2_ ) can include nonlinear interaction terms between latent coordinates (Supplementary Text S6). Thus, the collective modes define a natural coarse-graining of the landscape, in which the effects of many species-level interactions are compressed into effective interactions among just two ecological collective modes.

Our experimental data support these predictions. For the community-function landscape represented in Fig. 1, assembled in citrate and acetate as the sole carbon sources, the spectrum of **M** was strongly concentrated in two leading collective modes (Fig. 2B,C). For instance, for the biomass function, the leading eigenvalue accounted for ∼77%, of the total spectral weight and the first two eigenvalues together accounted for ∼96% of it. In pyoverdine, the corresponding values were ∼80% and ∼98%. The leading mode was also clearly separated from the second, with λ_1_ */*λ_2_= 4. 15 for biomass and λ_1_ */*λ_2_ = 4. 55 for pyoverdine, indicating a dominant collective axis in both landscapes (Fig. 2B, C). Consistently, the stable rank of **M**^35^–a measure of the effective number of orthogonal dimensions of a matrix–was close to 1 in both cases (1. 06 for biomass and 1. 05 for pyoverdine), confirming that the functional-effect matrix is effectively low-rank (see Methods).

Unlike classical biodiversity–ecosystem function relationships, which summarize community structure primarily through species biodiversity (here computed as the number of species in a community; Fig. 2B, C)^29,36–46^, the first two modes captured species-specific patterns of functional covariation across community backgrounds. For biomass, we found close agreement between the theoretical curve *q*(*ν*_1_) and the experiments, with a root-mean-square deviation of *RMSD* = 0. 013, or ∼11. 5% of the observed functional range (Fig. 2B). The theoretically expected two-dimensional surface *Q* (*ν*_1_, *ν*_2_) also exhibited a strong agreement with experiments, describing the structure of the landscape in the latent plane with a *RMSD* = 0. 015. For pyoverdine, biodiversity was much less informative (Fig. 2C), yet the theoretical *q*(*ν*_1_) strongly predicted the empirical pattern with a *RMSD* = 0. 005, or ∼7. 6% of the observed functional range (Fig. 2C). Similarly, the theoretical surface *Q* (*ν*_1_, *ν*_2_) again provided a close representation of the empirical landscape, with *RMSD* = 0. 0099.

The resulting latent coordinates {*ν*_α_ (**x**)}_*α*_ are analogous to those used in global-epistasis models, where an observed phenotype is represented as a nonlinear function of an underlying additive trait^47,48^. Here, however, the latent trait is not fitted phenomenologically; rather, it emerges from the leading eigenmodes of the functional-effect matrix **M**. Thus, the existence of predictive FEEs, the fact that interactions at different orders are linked together through simple linear models, and the collapse of the community-function landscape into one or two latent-coordinates, are all different manifestations of the latent simplicity of the community-function landscape (Supplementary Text S7).

This low-dimensional architecture was consistently found in all the other 30 community-function landscapes described in the previous section, each assembled in a different combination of resources. Across all these environments, the first two collective modes consistently captured most of the spectral weight of functional-effects and were separated from the other modes by a pronounced spectral gap (Supplementary Figs. S10–S14). We found similar results for the other previously published full-factorial datasets (Supplementary Figs. S27-S31). Although coefficient-level relationships were often noisier, the spectra of **M** were typically concentrated in one or two collective coordinates, and the corresponding latent-coordinate projections provided coherent representations of community function (Supplementary Text S9). This suggests that the latent simplicity observed in our experiments is not an artifact of our particular empirical system, but may be a broader feature of microbial community-function landscapes.

Our results are indeed consistent with recent studies finding that community-function landscapes often have low effective dimensionality, with much of their variation captured by a few collective variables^10,20^. Being inferred through data-driven coarse-graining, dimensionality reduction, or machine learning^10–13^, these collective variables typically lack a clear biological interpretation. To help us answer this question, we must resort to mechanistic ecological models.

### Microbial yields give biological meaning to the leading ecological collective mode

To go beyond mere statistical findings, we turned to studying mechanistic models of our microbial communities. These allow us to control species-level traits and potentially connect them to community-level outcomes. Considering the case where the community function is total biomass, we studied Consumer-Resource models (CRMs) in which species compete for either one or multiple public resources^49–51^ (Fig. 3A; Supplementary Text S8) as a proxy for our experimental setup. Despite their simplicity, CRMs have proven to capture the behavior of microbial communities reared under similar conditions to the ones we have studied here^52–54^, and they exhibit many of the emergent properties of natural microbiomes^49–51^, ^52–54^. Furthermore, competition for a common resource has been recently proposed as a mechanism for the emergence of FEEs ^55^. In such systems, any functional effect or interaction is exclusively mediated by resource depletion, whereby adding a species changes the fraction of the common resources available to the background community. This provides a mechanistic route to coordinated functional effects and low effective rank in **M**.

**Figure 3.**
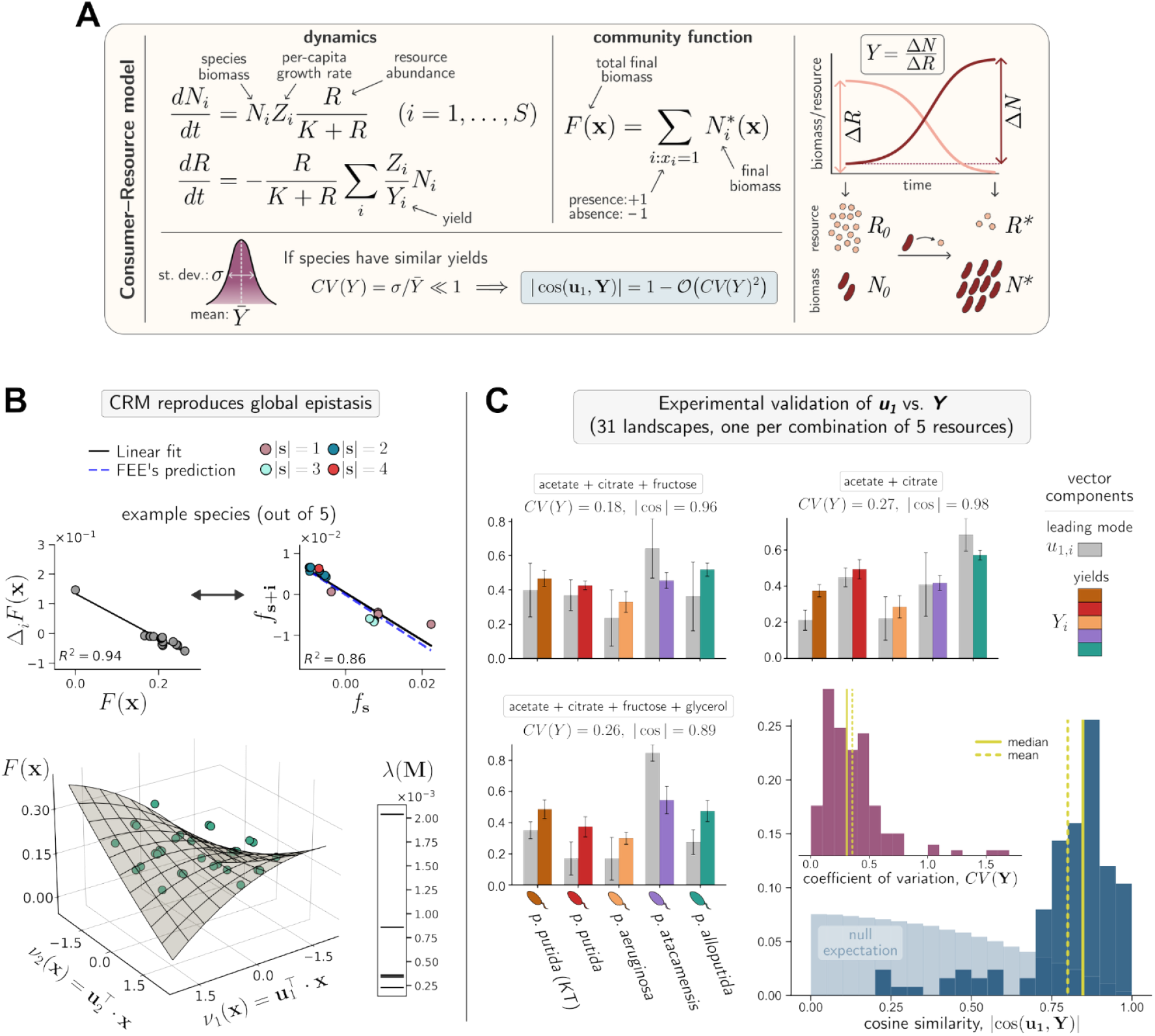
Microbial yields give biological meaning to the leading ecological collective mode. **(A)** Box illustrating the Consumer-Resource model (CRM) we used as a proxy for a community of *S* species competing for one (multiple) shared resource(s), together with our main analytical prediction. We predict that when yields are relatively similar, as quantified by a low coefficient of variation *CV*(***Y***), the leading collective mode ***u***_1_ of matrix **M** approximately aligns with the yield vector ***Y*** (|*cos* (***u***_1_, ***Y***) | approaches 1; see Supplementary Text S8 for derivation). **(B)** Results from a representative simulated landscape show that the model reproduces the existence of FEEs (for clarity, here shown for one species only) and of collective latent coordinates (***u***_1_,***u***_2_). **(C)** Comparison of ***u***_1_ versus ***Y*** (normalized) across the 4 biological replicates of the 31 empirical 5-species landscapes we built from all possible combinations of 5 carbon sources (124 data points). The distribution of cosine similarity between ***u***_1_ and ***Y*** (median: 0. 85; 95% CI: [0. 26, 0. 98]; skewness: − 1. 87) is strongly shifted toward high values compared to the null expectation between random normalized vectors in 5 dimensions (Kolmogorov-Smirnov test, *D*(124) = 0. 74, *p* < . 001). Correspondingly, *CV*(***Y***) is concentrated at low values (median: 0. 31; 95% CI: [0. 06, 1. 01]; skewness: 2. 34). The panel also reports the vectors ***u***_1_ and ***Y*** averaged over the 4 replicates (and their standard errors), broken down by components, for three different mixes of carbon sources. The displayed cosine similarities refer to the averaged vectors.

To study whether CRMs would display the same general principles of low effective dimensionality of community-function landscapes observed in our experiments, we simulated more than 2. 5 × 10^5^ landscapes, systematically accounting for across-species heterogeneity of traits (yield, ***Y***, and maximum per-capita growth rate, ***Z***) as well as correlations between traits. The full model details are sketched in Fig. 3A and thoroughly described in Supplementary Text S8. The simulations generally reproduce the global epistasis patterns found in experimental data, namely the existence of strongly predictive FEEs, the co-dependence of interactions at all orders following similarly simple trends as those observed empirically, and the collapse of the community-function landscape into one or two latent collective coordinates (Fig. 3B).

Given that CRMs exhibit a similar behavior to our experiments, but they are fully defined mechanistically, we set out to investigate whether we could mathematically explain how the leading mode is connected to species traits. As we elaborate in the Supplementary Text S8, it is possible to show that under empirically realistic limits the leading eigenvector ***u***_1_ of **M** is approximately parallel to the vector of yields, ***Y*** = (*Y*_1_, ···, *Y*_*N*_) (Fig. 3A). Our theoretical analysis proves that the alignment between the two vectors is primarily controlled by the heterogeneity among yields, as quantified by their coefficient of variation, *CV*(***Y***). When yields are narrowly distributed (*CV*(***Y***) is low), the leading collective mode is strongly aligned with the yield vector (|*cos*(***u***_1_, ***Y***)| approaches 1), independently of how per-capita growth rates (***Z***) are distributed across species. In this regime, the dominant eigenvector is thus mechanistically interpretable in terms of species yields.

This theoretical prediction was first successfully validated through simulations (Supplementary Fig. S16). To test whether, as suggested by the model, the first collective mode ***u***_1_ does indeed approximately represent the vector of species yields in our experimental communities, we estimated yields from species growth in monoculture in each resource environment (Supplementary Text S1) and then compared the empirical vector of yields with the leading eigenvector inferred from the empirical biomass landscapes. In Fig. 3C we show the comparison between ***u***_1_ and ***Y*** for three representative community-function landscapes with a small *CV*(***Y***). For all three landscapes we find that the leading eigenvector is very close to the yield vector (|*cos*(***u***_1_, ***Y***)| = 0. 89, 0. 96, 0. 98), in agreement with the prediction from the CRM. To assess whether this alignment was significant, we extended this analysis to the remaining 28 community-function landscapes in our previous experiments and compared the empirical cosine similarities against a random-vector null model (Supplementary Text S8). Figure 3C summarizes this comparison across all 31 resource environments, each replicated four times. We find that the empirical distribution of cosine similarities between ***u***_1_ and ***Y*** is strongly shifted toward high values compared to the null expectation, deviating significantly from it (Kolmogorov-Smirnov test, *D*(124) = 0. 74, *p* < . 001). Consistent with our theoretical predictions, the distribution of *CV*(***Y***) in our empirical landscapes is highly skewed toward low values (median: 0. 31; 95% CI: [0. 06, 1. 01]; skewness: 2. 34). This successful test of the theoretical prediction suggests that the leading collective mode in biomass functional landscapes can be directly interpreted from species traits, namely the metabolic efficiencies expressed as biomass production per unit carbon consumed.

This result requires the distribution of yields to be relatively narrow. If yields varied widely across species, the functional effects (and therefore the leading mode ***u***_1_) would become more sensitive to differences in resource-exploitation ability, which are governed by per-capita growth rates. However, as we show in Supplementary Text S8, when growth rates strongly correlate with yields, species form a competitive hierarchy that aligns ***u***_1_ to the vector of focal effects **f** = (*f*_1_, ···, *f*_*N*_), and **f** is in this case entirely determined by ***Y***. Therefore, within CRMs’ domain of validity, the leading collective mode can be accurately reconstructed from species traits also in this hierarchical regime.

### The leading biomass mode remains stable through population dynamics

An important caveat of the datasets analyzed in the previous sections is that they are all single-batch experiments. All species are always inoculated at an arbitrarily fixed initial density, which allows us to define the composition as a binary vector **x**. This implies, however, that the results will generally depend on the chosen inoculum density. One may argue that, if communities had been serially passaged instead, a vector containing presence/absence in the initial inoculum would not be informative of community function, as the composition will change at the onset of every passage until communities reach a stable state.

However, an intriguing hypothesis emerges from the results of the previous section. There, we found that the leading collective mode of biomass-production landscapes was strongly aligned to the vector of yields. The growth yield is a species trait related to its metabolic efficiency, which should be relatively robust to changes in the size of its population. Therefore, we hypothesized that the leading biomass mode (***u***_1_ ) should be roughly insensitive to the shifts in community composition that result from ecological interactions and population dynamics as communities approach a stable equilibrium.

To test this hypothesis, we followed a top-down community assembly approach to find a pool of five stably coexisting species in citrate (0. 07 *Cmol/L*) minimal M9 media under daily serial passaging (with a dilution factor 1: 100) every 24 *h* at 28°C. We found a community of five species that coexisted together under this regime: *Acinetobacter johnsonii, Serratia proteomaculans, Pseudomonas defluvii, Aeromonas hydrophila and Lelliottia amnigena* (Fig. 4A). We then assembled all 31 species assemblages we could form with these 5 species, by inoculating together every monoculture, pair, trio, 4-member and 5-member communities at equal initial abundances (Fig. 4A). We grew these assemblages in the same M9 medium supplemented with citrate as the sole carbon source and subjected them to the same daily serial-passaging regime used during the assembly stage (Methods and Supplementary Text S1). We measured total biomass every 24 *h* for 10 days in two independent biological replicates, with three technical replicates in one experiment and two in the other (Fig. 4A). This experiment generated a fully sampled community-function landscape where the function *F*(**x**, *t*) maps the function measured on day *t* of serial passaging to the presence/absence composition vector **x** on the day we inoculated our communities. Species extinctions were rare, and most (29*/*31) of the communities contained all inoculated species on day 10, even though species abundances differed among the communities and their composition diverged over time from that in the starting inocula (Supplementary Fig. S18).

**Figure 4.**
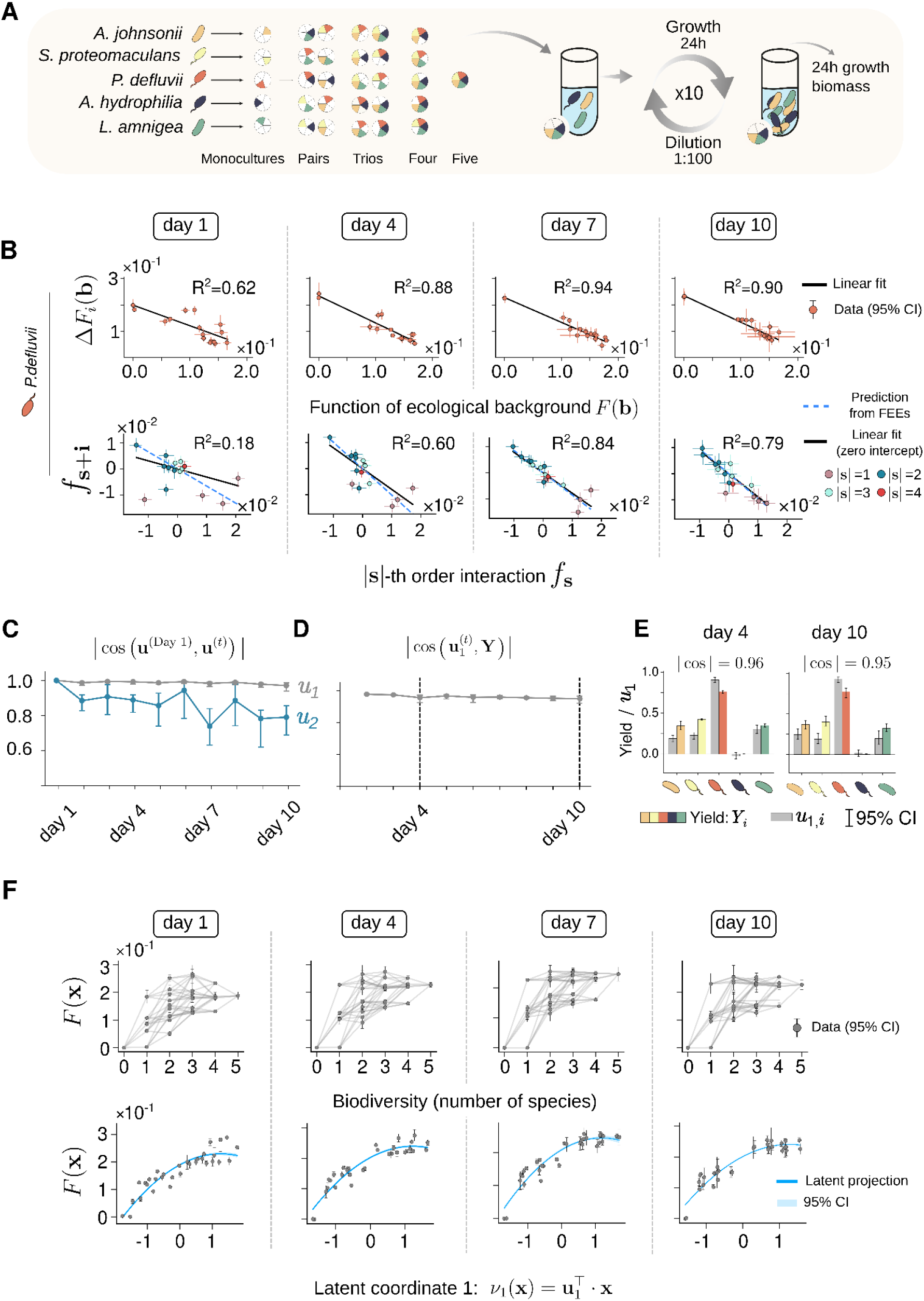
The dominant productivity mode is stable through population dynamics. **(A)** Experimental design. Complete biomass landscapes were reconstructed during serial dilution by assembling all (2^5^ − 1 ) non-empty communities from five stably coexisting species. Communities were grown for 24 *h* in citrate-limited M9 medium and serially diluted for 10 daily passages. Biomass was measured at each passage. **(B)** Statistical relationships organizing the biomass landscape for a representative focal species at selected passages (the complete analysis across all focal species and passages is shown in Supplementary Figs. S19-S20). Top: FEEs, Eq. (1), relate the functional effect of adding the focal species to the biomass of the background community. Bottom: interaction-space relationships compare interaction coefficients involving the focal species with the corresponding lower-order coefficients not containing it, as defined in Eq. (3). Both relationships show an initial transient and then become strongly linear through time, increasing from *R*^2^ = 0. 62 to *R*^2^ = 0. 90 for FEEs and from *R*^2^ = 0. 18 to *R*^2^ = 0. 84 for the interaction-coefficient relationship in the representative example shown. **(C)** Temporal stability of the first two collective modes, quantified as the absolute cosine similarity, |***u***_1_ (*Day* 1) · ***u***_1_ (*t*)|, between each mode at passage (*t*) and its orientation at the first passage (*t* = *Day* 1). The leading mode remains strongly conserved, with |***u***_1_ (*Day* 1) · ***u***_1_ (*t*)|∼[0. 95,1. 00], whereas the second mode is less conserved, |***u***_2_ (*Day* 1) · ***u***_2_ (*t*)|∼[0. 74,0. 78] at later passages. **(D)** Alignment between the leading mode ***u*** and the yield vector ***Y***, estimated from monoculture dynamics across passages (Supplementary Text S1). The high cosine similarity, |***u***_1_ (*t*) · ***Y***|∼[0. 94,0. 98], indicates that ***u***_1_ remains aligned with a productivity-related trait axis. **(E)** Species-level comparison between the components of ***u***_1_ and ***Y*** at representative passages. The two vectors remain closely aligned at both early and late passages, with |***u***_1_ (*t*) · ***Y***| = 0. 96 at day 4 and 0. 95 at Day10, showing that the conserved leading mode reflects species-specific productivity differences. **(F)** Biomass landscapes projected onto species biodiversity (computed as the total number of species) and onto the leading latent coordinate, *ν*_1_ (**x**) = ***u***_1_·^⊤^ **x**, across passages. Projection onto *ν*_1_ (**x**) reveals a stable functional organization of the biomass landscape through time. The corresponding projections for all daily passages are shown in Supplementary Figs. S21-S22.

Remarkably, we found that the binary community-composition vector **x**, which merely accounts for species presence/absence in the inoculum, remains highly informative of community function despite the underlying population dynamics. FEEs based solely on inoculum composition retained substantial predictive power throughout serial passaging, although their performance changed differently across species. For instance, for *P. defluvii*, the *R*^2^ of the FEE increased from 0. 62 after the first single batch (an experiment equivalent to those analyzed in Figs. 1-3), to 0. 88 after four days of serial passaging, remaining around 0. 9 on days 8-10 (Fig. 4B). A similar, although weaker and less monotonic, improvement was observed for *A. johnsonii* and *L. amnigena*, whereas predictive performance remained approximately stable or more variable for *S. proteomaculans* and *A. hydrophila* (Supplementary Figs. S19-S20).

Consistent with our hypothesis, the leading biomass mode of *F*(**x**, *t*), ***u***_1_ (*t*), barely changed over serial passaging and remained aligned with its day-1 orientation, with |*cos*(***u***_1_ (*Day* 1), ***u***_1_ (*t*))|∼ [0. 96, 0. 99] (Fig. 4C). By contrast, the second mode was less conserved, with |*cos*(***u***_2_ (*Day* 1), ***u***_2_ (*t*))|∼[0. 76,0. 89] at later passages (Fig. 4C). Also consistent with the foundation of our hypothesis, ***u***_1_ (*t*) remained strongly aligned with the yield vector throughout the experiment, with |*cos*(***u***_1_ (*t*), ***Y***)|∼[0. 95,0. 98] (Fig. 4D). This alignment was visible at the species level when comparing the components of ***u***_1_ (*t*) with species yields at representative early and late passages (Fig. 4E and Supplementary Fig. S23).

The strong conservation of ***u***_1_ (*t*) over time has an important consequence. If the leading collective coordinate, *ν*_1_ (**x**, *t*) = ***u***_1_ (*t*) · **x**, varies little over time, then the projection of the landscape, *F*(**x**, *t*) ≈ *q*(*ν*_1_(**x**, *t*)), should follow suit, revealing a temporally stable organization of biomass production (Fig. 4F). Accordingly, the projection of *F*(**x**, *t*) on ***u***_1_ after a single batch would provide a reasonable approximation of this projection even after several days of serial passaging. As we observed in our experiments, this holds true despite the population dynamics clearly causing the landscape to collapse into an apparently bimodal distribution of biomass over time (Fig. 4F, top panel). Thus, while community dynamics will push different communities towards different stable equilibria, and shift species abundances over time, the leading collective mode will remain stable and continue to organize variation in biomass production across community backgrounds.

### Discussion

The functions of microbial communities emerge from pairwise and higher-order interactions among their members. Simplicity in microbial community function has generally been understood as an absence of interactions: function is predictable when higher-order effects are weak enough to neglect^22^, or sparse enough that few of them matter^17^. In the landscapes analyzed here, interactions are reasonably strong at every order and function is nonetheless governed by one or two variables, because interactions at different orders are not independent from each other. For each species, its higher-order interactions are proportional to the corresponding lower-order ones, with a constant of proportionality fixed by the slope of its Functional Effect Equation^21^. Therefore, community functions are simple not because interactions are absent, but because interactions at all orders are proportional to one another.

This finding suggests that the number of interactions in a community provides a poor measure of its complexity, which is set instead by the number of independent ways in which its species affect function. In the dozens of community-function landscapes studied here this number is essentially one or two, each corresponding to a combination of species whose functional effects vary coherently across community backgrounds. These collective modes are inferred from the functional effects they organize, which leaves open what they represent biologically. Variables of this kind are ordinarily obtained by fitting^47,48^ or by data-driven dimensionality reduction^10,20^, and are correspondingly difficult to interpret. As we have shown for biomass production, the leading mode can instead be generally understood and derived mechanistically as the vector of species yields. Because yields are measured in monoculture, this coordinate can be determined before any community is assembled. Moreover, being species traits, yields are largely unaffected by population dynamics. This robustness is notable because ecological and evolutionary dynamics can restrict the community states that are accessible and thereby constrain responses to artificial selection^56,57^, whereas the dominant functional axis identified here remains stable as community composition changes.

Our findings must grapple with several limitations. First, our framework is most directly applicable to fully sampled or densely sampled community-function landscapes, whereas natural microbiomes are usually only sparsely observed, motivating sparse or structure-aware approaches for learning microbiome function^15,17^. Second, we considered binary community composition in the initial inoculum, although natural communities vary continuously in abundance. While we find that presence/absence at the time of inoculation does a good job at describing community composition even for serially passed communities, it is possible that our results will not extend to every community and for functions beyond biomass. More work needs to be done to explore the limits of this finding. On this note, complementary low-dimensional approaches are being developed for abundance data, including methods based on low-variance ecological constraints in microbial communities ^58^ and annotation-free inference of functional groups from patterns of taxonomic covariation ^59^. Third, the biological interpretation of collective modes is likely function-dependent: while the leading biomass mode can be typically linked to yield, modes associated with other functions, such as secondary-metabolite production, may require different species traits or mechanistic models.^46^.

Future work should establish the limits and boundaries of our findings, identifying what ecological mechanisms will create low-versus high-dimensional community-function landscapes. It is plausible that strong niche partitioning, facilitation, or environment-specific metabolic switches may generate multiple competing axes and weaken the dominance of one or two collective modes^17,49^. Consistent with this possibility, replicate communities assembled under similar conditions may converge in core metabolic functions while retaining composition-dependent differences in auxiliary functions^60,61^.

More broadly, our theoretical analyses can be applied to any system in which a function is defined over combinations of binary components, as is the case of genetics where these ideas originated^47,48 62,63^. We hope that our work will motivate the ongoing search for unifying principles between ecological community-function landscapes and genotype–phenotype maps^28,62,64–67^ and stimulate a similar convergence with recent work on evolutionary collective modes^68^, where long-term evolutionary dynamics are organized by collective directions in gene space.

## Acknowledgements

We are especially grateful to Alba Rodriguez Carreño for logistic and administrative assistance at all stages of this project.

## Methods

*Five species–five resources combinatorial assembly experiment* We constructed a fully sampled species-resources community-function landscape using five *Pseudomonas* species and five carbon sources: glycerol, glucose, fructose, citrate and acetate (Supplementary Table T1 and Fig. S1). All 32 presence-absence combinations of species were crossed with all 32 presence-absence combinations of resources, generating 1,024 species-resources combinations. Communities were assembled in M9 minimal medium and grown for 24 h. Biomass was measured as OD_600_, and pyoverdine production was measured from culture supernatants as OD_405_. Combinations with no species, no supplied carbon source, or both, were assigned zero biomass and zero pyoverdine production. The full experiment was performed across four independent biological replicates. Details of strains, media preparation and combinatorial assembly are provided in Supplementary Text S1.

### Serial-dilution experiment

To test whether the dominant productivity mode remains stable through ecological dynamics, we assembled all 31 non-empty communities from a second pool of five independently isolated bacterial species enriched for stable coexistence in citrate-limited M9 medium. Communities were grown with citrate as the sole carbon source and propagated by daily 1:100 serial dilution for ten passages. Biomass was measured every 24 h, generating a complete biomass landscape at each passage. Species-level productivity was estimated from monoculture dynamics and compared with the leading collective mode inferred from each daily landscape. Details of the enrichment, community assembly, serial-passaging protocol and yield estimation are provided in Supplementary Text S1.

### Landscape analysis

Community-function landscapes were represented on a binary hypercube, with species encoded by presence–absence variables. Interaction coefficients were computed using the Walsh-Hadamard transform. Functional Effect Equations were fitted for each focal species by regressing the effect of adding that species against the function of the corresponding background community. Relationships in interaction space were obtained by comparing coefficients involving the focal species with the corresponding lower-order coefficients not containing it. Mathematical definitions of the Walsh-Hadamard coefficients, Functional Effect Equations and interaction-coefficient relationships are provided in Supplementary Texts S2–S3. All landscape analyses were implemented using custom Python routines developed in the `epistasia2.0` package.

### Ecological collective modes

For each landscape, we computed the functional effects of all species across community backgrounds and used them to construct the second-moment matrix of marginal effects. The eigenvectors of this matrix define ecological collective modes, and the associated eigenvalues quantify the spectral weight captured by each mode. As a complementary measure of effective dimensionality, we computed the stable rank of **M**^35^,

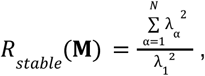

where λ_1_ is the largest eigenvalue of **M**. Values close to 1 indicate that most of the squared spectral weight is concentrated in a single collective mode. Communities were projected onto latent coordinates defined by the leading modes. One- and two-dimensional reconstructions were obtained by projecting the second-order Walsh-Hadamard approximation of each landscape onto these coordinates. Reconstruction accuracy was quantified using root mean squared deviation. Details of the marginal-effect matrix, spectral decomposition and latent-coordinate reconstructions are provided in Supplementary Texts S4–S6.

### Previously published datasets

The same analysis pipeline was applied to previously published fully sampled community-function landscapes spanning different microbial systems, community sizes and measured functions. Dataset descriptions and preprocessing details are provided in Supplementary Text S9 and Supplementary Table T2.

## Code availability

All analyses and figure-generating routines are implemented in the open-source Python package `epistasia2.0`, enabling the computation of FEEs, Walsh-Hadamard interaction spectra, second-moment matrices, collective modes and latent-coordinate projections from community-function landscapes. Codes and tutorials will be publicly available upon publication.

## Data availability

The data will be made publicly available upon official publication of the manuscript.

## Supplementary Information

### S1 Experimental methods

#### S1.1 Strains and media

We used two five-species bacterial pools in this study. The first pool was used for the five species– five resources combinatorial assembly experiment, and the second for the long-term serial-dilution experiment.

- **Five species–five resources experiment:** *Pseudomonas atacamensis* (P1), *Pseudomonas putida* (P2), *Pseudomonas alloputida* (P3), *Pseudomonas aeruginosa* PA14 (PA) [13], and *Pseudomonas putida* KT2440-GFP (KT) [16]. Strains P1, P2 and P3 were isolated from distinct geographical locations in Madrid, Spain.
- **Serial-dilution experiment:** *Acinetobacter johnsonii, Serratia proteomaculans, Pseudomonas defluvii, Aeromonas hydrophila* and *Lelliottia amnigena*. These strains were individually isolated from distinct geographical locations in Salamanca, Spain. To test whether these strains coexist, they were mixed and were serially passaged for seven days until stable coexistence of the five strains was observed.

Genomic DNA from isolates was subjected to whole-genome sequencing (WGS) on an Illumina NovaSeq X platform paired-end, 2×150 bp reads). Raw reads were processed with a custom Nextflow (DSL2) pipeline [6], including quality trimming with fastp [4], de novo assembly with SPAdes [1], and assembly quality control with QUAST [12] and CheckM2 [5]. Draft genomes were annotated using Bakta v1.12.0 with the full database (v6.0) [21], and taxonomically classified using the classifier module of PATO, a pangenome analysis toolkit [10]. Strains P3, PA and KT were routinely grown at 37^°^C, whereas the remaining strains were incubated at 28^°^C, unless otherwise stated. Cell stocks were stored at −80^°^C in 40% glycerol.

Minimal growth medium consisted of M9 salts supplemented with MgSO_4_ and CaCl_2_. Resource mixtures were built from five carbon sources: glucose, fructose, glycerol, citrate and acetate (Table T1). Carbon-source stock solutions were prepared at 5× concentration, sterilized using 0.22 *µ*m filters, and diluted during assembly so that each supplied carbon source reached a final concentration of 0.014 Cmol L^−1^.

#### S1.2 Five species – Five resources combinatorial assembly experiment

The five *Pseudomonas* species were assembled in all possible presence–absence combinations and crossed with all possible presence–absence combinations of five carbon sources, generating a complete 2^5^ × 2^5^ species–resource design (Fig. S1).

- **Starting inocula:** each strain was streaked from a glycerol stock onto LB agar and grown overnight at their respective strain-specific temperatures. A single colony of each species was inoculated into a 50 mL conical tube containing 25 mL of LB broth (Condalab) and grown overnight under static conditions at 32^°^C. Cultures were harvested by centrifugation at 2,500 rpm (1,215× *g*; Thermo Scientific SL Plus Centrifuge series) for 25 min. Cultures were washed three times in carbon-free M9 medium and adjusted to optical density at 600 nm (OD_600_) = 0.01 using a Multiskan SkyHigh microplate reader (Thermo Fisher Scientific).
- **Species combinations:** all 2^5^ = 32 species combinations were prepared following the previously described protocol by Diaz-Colunga et al. [8]. Whenever a species was present, its contribution was adjusted to OD_600_ = 0.002.
- **Resource combinations:** all 2^5^ = 32 carbon-source combinations were prepared in 96-deep-well plates. The total carbon concentration ranged from 0.014 to 0.07 Cmol L^−1^, depending on the number of supplied resources.
- **Assembly:** each species combination was crossed with a full carbon-source landscape. A volume of 200 *µ*L of bacterial inoculum was added to 200 *µ*L of resource medium, yielding a final volume of 400 *µ*L. This 1 :1 dilution brought the contribution of each species present down to an initial OD600 = 0.001 per well.
- **Replication and measurements:** the full landscape was assayed across four independent biological replicates. After 24 h of incubation at 32^°^C, cultures were homogenized, biomass was quantified as OD_600_. Plates were then centrifuged and pyoverdine production was quantified from the supernatant as OD_405_.

#### S1.3 Long-term stability combinatorial assembly experiment

The serial-dilution experiment was designed to test whether the leading productivity mode remained stable as community composition changed through population dynamics. We used five species that had previously been enriched under daily serial passaging in citrate-limited M9 medium.

- **Species pool:** *Acinetobacter johnsonii, Serratia proteomaculans, Pseudomonas defluvii, Aeromonas hydrophila* and *Lelliottia amnigena*.
- **Inoculum preparation:** strains were streaked from glycerol stocks onto ESBL Chromogenic Medium agar plates (Condalab) and incubated overnight at 28^°^C. Single colonies were grown overnight in LB broth under static conditions, washed three times in carbon-free M9, and resuspended in citrate-supplemented M9 medium.
- **Community assembly:** all 2^5^ = 32 species combinations were assembled following the protocol described by [8]. Whenever a species was present, its initial contribution was OD_600_ = 0.001, independent of community composition.
- **Growth and dilution:** communities were grown in M9 supplemented with citrate (0.07 mol C L^−1^). Every 24 h, cultures were homogenized, biomass was measured as OD_600_, and 3.75 *µ*L of culture were transferred into 371.25 *µ*L of fresh medium, corresponding to a 1:100 dilution.
- **Duration and replicability:** the experiment was continued for 10 daily passages, across two independent biological replicates, each comprising three technical replicates. A schematic representation of the experiment is shown in Fig. 4 of the main text.

The final community composition after 10 daily passages was assessed by plating the 31 assembled communities. Species loss was rare: only two final communities were missing one species, as indicated in Fig. S18.

##### Yield estimation

The yield of species *i* is defined as the amount of biomass produced per unit of initially available resource, and is estimated here from the net biomass accumulated during one daily growth cycle. The yield can be estimated by comparing the biomass growth of two consecutive days:

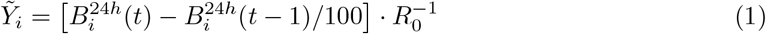

where *B*_*i*_ is the biomass of species *i* in monoculture, *R*_0_ is the initial resource concentration, and the factor 1*/*100 accounts for the daily dilution. This estimation is used for Fig. 4D-F of the main text.

For single-batch experiments, the yield has been estimated as 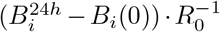, being *B*_*i*_(0) the initial biomass at inoculum. This estimation is used in Fig. 3C of the main text.

### S2 From community-function landscapes to interaction space

We consider ecological systems composed of *N* binary microbial species. A community configuration is encoded as a binary vector **x** = (*x*_1_, …, *x*_*N*_ ), where *x*_*i*_ = +1 denotes presence of species *i* and *x*_*i*_ = −1 its absence. The space of possible communities is therefore *X* = {−1, +1}^*N*^, the vertex set of an *N* -dimensional hypercube. Each configuration is assigned a quantitative community-level outcome, such as total biomass, community growth rate, or ecosystem productivity. This defines a *community-function landscape*, a mapping *F* : *X* →ℝ assigning a functional value to each combination of species.

#### S2.1 Context-dependent interactions as deviations from additivity

We define interactions operationally as deviations from additivity in finite differences of the community-function landscape. For a focal species *i*, its local first-order, or marginal, effect in a given back-ground configuration is

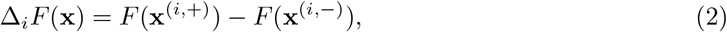

where **x**^(*i*,+)^ and **x**^(*i*,−)^ denote the configurations obtained by setting *x*_*i*_ = +1 and *x*_*i*_ = −1, respectively, while keeping all other coordinates fixed. Thus, Δ_*i*_*F* (**x**) depends only on the non-focal coordinates, which define the background context.

If species acted additively, the marginal effect of *i* would not depend on the state of another species *j*. The local second-order, or pairwise, interaction between *i* and *j* is therefore the corresponding difference of marginal effects

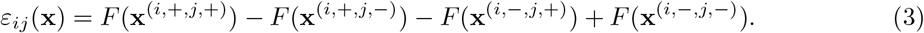

Higher-order local interactions are defined analogously as alternating finite differences over a set of *k* focal species [3]. We denote them by *ε*_**s**_(**x**), where **s** ∈ {0, 1}^*N*^ indicates the subset of focal species. This quantity measures, in the specified background context, the deviation of the joint effect of those species from all lower-order contributions.

#### S2.2 Walsh–Hadamard coefficients as background-averaged interactions

On the discrete hypercube X = {−1, +1}^*N*^, we use the uniform inner product

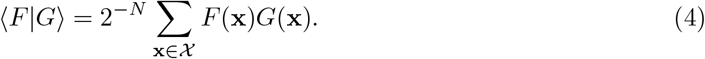

Under this inner product, the Walsh–Hadamard (WH) functions form an orthogonal basis for functions *F* : *X* → ℝ. Each basis function is indexed by a binary vector **s** ∈ {0, 1}^*N*^ :

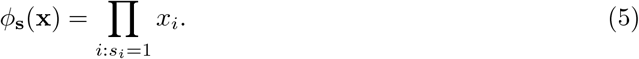

Here **x** denotes a community configuration, whereas **s** selects a subset of species. The WH expansion of the landscape is

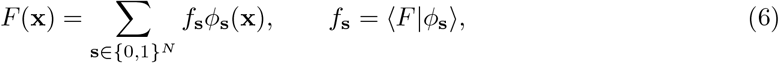

where the WH coefficients *f*_**s**_ correspond to background-averaged local interactions. For an index **s** of order 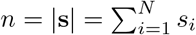, we have

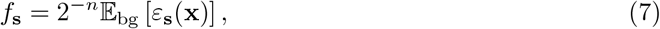

where the average is taken over the non-focal coordinates, with *x*_*i*_ = −1 for all indices with *s*_*i*_ = 1 before applying the finite-difference operator. Thus, after the normalization 2^−*n*^, first-order coefficients encode mean marginal effects, whereas higher-order coefficients encode background-averaged deviations from lower-order additivity.

#### S2.3 Variance decomposition across interaction orders

We define Var(*F* ) = ⟨*F* ^2^⟩ − ⟨*F* ⟩^2^, with averages taken uniformly over *X*. Since *f*_**0**_ = ⟨*F* ⟩, orthogonality of the WH basis {*ϕ*_**s**_(**x**)} implies Parseval’s identity,

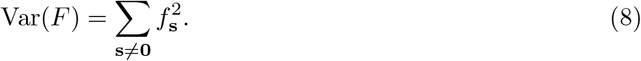

Grouping WH coefficients by order *n* = |**s**| yields the exact order-wise decomposition [3, 22]

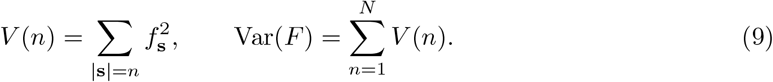

This orthogonal partition holds only when *F* is defined on the full hypercube and the inner product is uniform. For incomplete landscapes, WH basis functions are no longer orthogonal under the empirical inner product on the sampled configurations, so *V* (*n*) no longer defines an exact variance contribution (see [3] for details).

We used this decomposition to quantify how variance is distributed across interaction orders in the empirical biomass and pyoverdine landscapes. Across resource environments, the first two Walsh–Hadamard orders captured the dominant share of variance, typically accounting for approximately 70%–90% of biomass variance and 85%–95% of pyoverdine variance. This motivated the second-order approximation used in the spectral and latent-coordinate analyses (Figs. S2 and S3).

### S3 Functional Effect Equations constrain interaction coefficients

Functional Effect Equations (FEEs) relate the effect of adding species *i* to the function of the background community through simple linear relationships [9],

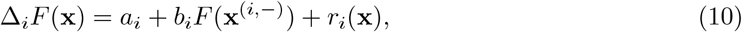

where Δ_*i*_*F* is the unnormalised finite difference defined in Eq. (2), and *r*_*i*_(**x**) captures deviations from linearity. We first consider the exact limit *r*_*i*_(**x**) = 0. In this limit, FEEs reduce the effective degrees of freedom of the community-function landscape, suggesting an underlying low-dimensional structure.

To make this constraint explicit, we express the landscape in the WH basis, Eq. (6). The unnormalised finite-difference operator acts on WH basis functions as

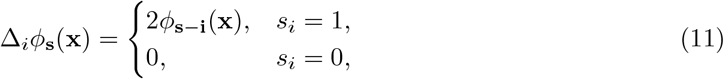

where **i** is the unit vector in direction *i*. Therefore,

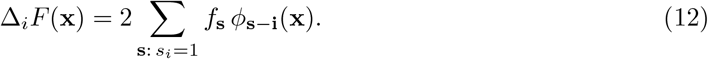

Similarly, the background function entering the FEE can be expanded on the reduced subcube *x*_*i*_ = −1 as

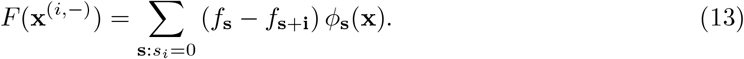

For an exact FEE, Eq. (10) defines a linear relationship between two functions expanded in the same WH basis on the reduced subcube. Matching coefficients gives, for all **s** with *s*_*i*_ = 0,

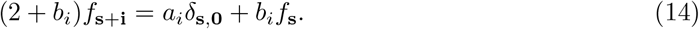

Thus, for non-constant terms |**s**| ≥ 1,

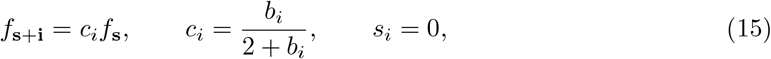

whereas the constant term fixes the first-order coefficient,

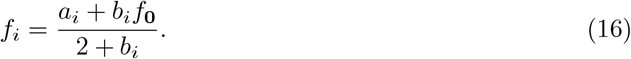

For compactness, we write *f*_*i*_, *f*_*ij*_, and more generally *f*_*S*_ for the WH coefficient whose index vector **s** has entries equal to one for components in {*i*}, {*i, j*}, or *S*, respectively, and zero otherwise. Equation (15) then reads

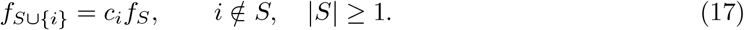

If exact FEEs hold for all focal species on the same landscape, then, for example,

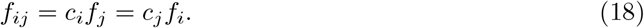

This symmetry imposes a consistency condition on the slopes and first-order coefficients. When these consistency conditions hold for every species *i* and the relevant first-order coefficients are nonzero, the ratios *f*_*i*_*/c*_*i*_ are equal across species. Denoting this common ratio by *κ* repeated application of Eq. (17) gives

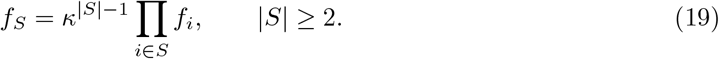

Thus, exact and mutually consistent FEEs imply that WH coefficients across orders are linked by species-specific multiplicative factors. More generally, the residual can be expanded in the WH basis of the reduced subcube as

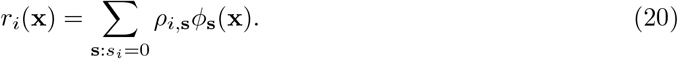

Matching coefficients in Eq. (10) then gives

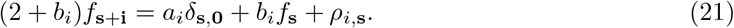

Therefore, deviations from the exact cross-order recursion are directly determined by the WH coefficients of the FEE residual. Empirical analyses across resource environments are shown for biomass in Figs. S5–S6, and for pyoverdine in Figs. S7–S8. Consistent with Eq. (21), more predictive FEEs are associated with stronger cross-order relationships among WH coefficients (Fig. S9). These relationships are generally noisier than the corresponding FEEs, as expected from the increasing uncertainty of higher-order WH coefficients [3].

### S4 Ecological collective modes and low-dimensional community-function landscapes

FEEs constrain WH coefficients across orders, Eqs. (18)-(19). We now express these constraints at the landscape level by collecting all marginal effects into a discrete gradient field and studying the second-moment matrix of this field. Its dominant eigendirections define *ecological collective modes*.

#### S4.1 Discrete gradients

To study all marginal effects simultaneously, we introduce a discrete gradient field. We define the normalised discrete derivative along coordinate *i* as

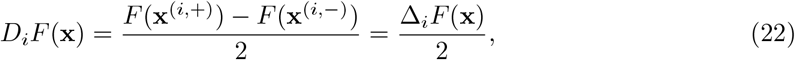

where the factor 1*/*2 accounts for the spin displacement Δ*x*_*i*_ = 2. This differs from the un-normalised finite difference Δ_*i*_*F* used so far. This convention keeps the local finite differences consistent with previous literature, while using the normalised derivative only for the gradient representation. The discrete gradient field is

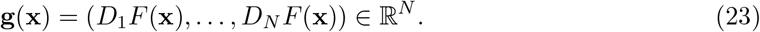

It collects the normalised local marginal effects of all species and plays the role of a gradient field on the hypercube. The normalisation removes the factor of 2 in the finite-difference action on WH basis functions, Eq. (11):

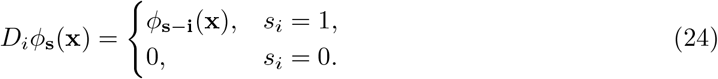

Therefore,

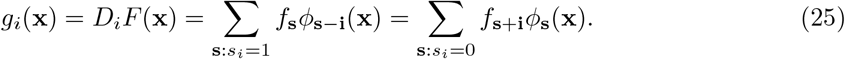

Averaging over configurations and using WH orthogonality gives

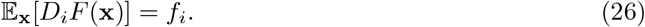

Thus, the background-averaged gradient equals the vector of first-order WH coefficients,

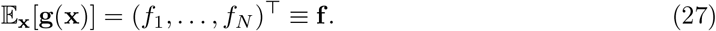

#### S4.2 Second-moment matrix of marginal effects

We now introduce the central geometric object of this work,

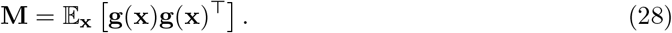

The matrix **M** measures the second moments and alignments of marginal effects across backgrounds. Let *G* be the matrix whose rows are configurations and whose columns are marginal effects,

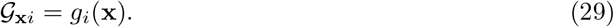

Then

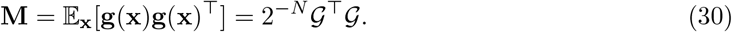

Thus, diagonalizing **M** is equivalent to performing an uncentred PCA, or equivalently an SVD, of the marginal-effect matrix *G*. Because **M** is symmetric and positive semidefinite, it admits the spectral decomposition

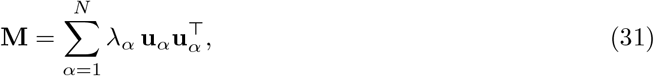

where {**u**_*α*_} form an orthonormal basis and *λ*_1_ ≥ *λ*_2_ ≥ · · · ≥ *λ*_*N*_ ≥ 0. Each eigenvector **u**_*α*_ defines a collective direction in species space along which marginal effects are coordinated, corresponding to an *ecological collective mode*. The associated eigenvalue *λ*_*α*_ quantifies the mean-square magnitude of the marginal-effect field along that direction. Using the WH expansion of *g*_*i*_, Eq. (25), we obtain, for *i*≠ *j*,

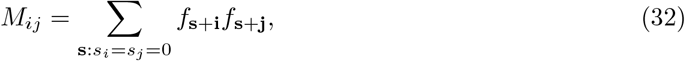

and

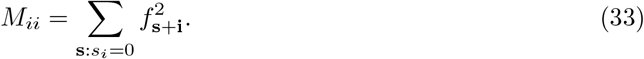

Equivalently, **M** can be decomposed by WH order,

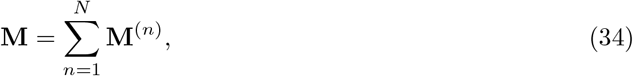

where **M**^(*n*)^ contains the contribution of order-*n* WH coefficients. Its entries are

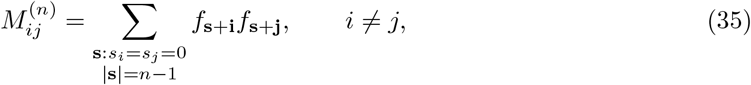

and

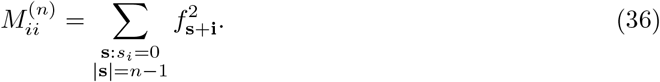

Thus, although **M** is built from first-order derivatives, it contains contributions from interactions of all orders. In components,

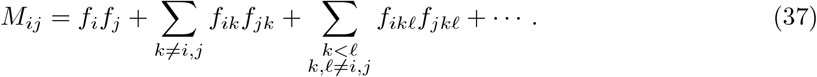

Taking the trace gives

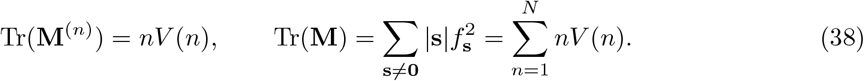

The total spectral weight of **M** is therefore the first moment of the variance spectrum. The empirical spectra of **M** across resource environments are shown in Fig. S14.

#### S4.3 Discrete curvature and pairwise interactions

For species *i*≠ *j*, we define the normalised second discrete derivative

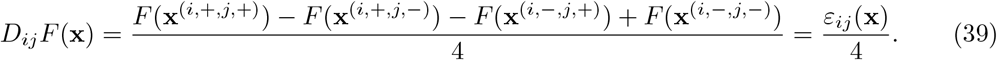

This quantity measures how the normalised marginal effect of *i* depends on the state of *j*. It vanishes for additive landscapes and provides a discrete notion of curvature on elementary faces of the hypercube. We define the global Hessian matrix as the background-averaged discrete curvature,

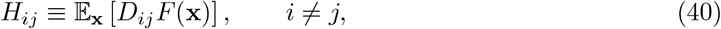

and set *H*_*ii*_ = 0. Using the WH expansion, second discrete derivatives act on WH basis functions as

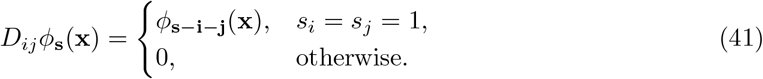

It follows that

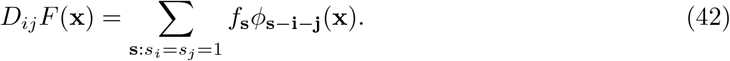

Averaging over configurations and using WH orthogonality gives

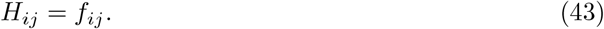

Thus, the global Hessian matrix coincides exactly with the matrix of second-order WH coefficients.

#### S4.4 Second-order WH truncation

The WH expansion Eq. (6) can be grouped by interaction order as

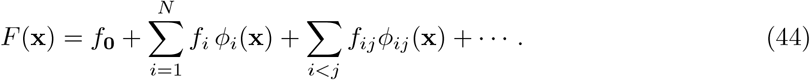

In community-function landscapes studied to date, the variance spectrum is typically dominated by first- and second-order terms, *V* (1) + *V* (2) ≫ _*n*≥3_ *V* (*n*), so the landscape can be approximated by retaining only additive and pairwise WH coefficients [3, 22]. Since *ϕ*_*i*_(**x**) = *x*_*i*_ and *ϕ*_*ij*_(**x**) = *x*_*i*_*x*_*j*_, the second-order truncation is

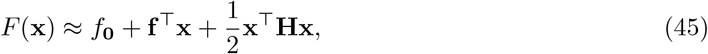

where **f** = (*f*_1_, …, *f*_*N*_ )^⊤^ and *H*_*ij*_ = *f*_*ij*_ for *i* ≠ *j*, with *H*_*ii*_ = 0. Thus, the landscape is approximated by a quadratic function on the hypercube. The vector **f** gives the background-averaged slopes, whereas **H** gives the background-averaged pairwise curvature.

#### S4.5 Closed-form expressions in the quadratic WH model

When the WH spectrum is dominated by first- and second-order coefficients, the landscape can be approximated by its second-order WH truncation. The gradient field then has the form

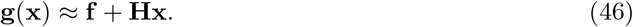

Equivalently, for each species *i*,

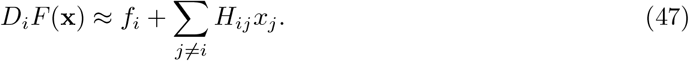

Taking the mean and variance across configurations, and using E[**x**] = **0** and E[**xx**^⊤^] = **I**, gives

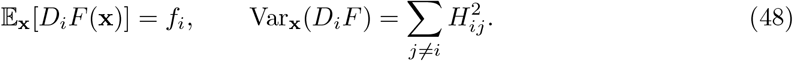

Thus, first-order coefficients set the mean marginal effects, whereas pairwise curvature terms determine their variability across backgrounds. The corresponding second-moment matrix is

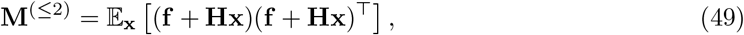

which yields

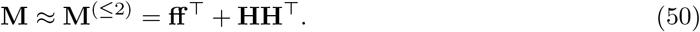

The term **ff** ^⊤^ is the contribution of the mean gradient, whereas **HH**^⊤^ is the covariance of the gradient field induced by pairwise curvature.

### S5 Rank–1 limit: single ecological collective mode

We now examine the idealized case in which **M** is exactly rank–1,

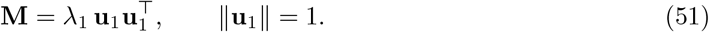

Geometrically, this means that all marginal-effect vectors are aligned with a single collective direction. Indeed, for any vector **v** orthogonal to **u**_1_,

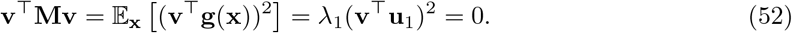

Since the integrand is nonnegative and the uniform measure assigns positive weight to every configuration, it follows that

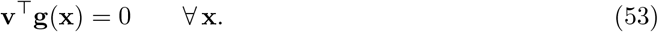

Thus, every gradient vector lies in the one-dimensional subspace spanned by **u**_1_. Equivalently, there exists a scalar field *α*(**x**) such that

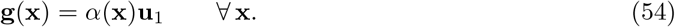

Since ∥**u**_1_∥ = 1, this scalar field is the projection of the gradient onto the collective mode,

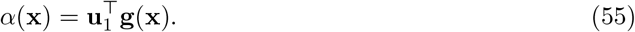

Its second moment is fixed by the nonzero eigenvalue:

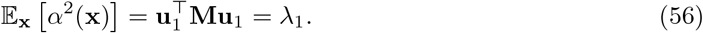

In the quadratic WH model,

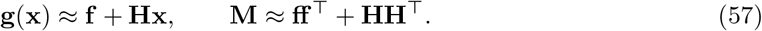

Exact rank–1 structure of **M** implies

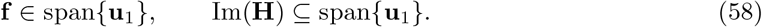

Thus, there exists a scalar *B* such that

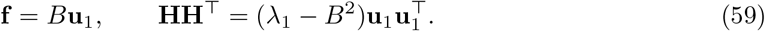

Here *B* is the additive projection along the leading mode, whereas *λ*_1_ − *B*^2^ is the spectral weight of the background-dependent contribution to the gradient field. If **H** were an unconstrained symmetric matrix, Im(**H**) ⊆ span{**u**_1_} would imply 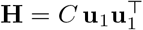, with *C*^2^ = *λ*_1_ −*B*^2^. However, in the WH representation the pairwise matrix has zero diagonal, *H*_*ii*_ = 0. Therefore, a nonzero symmetric rank–1 **H** is incompatible with this convention, since 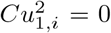 for all *i* forces *C* = 0 unless the mode is trivial. Hence, in a strict quadratic WH landscape, exact rank–1 **M** is only possible for purely additive landscapes, or as an idealized limiting case.

#### S5.1 Approximate rank–1 regime

When *λ*_1_ ≫ *λ*_2_, …, *λ*_*N*_, the gradient field is predominantly aligned with the leading collective mode **u**_1_. We decompose it as

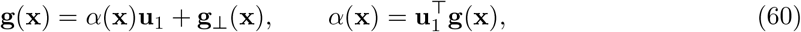

where **g**_⊥_(**x**) is orthogonal to **u**_1_. The spectral identity gives

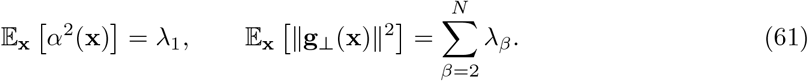

Thus, when 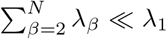, most of the mean-square gradient magnitude is captured by a single collective direction. This motivates projecting configurations onto

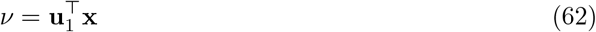

and seeking an approximate functional collapse of the form *F* (**x**) ≈ *q*(*ν*). Because eigenvectors are defined up to an overall sign, we oriented **u**_1_ so that *F* (**x**) varied with 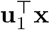 in the same direction as with species biodiversity.

#### S5.2 Functional collapse and latent projection

The approximate rank–1 regime motivates a one-dimensional representation of the landscape in terms of the collective coordinate *ν* defined in Eq. (62). In the quadratic WH approximation, we consider a functional collapse of the form

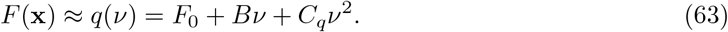

This corresponds to additive and pairwise coefficients organized around the same collective direction. Indeed,

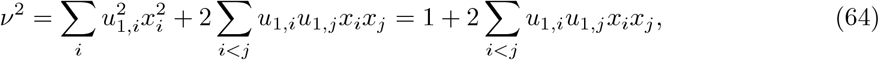

where ∥**u**_1_∥ = 1 and 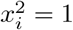. The diagonal contribution is constant on the spin hypercube and can be absorbed into the intercept, so

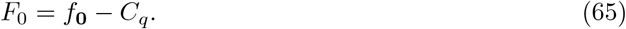

Thus, the zero diagonal of the WH pairwise matrix is compatible with a quadratic functional collapse; it only removes a constant contribution to *ν*^2^. For empirical landscapes, the linear coefficient along the collective coordinate is obtained by projection,

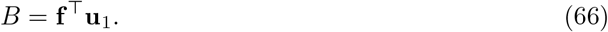

The total second moment of the gradient field along the same direction is

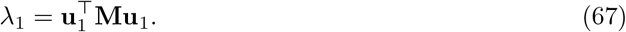

We define the quadratic coefficient used in the one-dimensional spectral prediction as

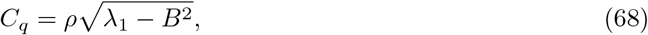

where the sign *ρ* = ±1 is chosen according to the observed orientation of the projected response. Thus, *C*_*q*_ is fixed in magnitude by the spectral weight of the leading mode and parameterizes the predicted response *q*(*ν*). It should be distinguished from *C*_*H*_, the coefficient of a specific aligned pairwise WH matrix.

At the coefficient level, a literal aligned pairwise WH structure can be written as

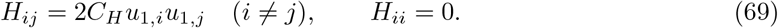

For such a structure,

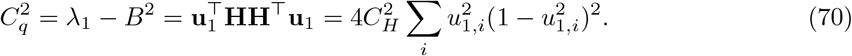

Therefore, the spectral coefficient *C*_*q*_ used in the projected response and the WH coefficient *C*_*H*_ are related, but not identical in general, because the pairwise WH matrix has zero diagonal. One-dimensional latent-coordinate projections for biomass and pyoverdine landscapes across resource environments are shown in Figs. S10 and S11.

#### S5.3 Gradient covariance structure

The exact rank–1 analysis above shows that low rank of **M** reflects alignment of the full gradient field, but does not by itself distinguish additive from background-dependent structure. To separate these contributions, we define the centered second moment of the gradient field,

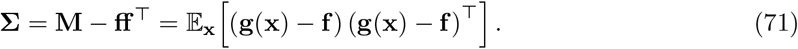

This matrix captures the background-dependent variability of marginal effects. For a purely additive landscape, *F* (**x**) = **a**^⊤^**x**, the gradient is constant, **g**(**x**) = **a**. Therefore,

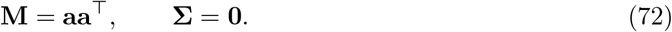

Thus, **M** can be rank–1 even in the absence of background-dependent interactions. In contrast, **Σ** vanishes for additive landscapes and isolates the component of gradient variation induced by interactions.

The two matrices therefore encode complementary notions of low dimensionality. Low rank of **M** indicates that the full marginal-effect field is aligned with a small number of collective directions. Low rank of **Σ** indicates that the deviations from the mean gradient are low-dimensional. A nontrivial one-dimensional functional collapse requires both the mean gradient and the background-dependent variability to be organized along the same collective coordinate. The corresponding covariance spectra across resource environments are shown in Fig. S15.

### S6 Multi-dimensional latent manifolds

When the spectrum of **M** is concentrated but not strictly rank–1, the alignment of marginal effects extends to a *K*-dimensional dominant subspace. Let 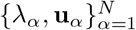 denote the eigenpairs of **M**, ordered by decreasing eigenvalue, and let **U** = (**u**_1_, …, **u**_*K*_) collect the leading eigenvectors. We define the latent coordinates as the orthogonal projections

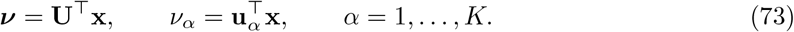

We then approximate the landscape by a quadratic function on this latent subspace,

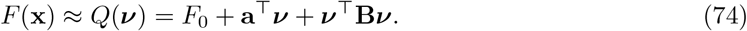

The projected coefficients are defined as

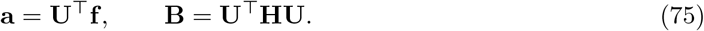

Thus, additive effects and pairwise curvature are reorganized in terms of collective coordinates rather than individual species. Because the configurations are uniformly distributed on the spin hypercube, E_**x**_[***ν***] = **0** and

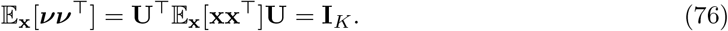

Therefore, the intercept is chosen so that the projected landscape has the same mean as the WH landscape:

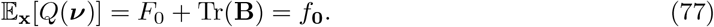

Hence

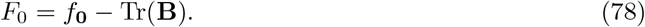

The matrix **B** encodes curvature within the latent manifold: diagonal entries describe curvature along individual collective coordinates, whereas off-diagonal entries describe coupling between collective coordinates. Thus, the relevant interaction structure of the landscape is reorganized from taxon/resource-level interactions to interactions among collective ecological modes. The spectral weight captured by the first *K* modes is

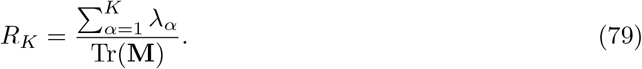

Choosing *K* such that *R*_*K*_ exceeds a prescribed threshold yields a low-dimensional geometric representation of the landscape. Projecting the quadratic expression for **M** onto an eigenvector gives

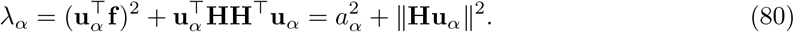

Expanding the second term in the complete eigenbasis of **M** gives

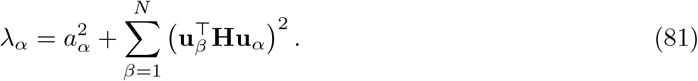

If the dominant subspace captures most of the curvature, this relation is approximated by

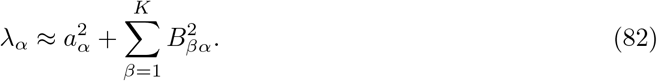

Thus, the variability captured by each collective direction is partitioned between the projected mean gradient and curvature within the latent subspace. For *K* = 1, this construction reduces to a one-dimensional projected quadratic collapse,

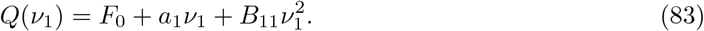

Note that *Q*(*ν*_1_, *ν*_2_ = 0) is a slice of the two-dimensional projected manifold and is not, in general, identical to the one-dimensional response *q*(*ν*_1_). More generally, for *K >* 1, the landscape is represented as a quadratic latent manifold embedded in the original component space. Directions associated with large eigenvalues define stiff collective modes of community function, whereas directions associated with small eigenvalues correspond to weakly constrained, sloppy modes. Two-dimensional latent manifolds for biomass and pyoverdine landscapes are shown in Figs. S12 and S13.

### S7 Unifying global epistasis

The term *global epistasis* is used to describe related but distinct forms of structured background dependence. In evolutionary genetics, global epistasis often denotes systematic relationships between mutational effects and background fitness, such as diminishing returns or increasing costs.

These relationships can be formulated as regression-based models in which the effect of a mutation depends approximately linearly on the fitness of its genetic background [19]. The FEEs used above are the ecological analogue of this formulation for community-function landscapes [7, 9].

A second formulation arises in biophysics and genotype–phenotype mapping, where the landscape is described as a nonlinear function of an additive latent coordinate,

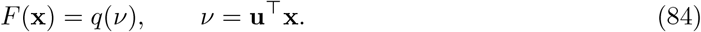

In this representation, species combine additively in the latent variable, while nonlinear background dependence is induced by the map *q* [18, 14]. The framework developed in this work connects these two perspectives through the geometry of the marginal-effect field. The central object is

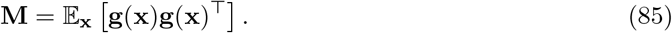

Spectral concentration of **M** implies that marginal effects vary primarily along one, or a few, collective directions. These directions define latent coordinates for functional collapse and, at the same time, constrain how marginal effects vary across backgrounds.

This connection can be made at different levels of approximation. The exact FEE constraint derived in Section S3 is an all-order statement about WH coefficients, Eq. (17). It is therefore a constraint on background-averaged interaction coefficients, rather than on the full distribution of local microscopic effects [19]. To connect this all-order recursion with the interaction-based interpretation used in previous work, and with the empirical dominance of low-order variance in microbial landscapes, we first specialize to the quadratic WH approximation. In this lowest-order limit, FEE predictability, factorized pairwise interactions, latent-coordinate nonlinearities, and spectral concentration of **M** become different projections of the same underlying organization.

#### S7.1 Low-order interpretation of FEE predictability

Section S3 showed that an exact FEE, imposed as an identity on the reduced subcube, recursively constrains WH coefficients at all orders,

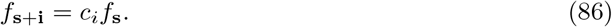

We now connect this all-order identity with the empirical regression diagnostics used to quantify global epistasis. Previous work derived low-order approximations for the variance of the background function, the variance of the focal effect, and their covariance, leading to approximate expressions for the FEE slope and the coefficient of determination 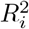 in terms of average effects and average pairwise interactions [9]. Here we rewrite the corresponding 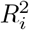 approximation in WH notation and show that it has a simple geometric interpretation as an alignment condition.

The coefficient of determination for the FEE of component *i* is

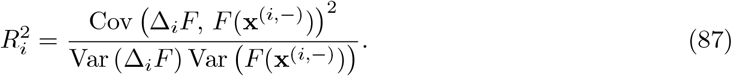

Keeping only the leading terms in the low-order expansion, the background-dependent part of Δ_*i*_*F* is determined by the pairwise WH coefficients *f*_*ij*_, whereas the leading variation of the background function on the reduced subcube is determined by *f*_*j*_ − *f*_*ij*_. This gives

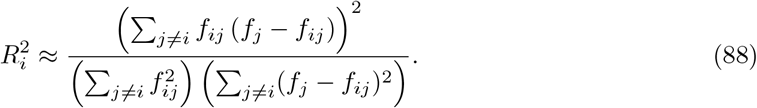

Defining

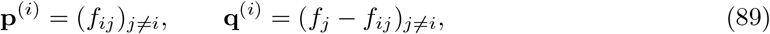

Eq. (88) becomes

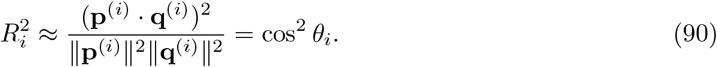

Thus, 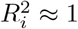 means that the vector of pairwise WH coefficients involving component *i* is approximately collinear with the linear component of the background function on the reduced subcube. Equivalently, there exists a scalar *α*_*i*_ such that

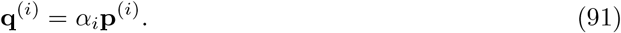

In components,

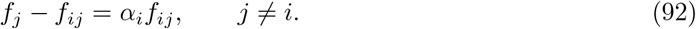

Solving for *f*_*ij*_ gives

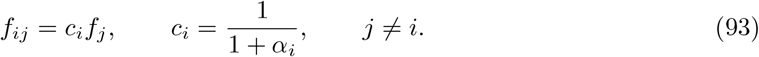

Therefore, perfect FEE predictability at leading order implies that the row of pairwise WH coefficients associated with component *i* is proportional to the first-order WH profile of the remaining components. This is the lowest-order, regression-based counterpart of the all-order recursive constraint in Eq. (15).

Conversely, if the pairwise WH coefficients factorize as

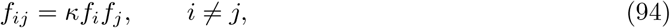

with *κf*_*i*_ ≠ 0. For a fixed component *i*, the vectors entering Eq. (90) become

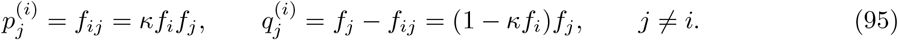

Hence

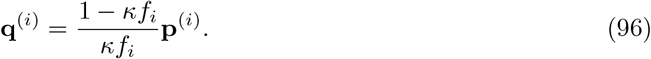

The two vectors are therefore collinear, and Eq. (90) gives

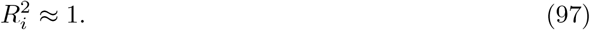

Thus, factorized pairwise WH coefficients are sufficient to generate perfect FEE predictability at leading order. Together with the previous result, this shows that high FEE predictability and factorized pairwise interaction profiles are equivalent low-order descriptions of the same structure.

#### S7.2 From per-component predictability to spectral concentration

We now impose Eq. (93) simultaneously for all species and examine the resulting consistency constraints. Assume that, for every component *i*, there exists a scalar *c*_*i*_ such that

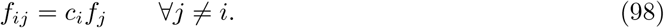

Using symmetry of pairwise coefficients, *f*_*ij*_ = *f*_*ji*_, we obtain

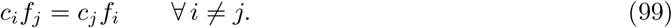

In the non-degenerate case *f*_*i*_ ≠ 0, this implies

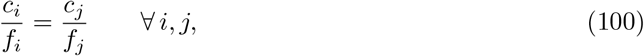

and therefore there exists a global scalar *κ* such that

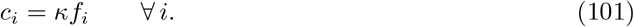

Substituting back gives the factorized pairwise structure

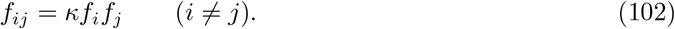

Thus, simultaneous FEE predictability at leading order constrains the off-diagonal pairwise coefficients to be organized by the additive profile **f** .

Within the quadratic WH approximation, this structure gives

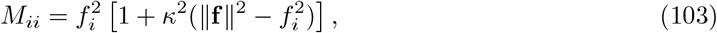

and, for *i* ≠ *j*,

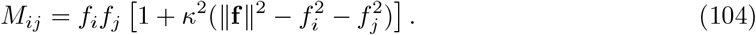

The leading coherent contribution to **M** is therefore organized by the outer product **ff** ^⊤^. If the additive effects are distributed over many species, so that no single 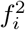 dominates ∥**f** ∥^2^, the diagonal-exclusion corrections are small and

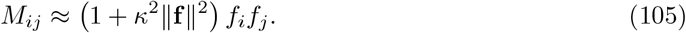

In this regime,

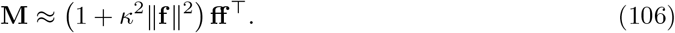

Thus, the coherent part of **M** is approximately rank–1, with leading eigenvector aligned with **f** . In the strict quadratic WH representation, exact rank–1 structure with nonzero pairwise curvature is generally obstructed by the zero diagonal of **H**. The factorized interaction structure therefore induces spectral concentration, rather than exact rank–1 degeneracy. In this sense, global epistasis appears geometrically as a low-dimensional organization of marginal effects.

In empirical landscapes, 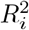 is typically high but not exactly one. Then the proportionality condition in Eq. (93) holds only approximately, and the pairwise coefficients can be written as

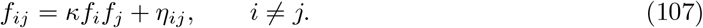

When the residual structure *η*_*ij*_ is small or itself low-dimensional, the spectrum of **M** remains concentrated along one or a few collective directions. Thus, empirical global epistasis corresponds to spectral concentration of the marginal-effect field rather than exact rank–1 structure.

#### S7.3 Connection to latent-coordinate models

Classical latent-coordinate models of global epistasis write the observed function as a nonlinear transformation of an additive latent trait,

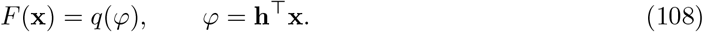

In such models, the latent direction **h** is either inferred statistically from genotype–phenotype data [18] or associated with a collective physical mode of the system [14]. The framework developed here provides a geometric interpretation of this construction in terms of the marginal-effect field.

In Section S5.2, we showed that when the spectrum of **M** is concentrated around a single dominant mode, configurations can be projected onto 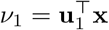, where **u**_1_ is the leading eigenvector of the second-moment matrix of marginal effects. In the second-order WH approximation, this gives the quadratic collapse in Eq. (63). Thus, the additive latent trait of classical global-epistasis models is identified with the leading collective coordinate of the marginal-effect field.

This interpretation extends naturally to multiple latent coordinates, as described in Section S6. When higher-order WH coefficients are also organized along the same dominant coordinate, the projected response is no longer restricted to a quadratic polynomial, but becomes a more general nonlinear function *q*(*ν*_1_).

#### S7.4 Collective epistasis as interaction-induced variability

We have focused on **M** because its spectral concentration constrains the full marginal-effect field. This is the structure that connects the regression-based view of global epistasis, through FEEs, with the latent-coordinate view, *F* (**x**) ≃ *q*(*ν*). If marginal effects are aligned along one, or a few, collective directions, then they can be predicted from low-dimensional background information and the landscape admits a low-dimensional collapse.

However, as discussed in Section S5.3, **M** does not distinguish whether this organization is generated by interactions or by purely additive effects. A purely additive landscape has perfectly aligned, but background-independent, marginal effects. To isolate the interaction-induced component of marginal-effect variation, we use the centered covariance matrix **Σ**.

We refer to low-dimensional structure in **Σ** as *collective epistasis*: interactions generate context dependence, but this context dependence is itself coordinated along a small number of collective directions. Thus, **M** captures the global organization underlying FEEs and latent-coordinate collapse, whereas **Σ** isolates the background-dependent component of that organization.

### S8 Mechanistic origin of global epistasis in consumer–resource systems

The geometric framework above applies to any community function *F* . In this section, we specialize to the case where *F* is total biomass. Our goal is to identify a mechanistic origin for the leading ecological collective mode **u**_1_ for this function (or any other that is strongly correlated with it), and to determine when marginal effects align across configurations, giving rise to a low-rank structure of **M**.

As a proxy to our experiments, we model biomass production through Consumer–Resource models (CRMs), these providing a minimal setting in which collective modes can acquire a trait-based interpretation. In their simplest form, the models assume a single public resource accessed by all species. Multiple resources are considered in Sec. S8.1.3.

For any given community **x** ∈ {−1, 1}^*S*^, species biomasses ({*N*_*i*_}_*i*=1,…,*S*_) and resource abundance (*R*) obey

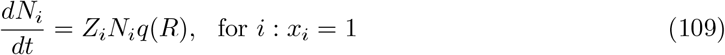

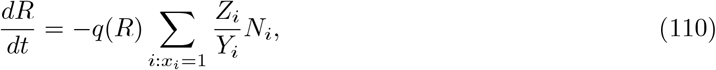

where *Z*_*i*_ and *Y*_*i*_ are the (maximum) per-capita growth rate and yield of species *i*, respectively, and *q* : ℝ → [0, 1] is a function modulating the per-capita growth rates based on the currently available resource, thereby specifying a particular CRM. We can nonetheless make *q* disappear from Eqs. (109)-(110) through the reparameterization *t* ↦ *qt* ≡ *τ* (**x**) of the time parameter, which means all CRMs above have the same formal solution, independently of the functional form of *q*. Equations (109)-(110) then admit the general solution

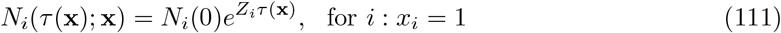

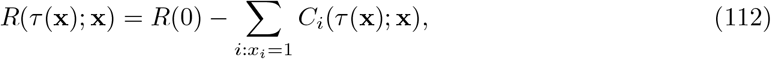

where

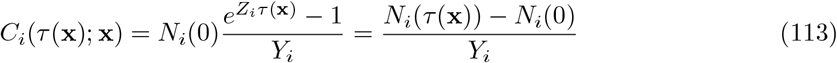

is the total amount of resource *i* consumed up to the effective time *τ* (**x**). Notice the latter is connected to physical time *t* through 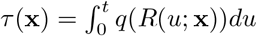.

#### S8.1 Biomass landscape

We define the community function as the total final biomass,

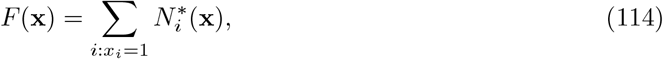

where 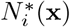 is the biomass species *i* reached at the end of the growth cycle. We henceforth assume the cycle is stopped when a fixed amount *R*^∗^ of resource has been depleted (potentially all of it, *R*^∗^ = *R*(0)) and that species have equal inocula, *N*_*i*_(0) = *n*_0_. If *C*_*i*_(**x**) is the total resource consumed by species *i*, then 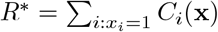.

Let us now build the second-moment matrix of functional effects, **M** = ⟨**Δ***F* **Δ***F*⟩^⊤^ _**x**_ (notice this is just 4 times the matrix ⟨**gg**^⊤^⟩ _**x**_ considered elsewhere in this work). The local effect Δ_*i*_*F* (**x**) reads

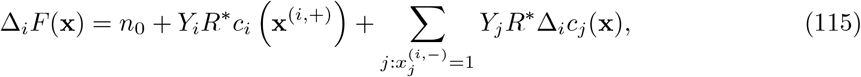

where we introduced the fraction of resource depleted by species *i, c*_*i*_ = *C*_*i*_*/R*^∗^, and the shorthand Δ_*i*_ *c*_*j*_ (**x**) = *c* **x**^(*i*,+)^ *c* **x**^(*i*,−)^ . Isolating the average yield from *Y*, as 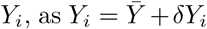 (in vectorial form, 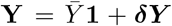, and using the constraint 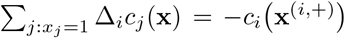 (from *R*^∗^ being fixed), we can recast Eq. (115) into

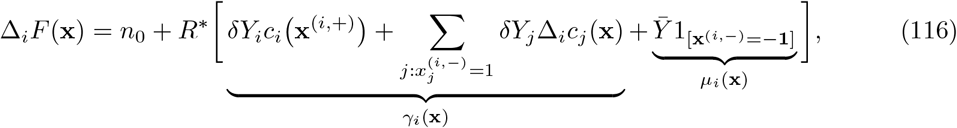

where *γ*_*i*_(**x**) accounts for resource-mediated species interactions, which are determined by yield deviations (***δY*** ), while *µ*_*i*_(**x**) accounts for the remaining contribution from monocultures (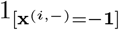 checks that **x**^(*i*,−)^ is the empty community). In vectorial form,

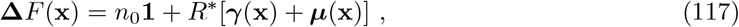

with

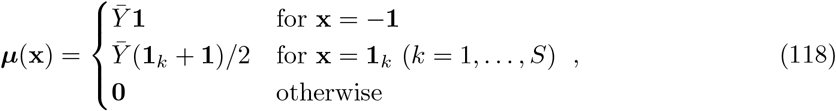

where **1**_*k*_ is a monoculture vector, having 1 at entry *k* and −1 elsewhere, and (**1**_*k*_ + **1**)*/*2 is thus 1 at *k* and 0 elsewhere.

It is convenient to express the consumed fraction as *c*_*i*_ (**x**) = *G*_*i*_ (**x**) */*(*R*^∗^*Y*_*i*_), being 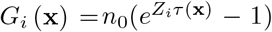 the total biomass produced by species *i* in community **x**. By introducing the vector **y** of entries *y*_*i*_ = ***δ****Y*_*i*_*/Y*_*i*_, and the matrices **K**(**x**), of elements 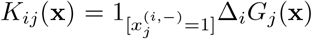, and **A**(**x**) = [diag ({*G*_*i*_(**x**^(i,+)^)}_*i*_) + **K**(**x**) */R*^∗^, we can express the interaction term ***γ*** as linear transformation of **y**, i.e., ***γ***(**x**) = **A**(**x**)**y**. In this form, the dependence of ***γ*** on per-capita growth rates (**Z**) is isolated within **A**.

Let us now compute **M** = ⟨**Δ***F* **Δ***F*⟩^⊤^ _**x**_ by averaging across communities. The vector ***µ*** has first and second moments 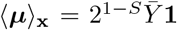 and 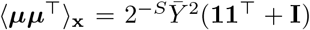, being **I** the identity matrix. We thus get

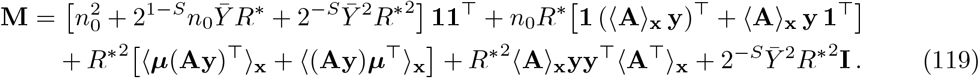

Equation (119) provides an exact expression that sets the stage to connect the leading eigenmode **u**_1_ of **M** to the yield vector **Y**. In fact, it already reveals that the structure of **M** is primarily governed by yield heterogeneity (**y**), whereas variation in per-capita growth rate alone (entering only through **A**) does not affect **M**. In Sec. S8.1.1, we use perturbation theory to make this statement precise in the limit of small yield variation. Then, in Sec. S8.1.2, we show how **u**_1_ can be accurately predicted from **Y** also in a regime where yields and per-capita growth rates are strongly correlated.

##### Numerical integration of the model

Equations (109)-(110) were integrated numerically using the 4th-order Runge-Kutta method. Integration was stopped when *R* reduced to 1% of its initial value *R*(0), i.e., *R*^∗^ = 0.01 *R*(0). The integration was performed for each of the 2^*S*^ species combinations and the biomass landscape computed. Species present in a community were all assigned a starting biomass *n*_0_ = 0.001. The function *q* was chosen according to a standard Michaelis–Menten kinetics, *q*(*R*) = *R/*(*K* + *R*), being *K* = 0.01 the half-saturation constant. Species traits **Y** and **Z** were sampled from uniform distributions of fixed means and variable standard deviations to allow us to systematically adjust their coefficients of variation. The means were fixed to a scale consistent with that observed experimentally, respectively, 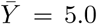 and 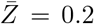, upon setting *R*(0) = 0.05. Standard deviations of both were instead varied through steps of 0.15 from 0 to, respectively, 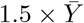 and 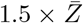, so that both CV(**Y**) and CV(**Z**) took 51 equally-spaced values between 0 and 1.5. For each of these values, we sampled 10 times the vectors **Y** and **Z**, resulting in a total of 510 × 510 = 260100 simulated landscapes. Results for *S* = 5 species are shown in Fig. 3B of the main text and in Fig. S16. It is worth noticing that the values chosen for the parameters entering the model are not critical from a qualitative standpoint.

#### S8.1.1 Small yield heterogeneity: yield-driven regime

Let *y*_*i*_ ≪ 1 for any species *i* and expand Eq. (119) with respect to the small parameter ||**y**|| ≪ 1 (||·|| denotes vector norm). At first order in 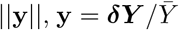, which is orthogonal to **1**. Correspondingly, **M** = **M**^(0)^ + **M**^(1)^, with [**M**^(0)^ ]_*ij*_ ∈ *O*(1) and [**M**^(1)^ ]_*ij*_ ∈ *O*(||**y**||), ∀(*i, j*); precisely,

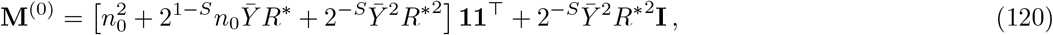

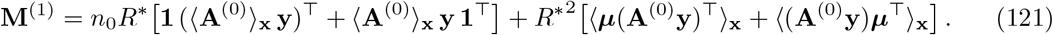

Since any *O*(||**y**||^2^) is discarded, matrix **A** only enters in its unperturbed form **A**^(0)^, which exclusively depends on growth rates.

The unperturbed matrix, **M**^(0)^, has leading eigenvector 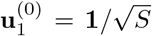 and eigenvalue 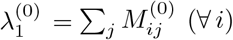, while the remaining *S* − 1 eigenvectors span the degenerate orthogonal subspace associated to the eigenvalue 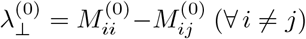, leading to a large spectral gap 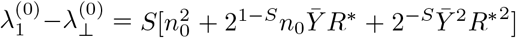 that increases roughly linearly with pool size *S*. We can thus resort to non-degenerate perturbation theory to estimate **u**_1_ through the first-order expansion 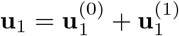. The first-order correction reads

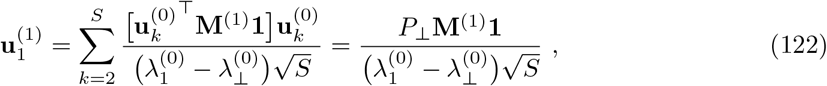

where in the second step we inserted the orthogonal projector 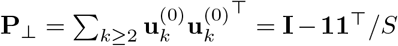, from the identity 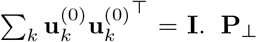 filters out the component of **M**^(1)^**1** parallel to **1**, while acting as the identity on the orthogonal subspace. Naming *c*_∥_**v**_∥_ such component and *c*_⊥_**v**_⊥_ the orthogonal one (||**v**_∥_|| = ||**v**_⊥_|| = 1), **P**_⊥_**M**^(1)^**1** = *c*_⊥_**v**_⊥_. Importantly, since ||**M**^(1)^**1**|| ∈ O(||**y**||), *c*_⊥_ ∈ O(||**y**||) as well.

Upon the perturbation, **u**_1_ thus reads

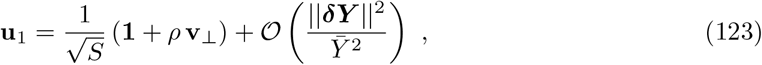

With 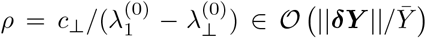. The yield vector, on the other hand, tilts as 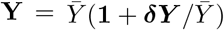. Denoting with 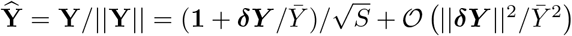 the normalized yield vector, we can thus state the main result of this section:

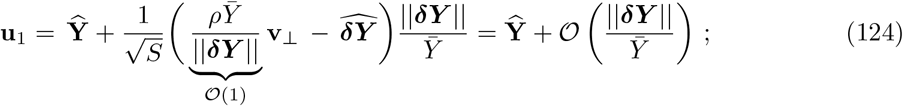

and accordingly,

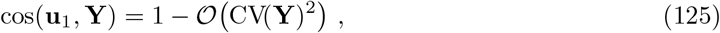

Like **Y, u**_1_ rotates towards the subspace orthogonal to **1** (specifically, in the direction picked up by **M**^(1)^**1**) by an angle whose magnitude is of the order of 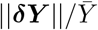. The cosine of the angle between **Y** and **u**_1_ thus decreases by an amount of the order of CV(**Y**)^2^, that is, only quadratically in yield heterogeneity. As shown in Figs. S16(A)-(B), numerical simulations corroborate our analytical prediction.

As a special case, we show below that the perturbation **v**_⊥_ to **u**_1_ becomes exactly 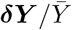 when the per-capita growth rate *Z* either takes the same value for all species or, more generally, it is drawn from a permutation-symmetric distribution, so that species becomes indistinguishable (except for their yield) upon ensemble averaging.

In a nutshell, the across-species variation in per-capita growth rates determines the direction of the perturbation to **u**_1_, yet it is the across-species variation in yields to establish the order of magnitude of such perturbation. As a consequence, the misalignment between **u**_1_ and **Y** is small as long as CV(**Y**) is, no matter how growth rates are assigned or distributed.

##### Permutation-symmetric distribution of per-capita growth rates

Special cases of such distributions are i.i.d. *Z*s or identical *Z*s. Under ensemble expectation, species are indistinguishable and matrix **A**^(0)^ collapses into the permutation-symmetric form **A**^(0)^(**x**) = (*G*(**x**)*/R*^∗^) **I** + (*K*(**x**)*/R*^∗^) (**11**^⊤^ − **I**); respectively, 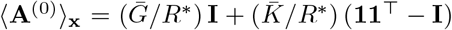. Observing that **y** is an eigenvector of both **P**_⊥_ and **A**^(0)^, the four terms in Eq. (121) acts on **1** to give:

1. [**1** (⟨**A**^**(0)**^⟩_**x**_ **y**)^⊤^] **1**
2. 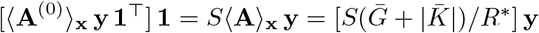 (where we used 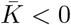)
3. [⟨***µ***(**A**^(0)^**y**)^⊤^⟩_**x**_] **1** = ⟨[*S*(*G* + |*K*|)*/R*^∗^] ***µ***⟩_**x**_ (**y**^⊤^**1**) = 0
4. 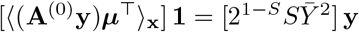

Applying **P**_⊥_ on the left, the first term is annihilated, while the second and fourth are unaffected. Putting all together, we get

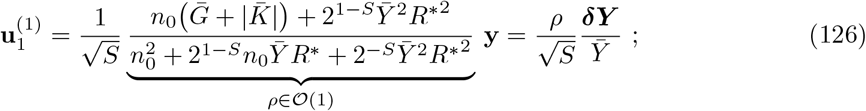

and finally

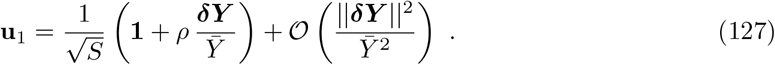

Comparing with Eq. (123), we see that 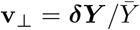 in this case. The leading eigenmode **u**_1_ thus rotates towards the same direction +***δY*** (since *ρ >* 0) as **Y**. Specifically, 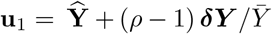 and cos(**u**_1_, **Y**) = 1 − [(*ρ* − 1)^2^*/*2] CV(**Y**)^2^.

#### S8.1.2 Strongly correlated yields and per-capita growth rates: hierarchical regime

Let us assume that yields and per-capita growth rates are strongly correlated, namely **Z** = *a***Y** +**V**, with **V** ⊥ **Y** such that ||**V**|| ≪ *a*||**Y**||. Conveniently, let us label species from most to least dominant, i.e., in descending order of per-capita growth rate (therefore of yield), so that *Z*_1_ *>* · · · *> Z*_*S*_ (*Y*_1_ *>* · · · *> Y*_*S*_). Let 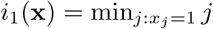 denote the dominant species in community **x**.

Assuming that CV(**Z**) (hence CV(**Y**)) is large and there is enough resource to guarantee 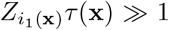 (equivalently 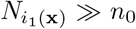), from 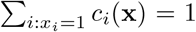 we obtain the approximation 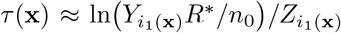, with a relative error smaller than 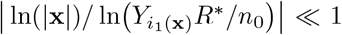 (notice |**x**| indicates the number of species in **x**). Consequently, 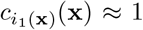, and *c*_*j*_(**x**) ≈ 0 for any (non-dominant) species *j > i*_1_(**x**) in **x**. That is, virtually all the available resource is depleted by the dominant species in the community, while the others barely grow from the inoculum. The matrix **A**(**x**) becomes

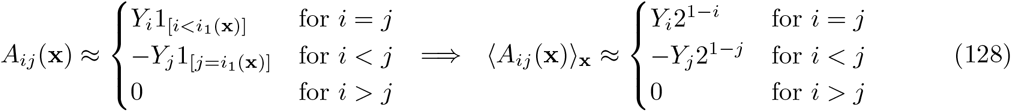

being 2^1−*j*^ the fraction of communities in which species *j* is dominant (species *j* is dominant if all species from 1 to *j* − 1 are absent). The factor 2^1−*j*^ reflects the competitive hierarchy among species, weighting them geometrically from the most to the less dominant one.

Recalling the assumption of abundant resource, which implies *Y*_*i*_*R*^∗^ ≫ *n*_0_ for any *i*, and assuming 2^−*S*^ ≪ 1, Eq. (119) reduces to

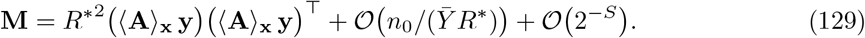

At leading order, **M** is thus a rank-1 matrix whose principal eigenvector **u**_1_ is parallel to the approximate focal effects vector **f** ^(rank)^ = *R*^∗^⟨**A**⟩_**x**_ **y**. Using Eq. (128),

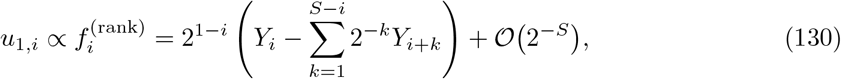

which is entirely determined by yields. Therefore, once measured the yield of each species from monocultures, we can accurately reconstruct the leading collective mode from Eq. (130). As shown in Fig. S16(C), numerical simulations confirm this prediction.

#### S8.1.3 Multiple resources

The results in Secs. S8.1.1-S8.1.2 extend directly to the case of multiple shared resources. Given *L* resources, {*R*_*r*_}_*r*=1,…,*L*_, the dynamical equations read

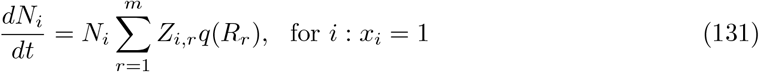

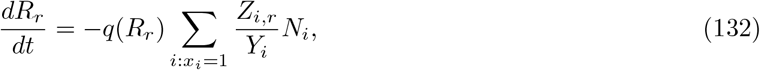

where *Z*_*i,r*_ is the (maximum) per-capita growth rate of species *i* when consuming resource *r*, and *Y*_*i*_ is the yield of species *i* in the environment specified by the *m* present resources. By the latter we mean that *L* resources together define an environment and in such environment we measure a community-function landscape. Then, the *S* yields are computed from the *S* monocultures in such environment. Therefore, we can always assign to species a measurable, ecologically meaningful value of yield ({*Y*_*i*_}_*i*_) within each landscape (environment), while keeping the heterogeneity across resources in the values of the growth rates ({*Z*_*i,r*_}_*i,r*_).

The solution to Eqs. (131)-(132) takes the same form as Eqs. (109)-(110), namely

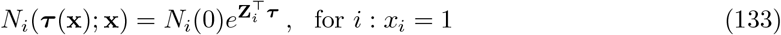

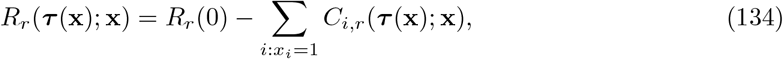

being ***τ*** = (*τ*, …, *τ* ) the vector of effective times, with 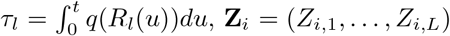 the vector of per-capita growth rates of species *i*, and *C*_*i,r*_ the amount of resource *r* consumed by this species. Assuming that all species are inoculated with the same biomass *n*_0_ and growth is stopped when a fixed total abundance *R*^∗^ of resources has been depleted, a formally identical analysis to the one made in Sec. S8.1 follows in terms of the total amount of resource consumed by each species 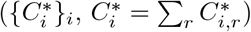.

#### S8.2 Null distribution for pairwise angles

To assess whether the observed alignment between the leading eigenvector **u**_1_ and the yield **Y** exceeds what is expected by chance, we compare empirical angle distributions to a geometric null model corresponding to random directions. In an *S*-dimensional space, the cosine of the angle *ω* between two normalized vectors chosen uniformly at random follows a scaled Beta distribution [2]

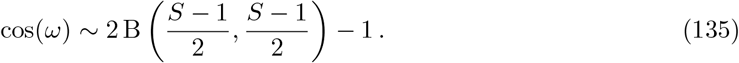

### S9 Analysis of previously published datasets

In addition to the empirical landscapes generated in this work, we applied our framework to six previously published full-factorial community-function datasets [3, 8, 11, 15, 17, 20]. These datasets span distinct microbial systems, community sizes, and measured functions, these including biomass, population size, metabolic activity, and host life-history traits. Community size ranged from *N* = 5 to *N* = 10 components (Table T2). We used these data to test whether the organization of marginal effects observed in our experiments also appears across different ecological systems and functions.

For each focal component *i*, we first evaluated the corresponding Functional Effect Equation defined in Eq. (10). We then transformed each complete landscape into WH interaction space and compared coefficients involving the focal component, *f*_**s**+**i**_, with the corresponding lower-order coefficients, *f*_**s**_. The slopes predicted from the FEEs were calculated using Eq. (15) and compared with the empirical relationships among WH coefficients.

Several focal components exhibited approximately linear FEEs together with the corresponding cross-order organization in interaction space (Figs. S24, S25 and S26). These relationships were observed for functions distinct from those measured in our experiments. For example, the population-size landscapes of Ishizawa et al. [15] quantify how the presence of one community member modifies the abundance of another, bringing the measured function closer to the classical population-dynamical definition of ecological interactions. Similar patterns were also observed for species richness, biomass, and host-associated traits.

The coefficient-level relationships were generally less pronounced than in our replicated experiments. Most published datasets contained a single measurement for each community composition. WH coefficients estimated from unreplicated landscapes are highly sensitive to measurement noise, particularly at high interaction orders [3]. We therefore restricted the interaction-space analysis to first- and second-order background coefficients. These correspond to pairwise and third-order coefficients after addition of the focal component. The resulting comparisons should be interpreted as qualitative evidence of cross-order organization rather than as precise tests of the slopes predicted by the FEEs.

Departures from a single global FEE also revealed structured local rules. In the biomass landscape of Sanchez-Gorostiaga et al. [20], the functional effect of *Paenibacillus polymyxa* separated into two approximately linear branches depending on whether *Bacillus thuringiensis* was present or absent from the ecological background (Fig. S24). Thus, the presence of one species can modify the functional rule followed by another and partition the landscape into distinct local regimes. Such conditional FEEs are not inconsistent with an organized landscape. Instead, they suggest that some systems may be described by a small number of locally valid rules rather than by a single relationship applying across all ecological backgrounds.

For each published landscape, we next constructed the second-moment matrix of marginal effects defined in Eq. (28) and calculated its ecological collective modes. The corresponding spectra were generally concentrated in the first one or two modes (Fig. S30 and S31). We then projected community composition onto the leading latent coordinate defined in Eq. (62). The resulting one-dimensional projections, obtained using Eq. (63), generally organized community function more effectively than species biodiversity alone (Fig. S27 and Fig. S29). When the second mode carried appreciable spectral weight, the two-dimensional latent manifolds defined in Eq. (74) captured additional landscape structure (Fig. S28).

The *Drosophila* microbiome dataset of Gould et al. [11] included several host traits measured over the same set of microbial communities. Total fecundity, daily fecundity, and development time displayed distinct community-function landscapes, but all three showed concentrated marginal-effect spectra and could be represented using one or two latent coordinates (Fig. S29). This illustrates that collective modes are function-dependent, while different functions defined over the same compositional space can nevertheless exhibit comparable low-dimensional organization.

Together, these reanalyses reveal a clear separation between coefficient-level and landscape-level inference. Approximate FEEs and their interaction-space counterparts were detectable across systems, but were often obscured by the lack of replication and the resulting uncertainty in individual WH coefficients. By contrast, the spectral organization of the marginal-effect matrix **M** was consistently recovered: marginal effects concentrated along a small number of collective directions, and the resulting one- or two-dimensional projections provided coherent representations of every published landscape examined.

## S10 Supplementary Tables

**Table T1:** List of carbon sources used in this study.

| Reagent type | Carbon source | 1×<br>C-mol/L | 5×<br>C-mol/L | Supplier | Reference |
| --- | --- | --- | --- | --- | --- |
| Chem. compound | CH<sub>3</sub>COONa | 0.014 | 0.07 | Sigma-Aldrich | W302406 |
| Chem. compound | NaC<sub>6</sub>H<sub>7</sub>O<sub>7</sub> | 0.014 | 0.07 | Sigma-Aldrich | 71498 |
| Chem. compound | D-(–)-Fructose | 0.014 | 0.07 | Sigma-Aldrich | F0127 |
| Chem. compound | D-(+)-Glucose | 0.014 | 0.07 | Sigma-Aldrich | G7021 |
| Chem. compound | Glycerol | 0.014 | 0.07 | Sigma-Aldrich | G9012 |

**Table T2:**
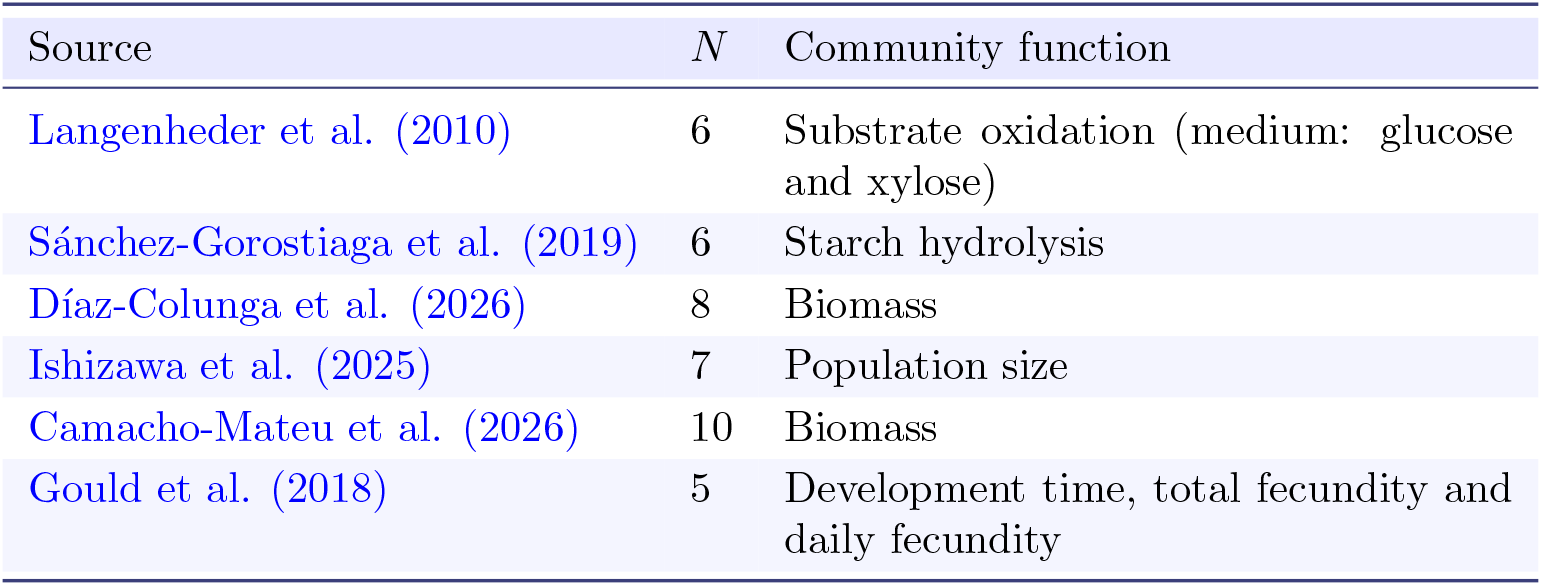
Previously published community-function landscapes analyzed in this work.

## S11 Supplementary Figures

**Figure S1:**
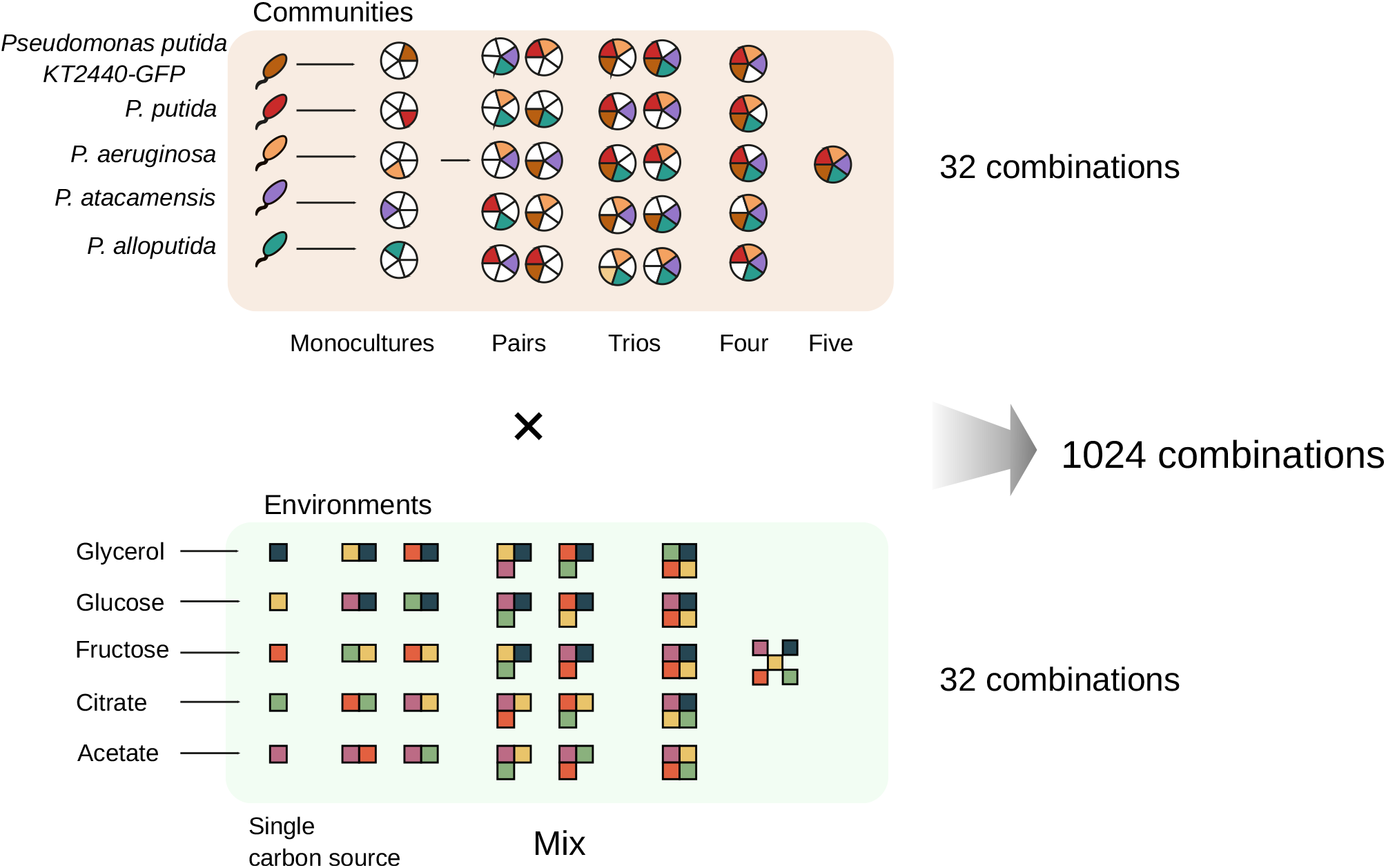
Full-factorial design of the five-species–five-resource combinatorial assembly experiment. See Sec. S1.2. Five *Pseudomonas* species were assembled in all 2^5^ = 32 possible presence–absence combinations, ranging from the empty community to the complete five-species community. Independently, five carbon sources—glycerol, glucose, fructose, citrate, and acetate— were supplied in all 2^5^ = 32 possible presence–absence combinations, ranging from no-resource controls to mixtures containing all five resources. Crossing the species and resource landscapes generated 2^5^ × 2^5^ = 1,024 community–environment combinations. Community function was set to zero for conditions containing no species or no supplied carbon source.

**Figure S2:**
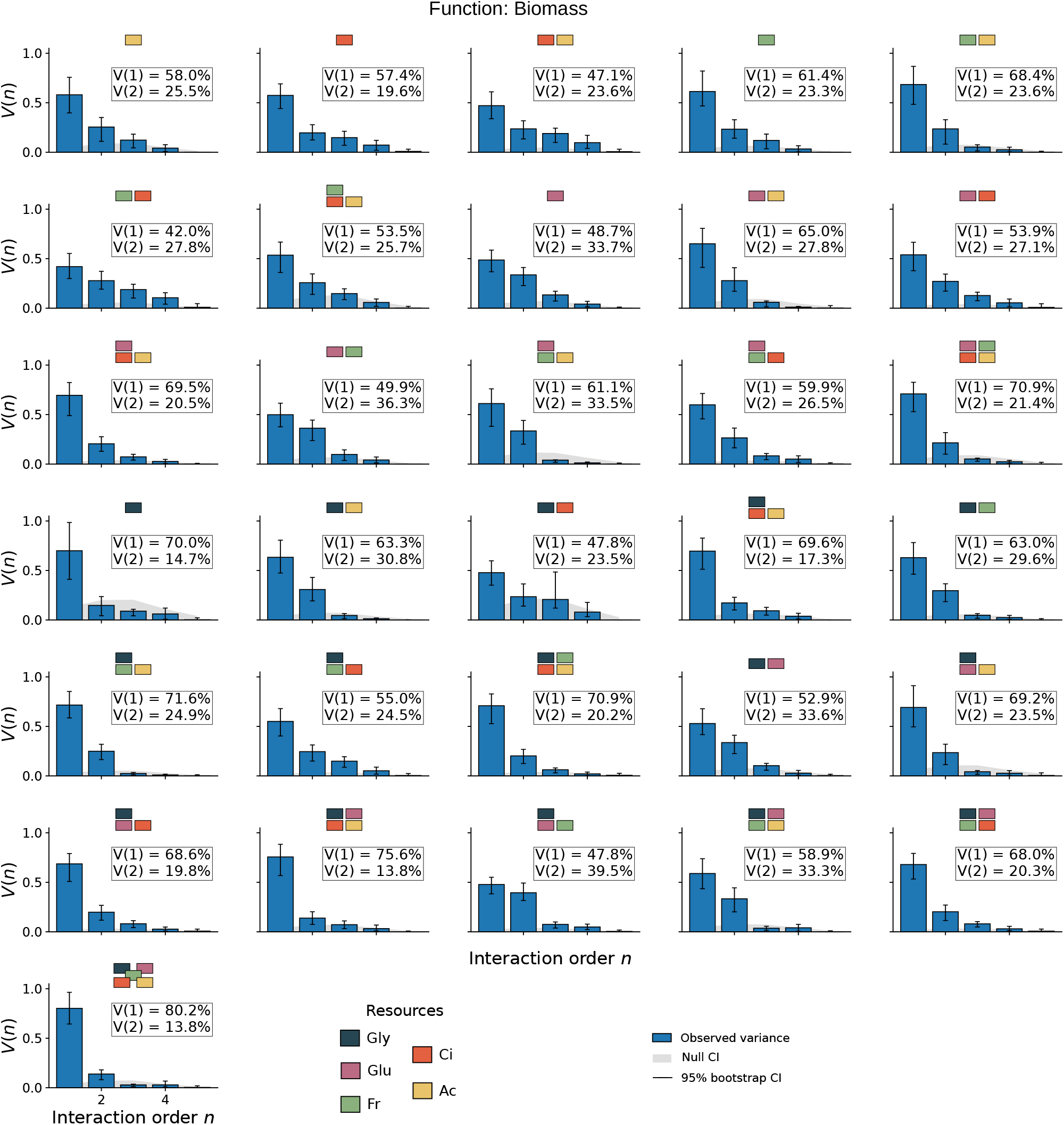
Variance decomposition of biomass community-function landscapes across WH interaction orders. For each resource environment, the biomass community-function landscape was decomposed into WH coefficients grouped by interaction order *n*. Bars show the fraction of total landscape variance contributed by coefficients of order *n, V* (*n*), as defined in Eq. (9). Error bars indicate 95% bootstrap confidence intervals, and gray bands show the corresponding null expectation (both the null model and the bootstrap confidence intervals are computed following [3]). Resource composition is indicated above each panel by colored squares. Labels report the contributions of first- and second-order coefficients, *V* (1) and *V* (2). Across environments, the first two interaction orders together accounted for most of the total variance, supporting the second-order WH approximation used in the spectral and latent-coordinate analyses.

**Figure S3:**
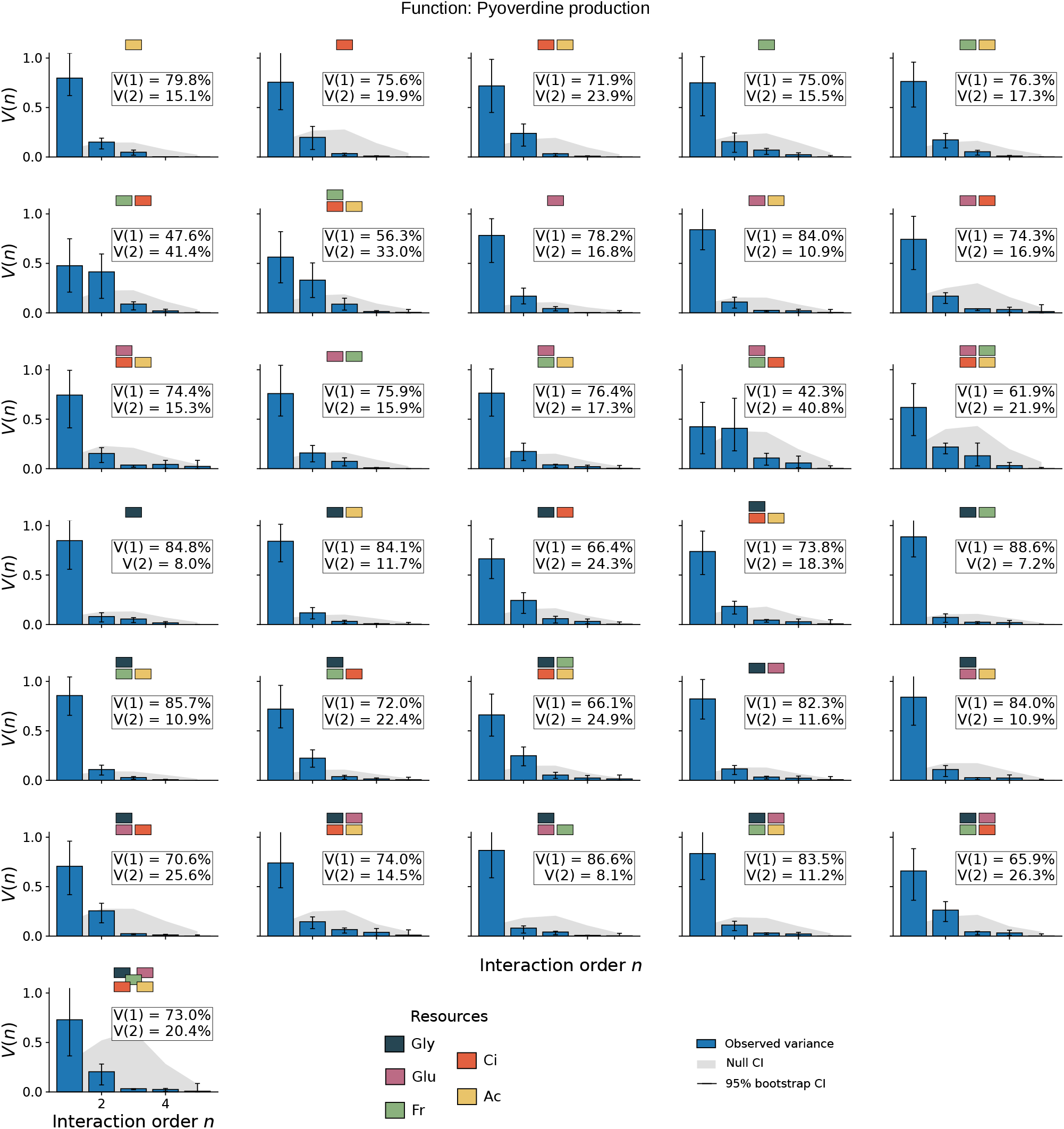
Variance decomposition of pyoverdine community-function landscapes across WH interaction orders. For each resource environment, the pyoverdine community-function landscape was decomposed into WH coefficients grouped by interaction order *n*. Bars show the fraction of total landscape variance contributed by coefficients of order *n, V* (*n*), as defined in Eq. (9). Error bars indicate 95% bootstrap confidence intervals, and gray bands show the corresponding null expectation. Resource composition is indicated above each panel by colored squares. Labels report the contributions of first- and second-order coefficients, *V* (1) and *V* (2). Across most environments, these two orders together accounted for approximately 90% or more of the total variance, with higher-order coefficients contributing only a small residual fraction. This concentration supports the second-order Walsh–Hadamard approximation used in the spectral and latent-coordinate analyses of pyoverdine production.

**Figure S4:**
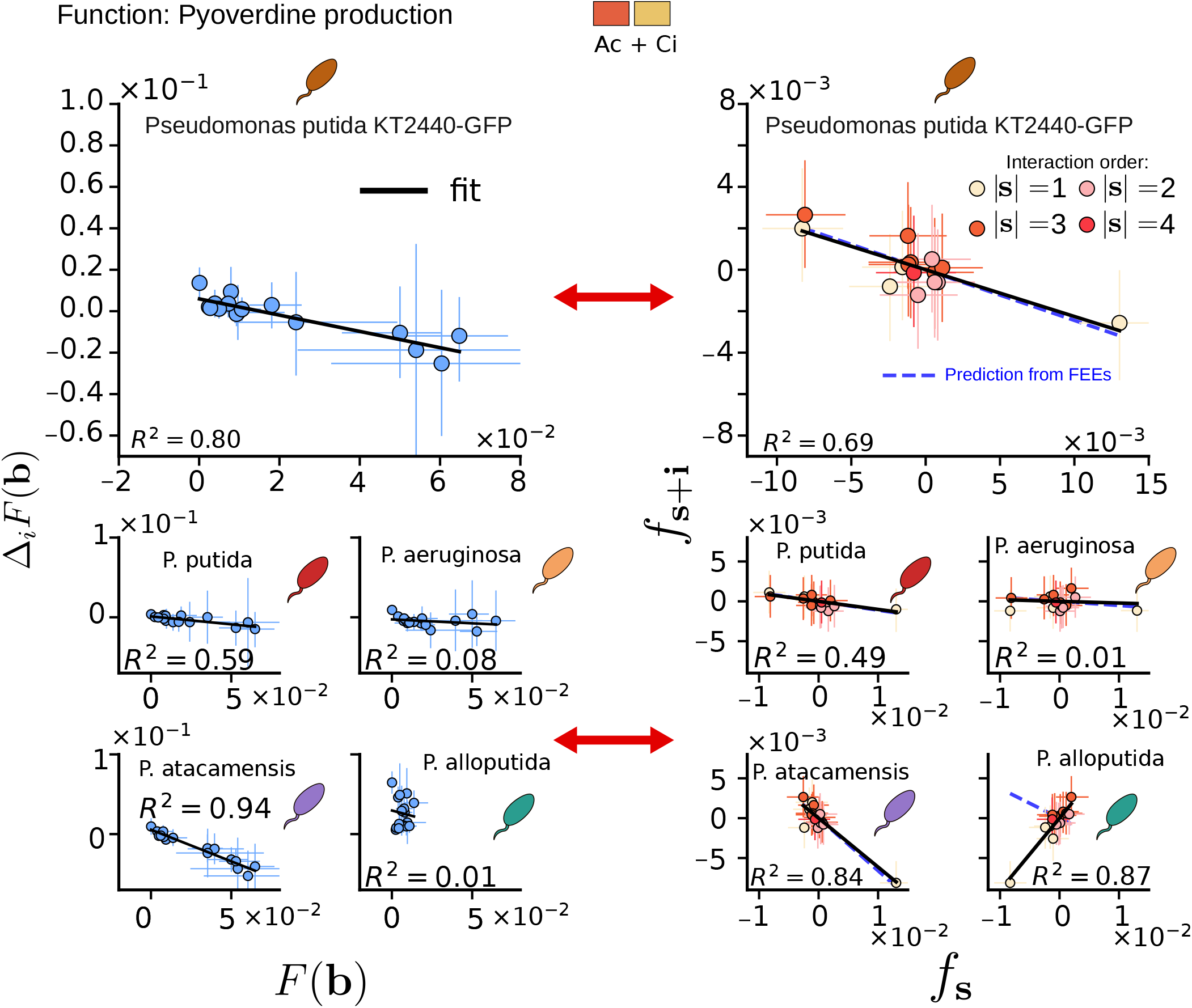
FEEs predict a hierarchical organization of interaction coefficients for pyoverdine production. This figure complements Fig. 1 of the main text, which presents the corresponding analysis for biomass. Here, the same analysis is shown for pyoverdine production in citrate + acetate medium. Left panels show, for each focal species *i*, its functional effect, Δ_*i*_*F* (**b**), as a function of the background community function, *F* (**b**), as defined in Eq. (10). Right panels show the corresponding relationships in interaction space by comparing WH coefficients involving the focal species, *f*_**s**+**i**_, with the associated coefficients one order below, *f*_**s**_, as predicted by Eq. (15). Solid black lines show empirical linear fits, whereas dashed blue lines indicate the slopes predicted from the fitted FEE slope *b*_*i*_. Error bars denote 95% bootstrap confidence intervals, computed as described in [3].

**Figure S5:**
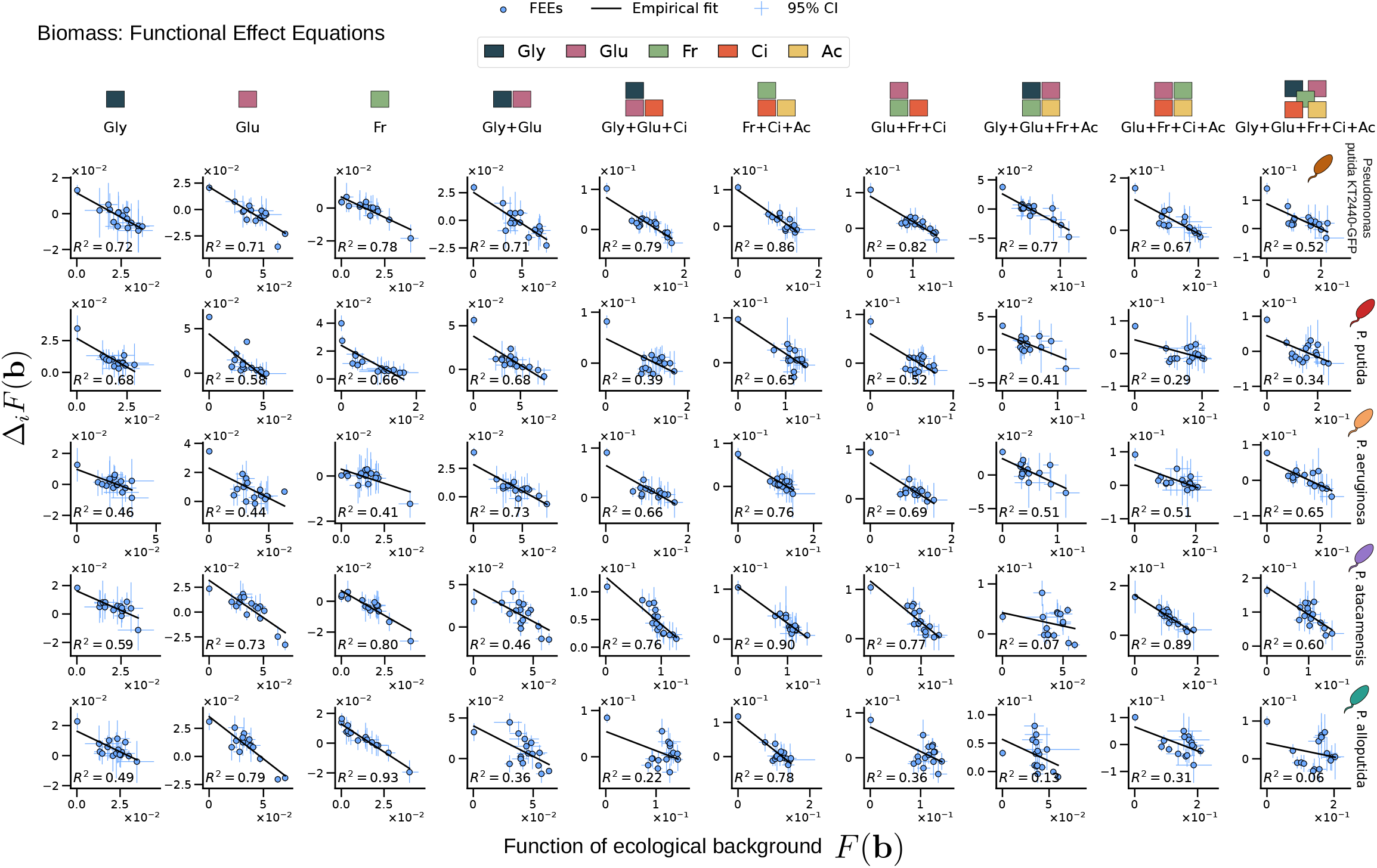
Functional Effect Equations for biomass production across randomly resource environments. This figure extends the FEE analysis shown in Fig. 1B to additional resource environments. Each column corresponds to a different resource environment, and each row to a focal species. Resource composition is indicated above the columns by colored rectangles denoting the carbon sources present in each medium—glycerol, glucose, fructose, citrate, and acetate—. In each panel, the functional effect of adding focal species *i*, Δ_*i*_*F* (**b**), is plotted against the function of the corresponding background community, *F* (**b**). Points show measured functional effects, vertical error bars denote 95% bootstrap confidence intervals, and solid black lines show empirical linear fits. The coefficient of determination, 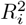, is reported in each panel.

**Figure S6:**
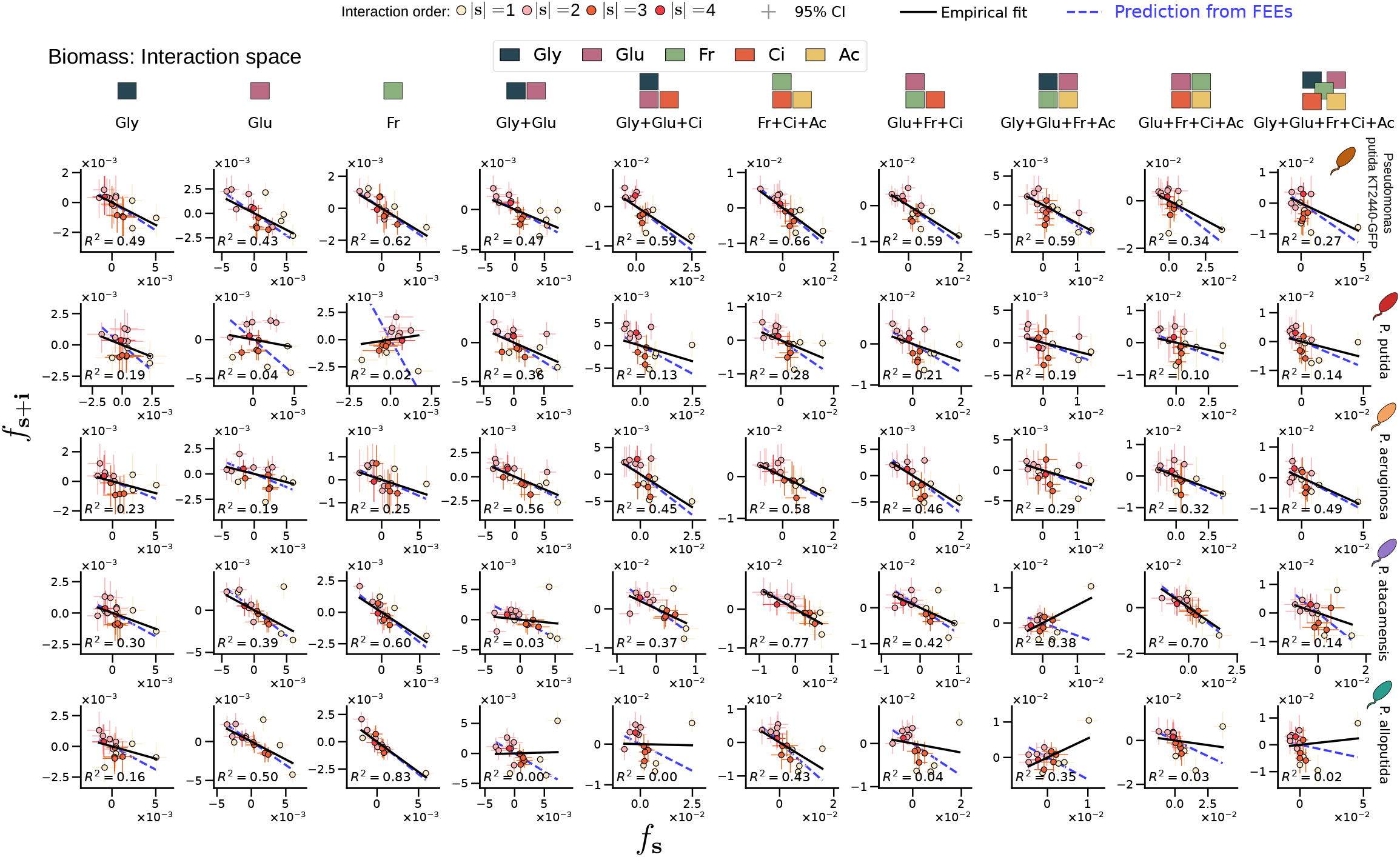
WH interaction coefficients follow simple cross-order relationships for biomass production across randomly selected resource environments. This figure extends the WH coefficient analysis shown in Fig. 1C to a representative subset of resource environments. Each column corresponds to a different resource environment, and each row to a focal species. Resource composition is indicated above the columns by colored rectangles denoting the carbon sources present in each medium—glycerol, glucose, fructose, citrate, and acetate. In each panel, WH coefficients involving focal species *i, f*_**s**+**i**_, are plotted against the corresponding coefficients one order below, *f*_**s**_, for subsets **s** not containing species *i*. Points show estimated WH coefficients and are colored by interaction order. Solid black lines show empirical linear fits, whereas dashed blue lines indicate the slopes predicted from the corresponding fitted FEE slope *b*_*i*_, according to Eq. (15). Vertical error bars denote 95% bootstrap confidence intervals. The coefficient of determination of the interaction-space fit, 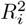, is reported in each panel.

**Figure S7:**
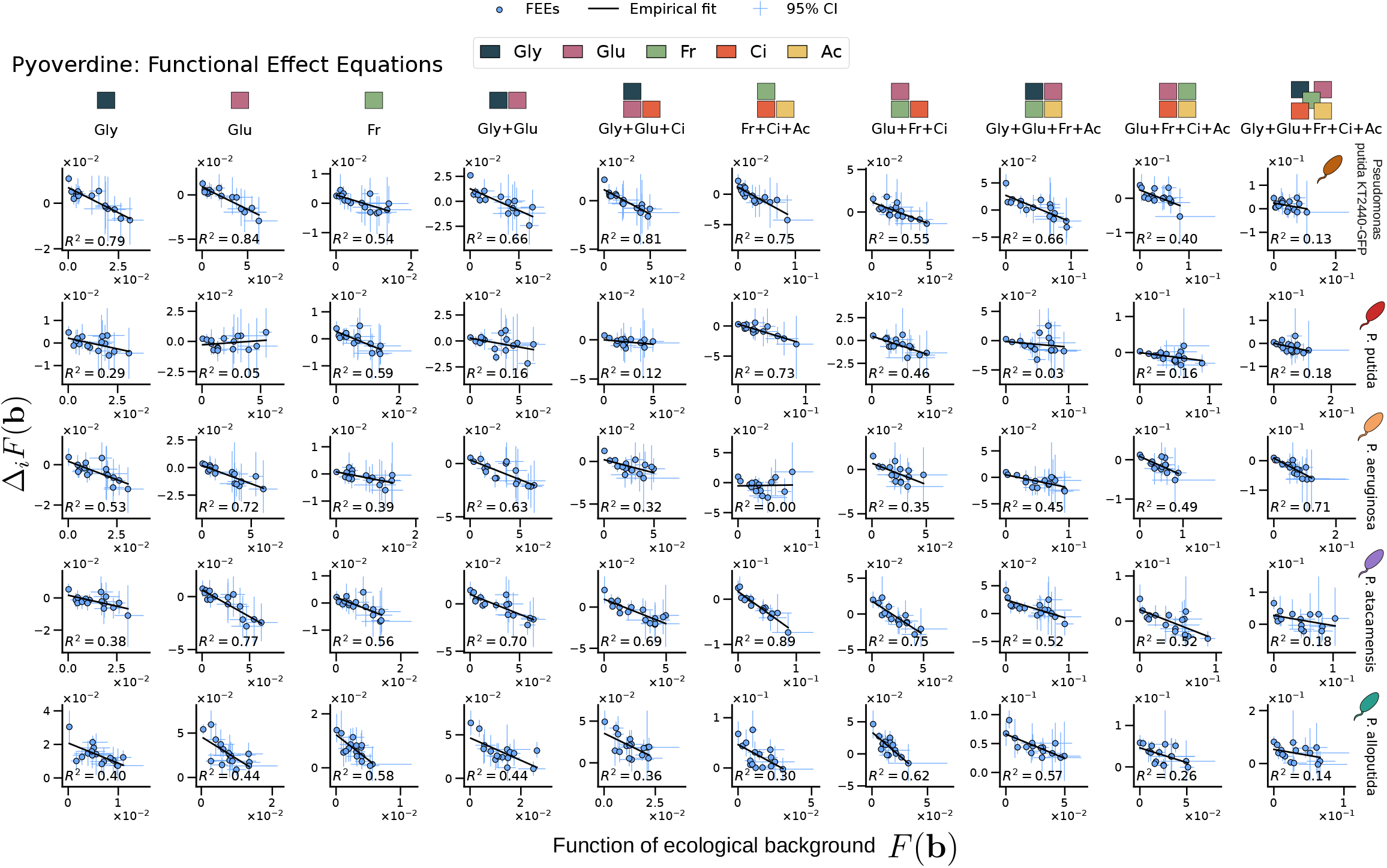
Functional Effect Equations for pyoverdine production across randomly selected resource environments. This figure presents the same analysis as Fig. S5, using pyoverdine production as the community function. Each column corresponds to a different resource environment, and each row to a focal species. In each panel, the functional effect of adding focal species *i*, Δ_*i*_*F* (**b**), is plotted against the function of the corresponding background community, *F* (**b**). Points show measured functional effects, vertical error bars denote 95% bootstrap confidence intervals, and solid black lines show empirical linear fits. The coefficient of determination, 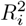, is reported in each panel.

**Figure S8:**
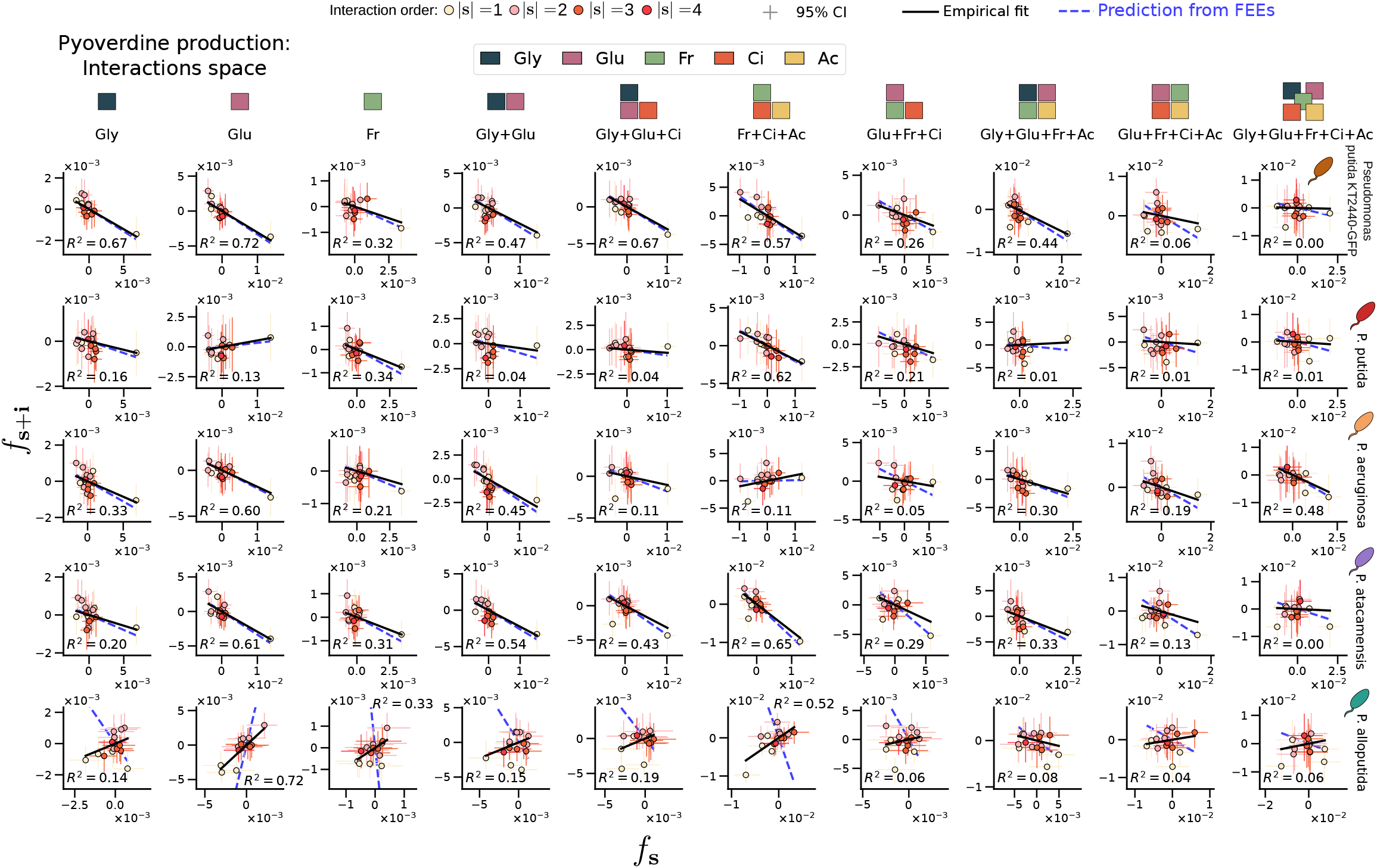
WH interaction coefficients follow simple cross-order relationships for pyoverdine production across randomly selected resource environments. This figure presents the same analysis as Fig. S6, using pyoverdine production as the community function. Each column corresponds to a different resource environment, and each row to a focal species. In each panel, WH coefficients involving focal species *i, f*_**s**+**i**_, are plotted against the corresponding coefficients one order below, *f*_**s**_, for subsets **s** not containing species *i*. Points show estimated WH coefficients and are colored by interaction order. Solid black lines show empirical linear fits, whereas dashed blue lines indicate the slopes predicted from the corresponding fitted FEE slope *b*_*i*_, using *c*_*i*_ = *b*_*i*_*/*(2 + *b*_*i*_) as given by Eq. (15). Vertical error bars denote 95% bootstrap confidence intervals. The coefficient of determination of the interaction-space fit, 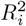, is reported in each panel.

**Figure S9:**
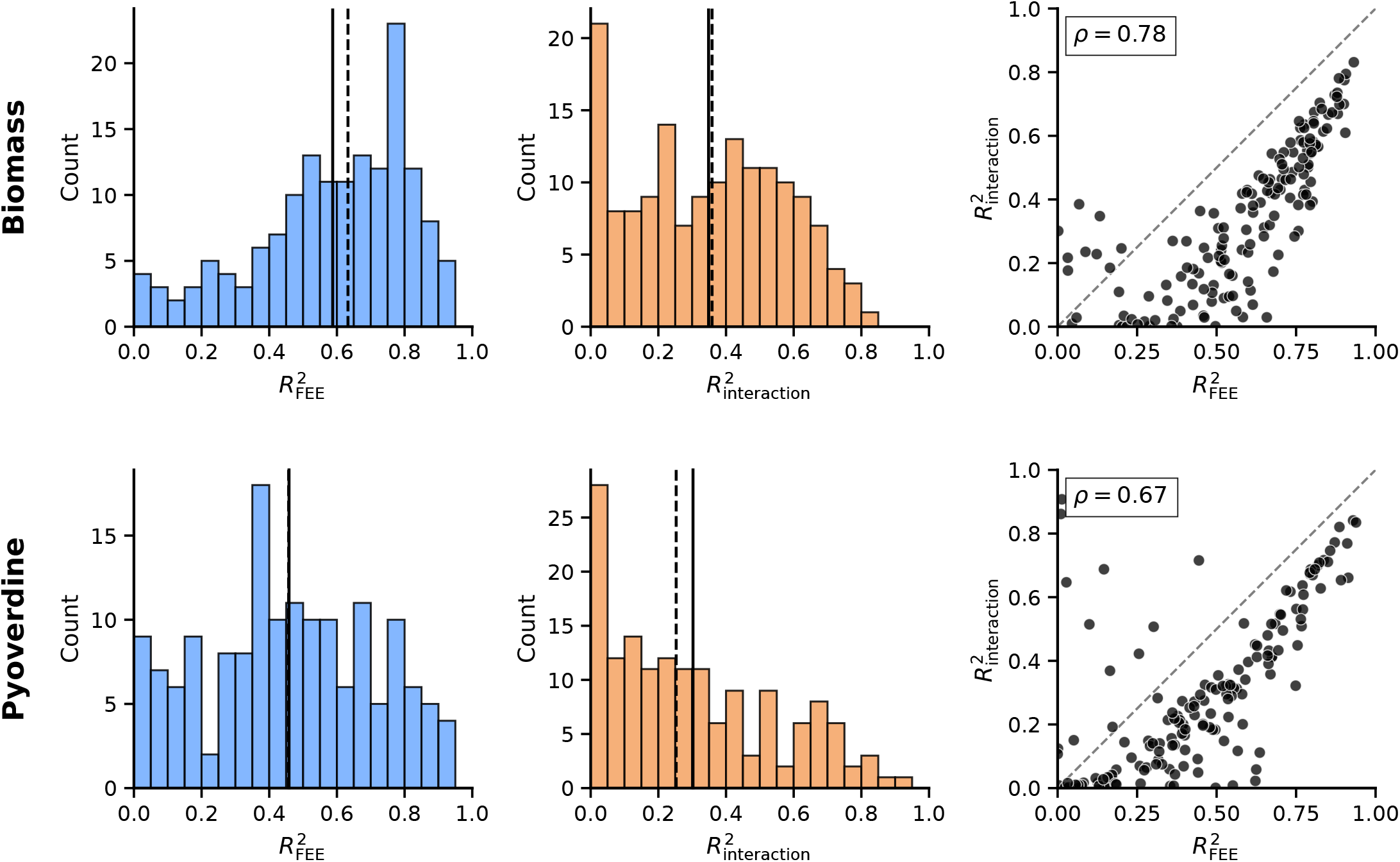
FEE predictability is associated with stronger cross-order relationships among WH interaction coefficients. Distributions of the coefficients of determination for Functional Effect Equations, 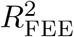, and for the corresponding cross-order relationships among WH interaction coefficients, 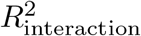, across focal species and resource environments. Top panels show biomass production and bottom panels pyoverdine production. Left and middle panels show the distributions of 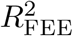 and 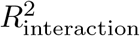, respectively; solid and dashed vertical lines indicate the mean and median. Right panels compare both quantities for each focal species–environment combination. The two measures are strongly positively correlated for biomass (*ρ* = 0.78) and pyoverdine (*ρ* = 0.67), showing that more predictive FEEs are associated with more pronounced cross-order organization in interaction space. Most points fall below the identity line, consistent with the greater statistical uncertainty involved in estimating WH coefficients, particularly at higher interaction orders.

**Figure S10:**
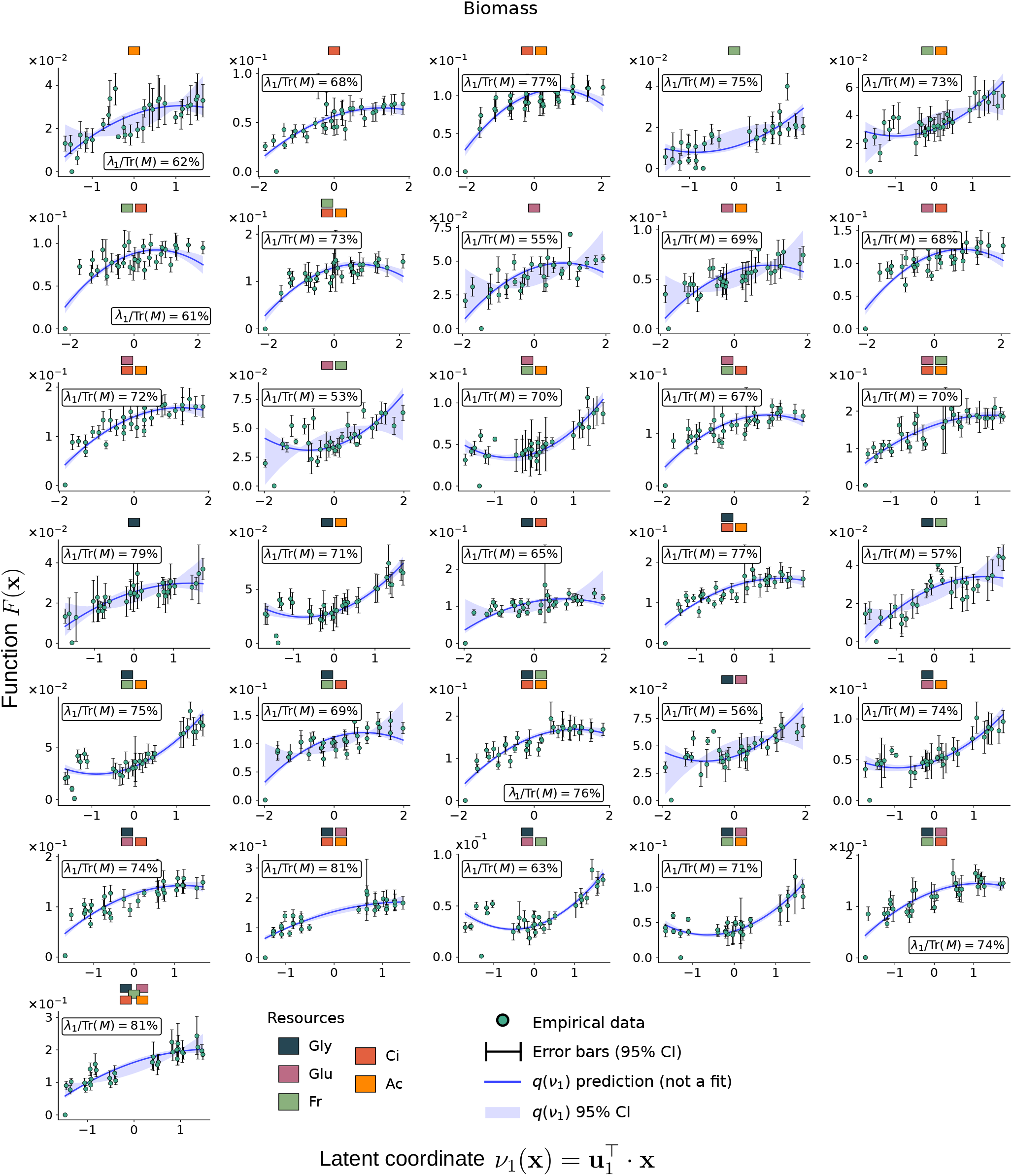
One-dimensional latent-coordinate projections of biomass community-function landscapes across resource environments. Each panel shows a biomass community-function landscape projected onto its leading latent coordinate, 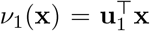. Points show measured community functions, *F* (**x**), and error bars denote 95% bootstrap confidence intervals. Solid blue lines show the latent-coordinate prediction, *q*(*ν*_1_), defined in Eq. (63), and shaded blue regions denote the corresponding 95% bootstrap confidence intervals. The label in each panel reports the fraction of the total spectral weight captured by the leading mode, *λ*_1_*/*Tr(**M**).

**Figure S11:**
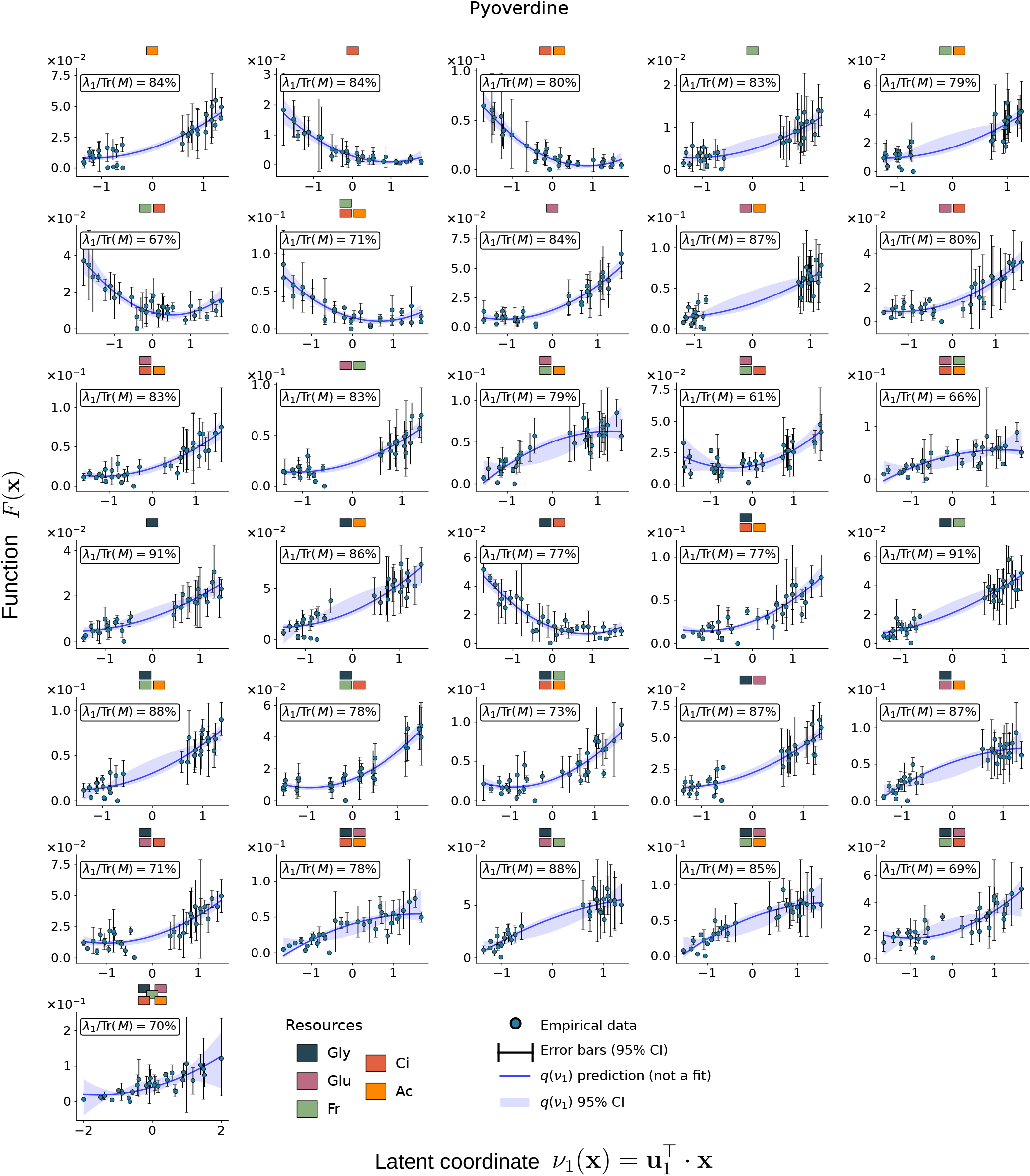
One-dimensional latent-coordinate projections of pyoverdine community-function landscapes across resource environments. Each panel shows a pyoverdine community-function landscape projected onto its leading latent coordinate,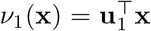. Points show measured community functions, *F* (**x**), and error bars denote 95% bootstrap confidence intervals. Solid blue lines show the latent-coordinate prediction, *q*(*ν*_1_), defined in Eq. (63), and shaded blue regions denote the corresponding 95% bootstrap confidence intervals. The label in each panel reports the fraction of the total spectral weight captured by the leading mode, *λ*_1_*/*Tr(**M**).

**Figure S12:**
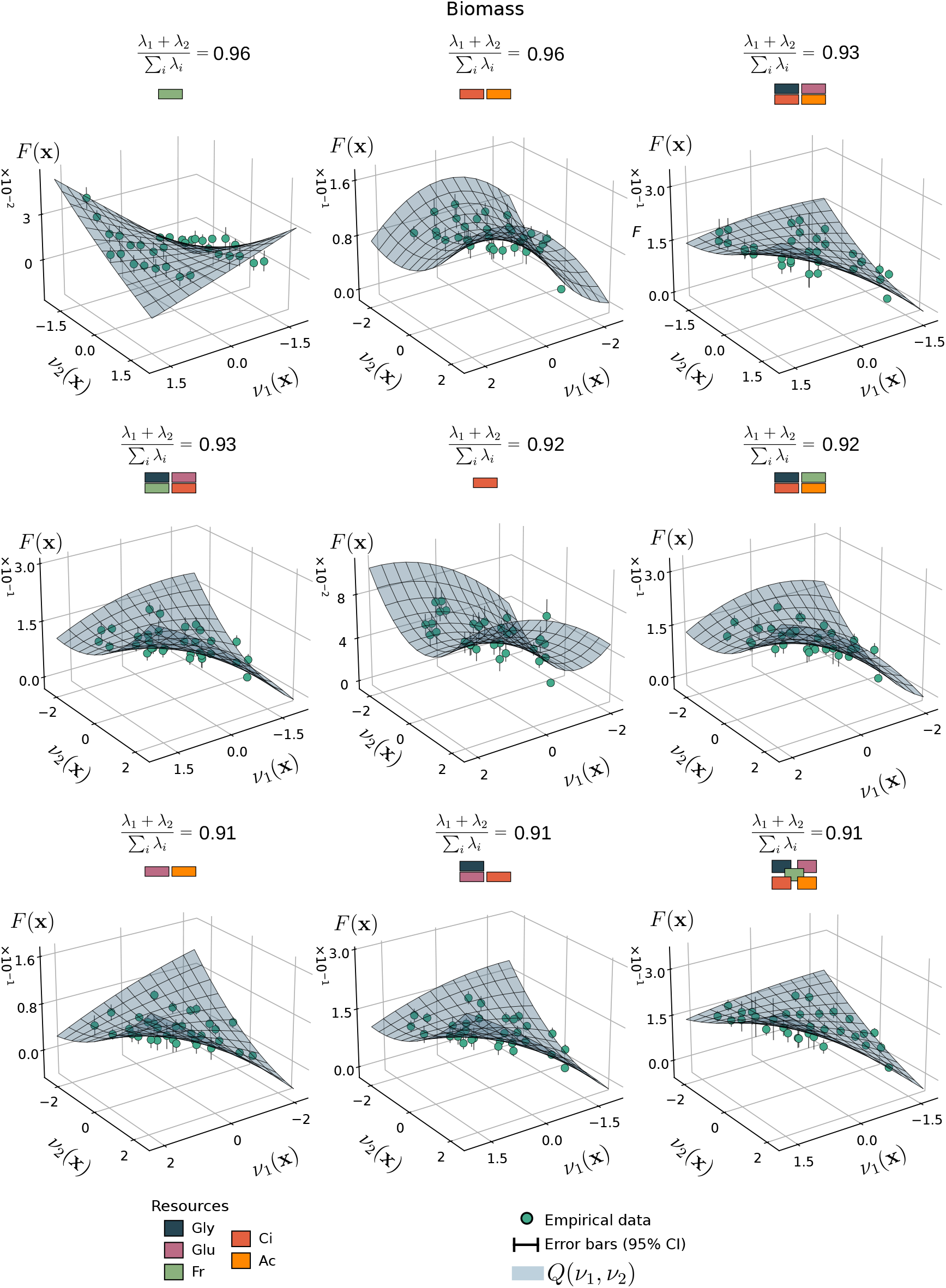
Two-dimensional latent-coordinate manifolds of biomass community-function landscapes across selected resource environments. Each panel shows a biomass community-function landscape projected onto its first two latent coordinates, 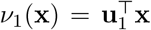 and 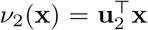. Points show measured community functions, *F* (**x**), after 24 h of growth, and error bars denote 95% bootstrap confidence intervals. Blue surfaces show the predicted quadratic latent manifold, *Q*(*ν*_1_, *ν*_2_), defined in Eq. (74). The nine panels correspond to the resource environments for which the *K* = 2 manifold captured the largest fraction of biomass variation.

**Figure S13:**
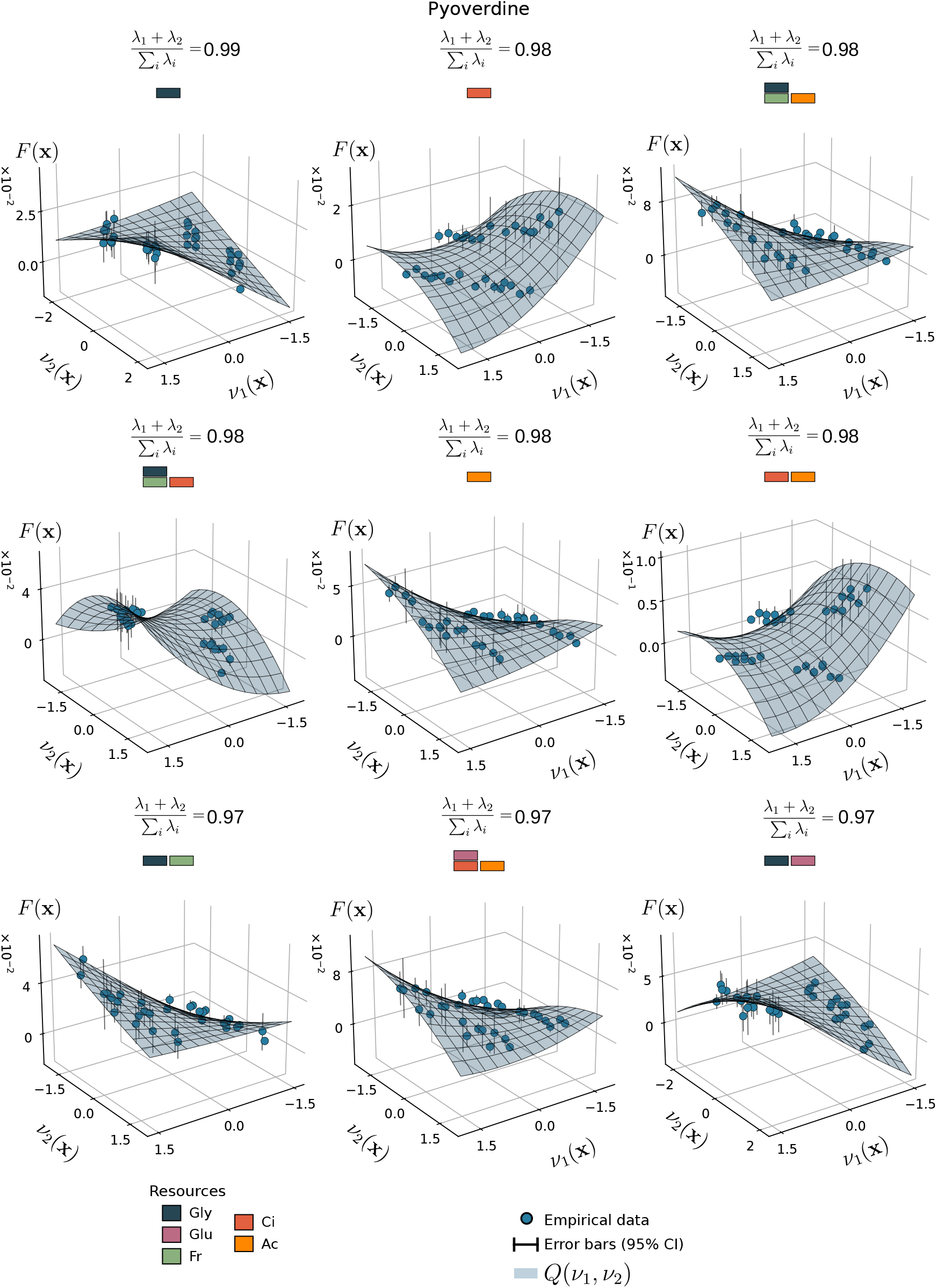
Two-dimensional latent-coordinate manifolds of pyoverdine community-function landscapes across selected resource environments. Each panel shows a pyoverdine community-function landscape projected onto its first two latent coordinates, 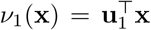 **x** and 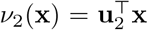. Points show measured community functions, *F* (**x**), after 24 h of growth, and error bars denote 95% bootstrap confidence intervals. Blue surfaces show the predicted quadratic latent manifold, *Q*(*ν*_1_, *ν*_2_), defined in Eq. (74). The nine panels correspond to the resource environments for which the *K* = 2 manifold captured the largest fraction of pyoverdine variation.

**Figure S14:**
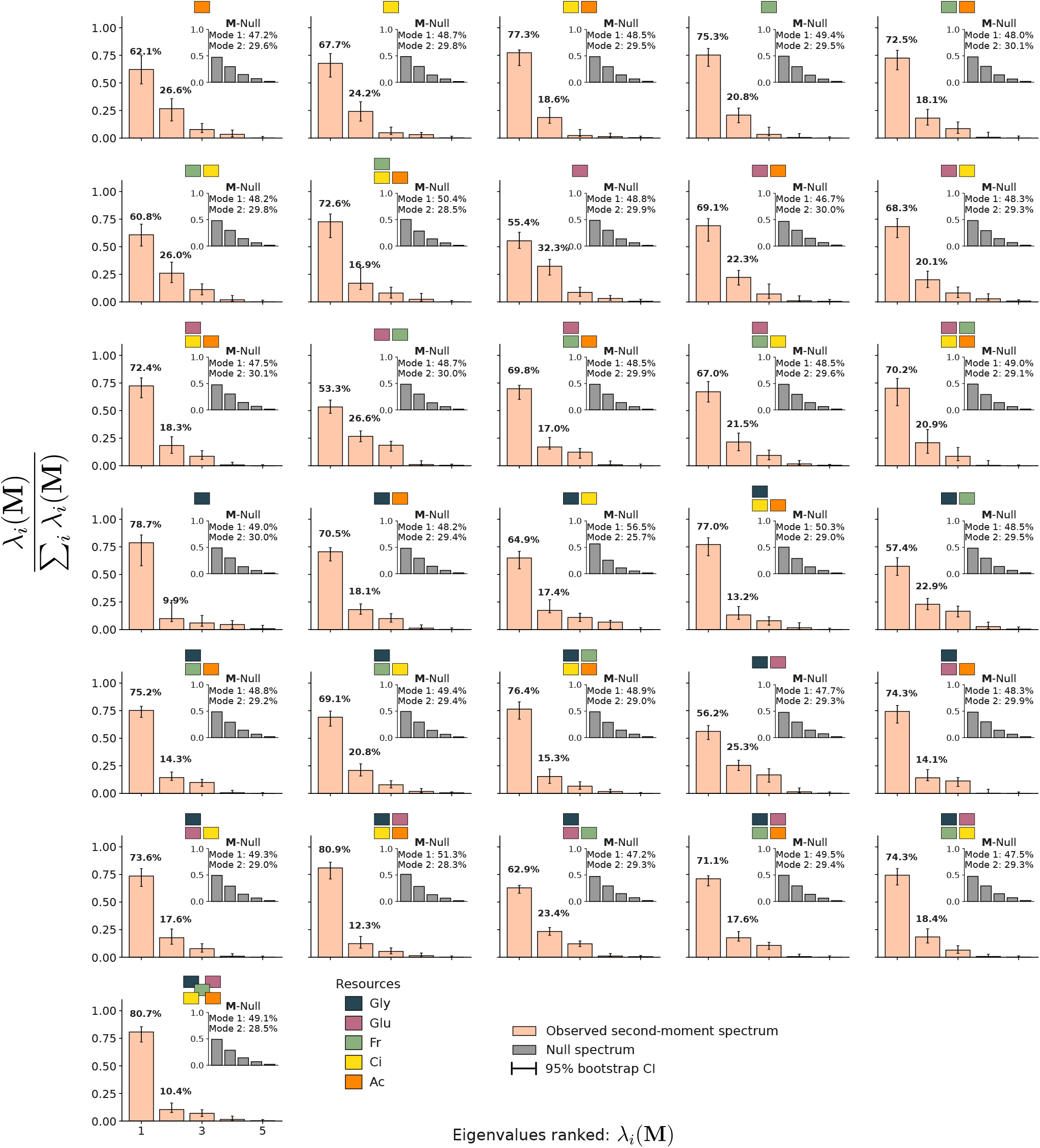
Low-rank second-moment spectra across resource environments. For each resource environment, we approximated the second-moment matrix of marginal effects, **M**, by retaining only first- and second-order WH coefficients. Under this approximation, **M** ≈ **ff** ^⊤^+**HH**^⊤^, as given by Eq. (50), where **f** is the vector of first-order WH coefficients and **H** is the matrix of second-order WH coefficients. Bars show the normalized eigenvalue spectrum, *λ*_*i*_(**M**)*/* _*j*_ *λ*_*j*_(**M**), and error bars denote 95% bootstrap confidence intervals. Insets show the corresponding spectra obtained under the null model, **M**_null_. Across resource environments, the observed spectra are dominated by one or two leading modes, whereas the null spectra are more broadly distributed. Resource composition is indicated above each panel by colored rectangles denoting the carbon sources present in each medium.

**Figure S15:**
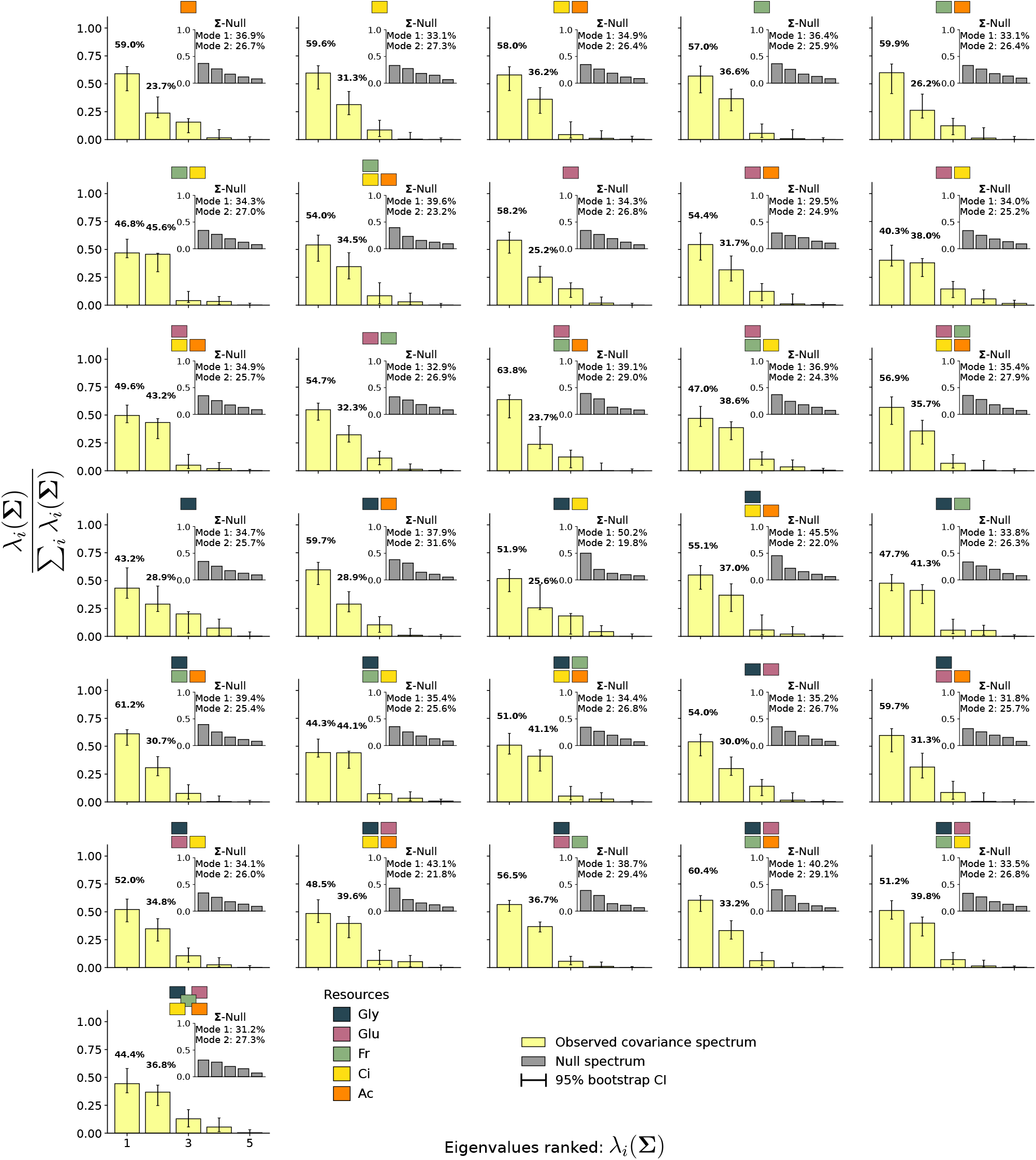
Low-rank covariance spectra across resource environments. For each resource environment, we approximated the covariance matrix of marginal effects, **Σ**, by retaining only first- and second-order WH coefficients. Since **Σ** = **M** − **ff** ^⊤^, Eq. (71), the second-order approximation gives **Σ** ≈ **HH**^⊤^, where **H** is the matrix of second-order WH coefficients. Unlike the uncentered second-moment matrix **M, Σ** removes the contribution of the mean marginal-effect vector and therefore isolates background-dependent variation. Bars show the normalized eigenvalue spectrum, *λ*_*i*_(**Σ**)*/* _*j*_ *λ*_*j*_(**Σ**), and error bars denote 95% bootstrap confidence intervals. Insets show the corresponding spectra obtained under the null model, **Σ**_null_. Across resource environments, the observed covariance spectra are dominated by one or two leading modes, indicating that the background-dependent component of marginal effects is also low-dimensional. Resource composition is indicated above each panel by colored rectangles denoting the carbon sources present in each medium.

**Figure S16:**
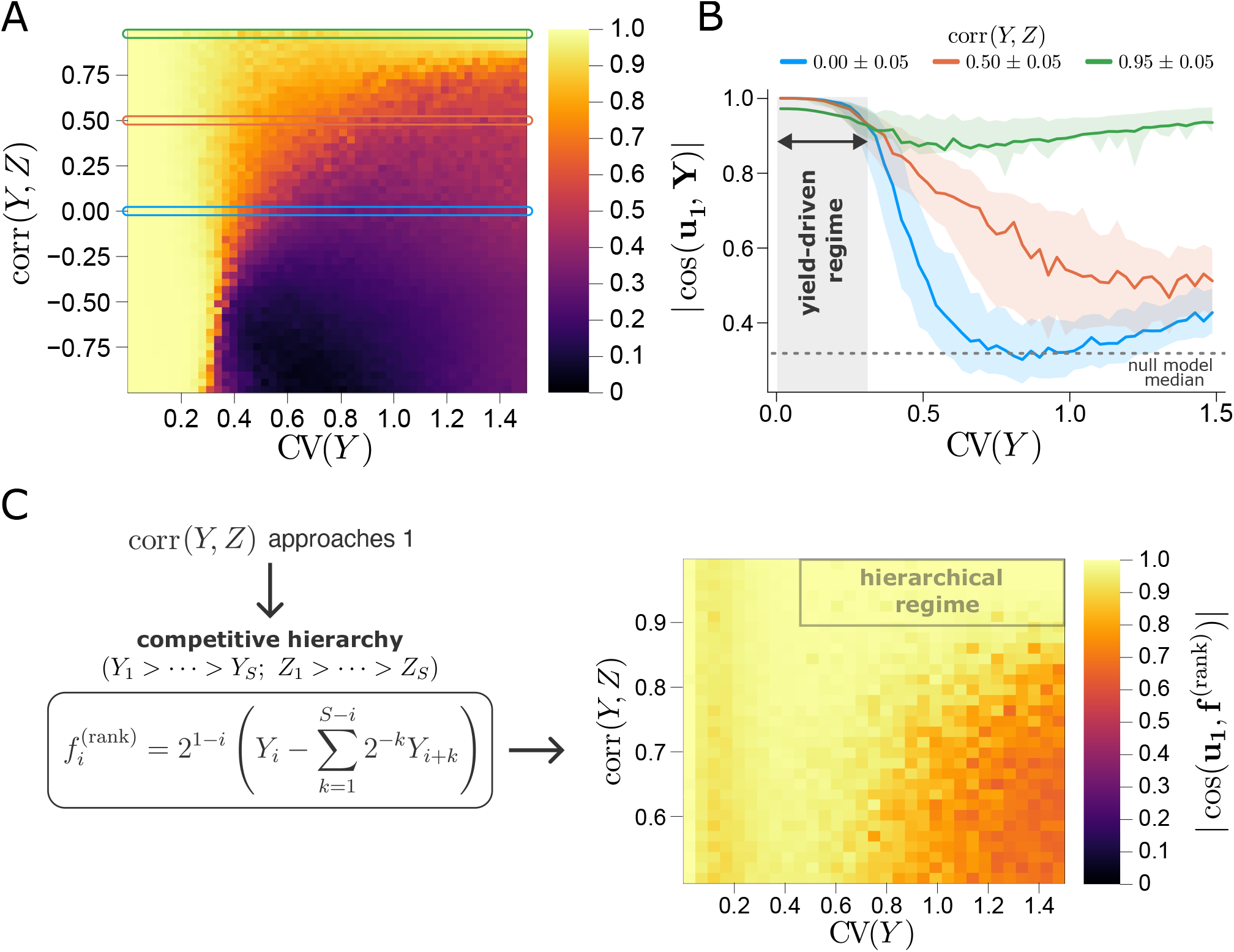
Consumer-Resource models predict the leading collective mode from species traits. Results from the numerical integration of the CRM with one public resource and 5 species (Eqs. (109)-(110); see Sec. S8.1 for details). Yields (*Y* ) and per-capita growth rates (*Z*) were sampled from uniform distributions of fixed means and variable standard deviations to produce coefficient of variations CV(**Y**) and CV(**Z**) varying from 0 to 1.5 in 51 equally-spaced steps. The traits vectors **Y** and **Z** were sampled 10 times for each value of CV(**Y**) and CV(**Z**), resulting in a total of 260100 simulated landscapes. The Pearson correlation coefficient between **Y** and **Z**, denoted as corr(**Y, Z**), was also computed. **(A)** Heatmap showing the median of | cos(**u**_**1**_, **Y**)|, the cosine similarity between the leading collective mode **u**_**1**_ and the yields vector **Y**, against CV(**Y**) and corr(**Y, Z**). Simulations confirm the yield-driven regime at low CV(**Y**) (independent of growth rates) predicted in Eq. (122). **(B)** Median and IQR of |cos(**u**_1_, **Y**)| are shown against CV(**Y**) for uncorrelated (corr(**Y, Z**) = 0.00 ± 0.05), moderately correlated (corr(**Y, Z**) = 0.50 ± 0.05), and strongly correlated (corr(**Y, Z**) = 0.95 ± 0.05) yields and growth rates. The median of the cosine similarity between two random normalized 5-dimensional vectors (see Sec. S8.2) is also reported for comparison. **(C)** As corr(**Y, Z**) approaches 1, the system enters a regime of competitive hierarchy where the leading eigenmode **u**_1_ aligns with the vector of focal effects **f** ^(rank)^, computed in Eq. (130) and fully determined by yields.

**Figure S17:**
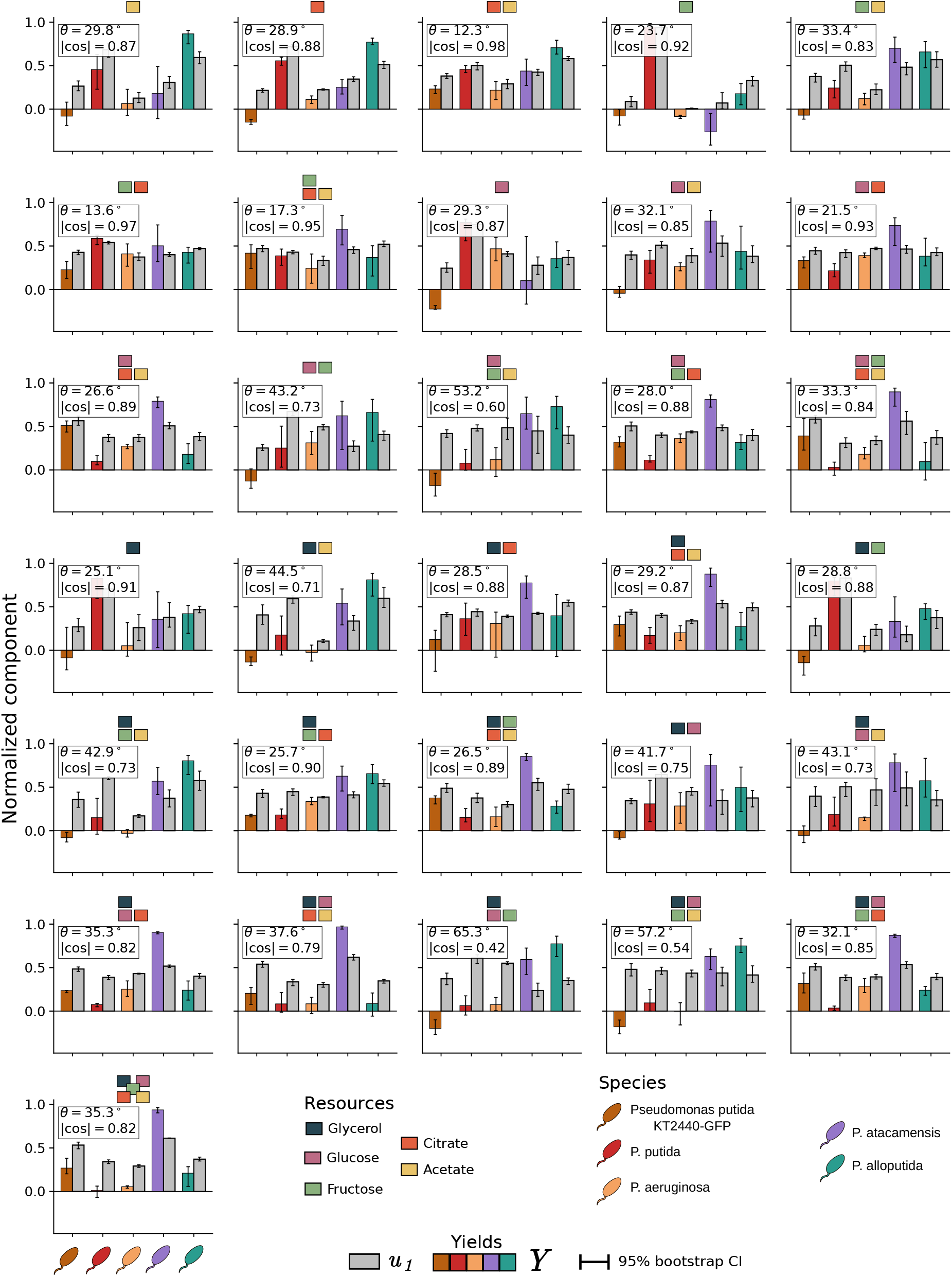
Yield productivity provides a proxy for the leading collective mode. Each panel corresponds to a different resource environment and compares the normalized components of the leading collective mode, **u**_1_, with the normalized yield productivity vector, **Y** ≈ **B**_mono_, computed from the monoculture biomass as proxy. The sign of **u**_1_ was chosen to maximize its overlap with **Y.** Insets report the sign-invariant angle, 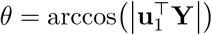, and the corresponding absolute cosine similarity, 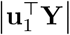. Error bars denote 95% bootstrap confidence intervals.

**Figure S18:**
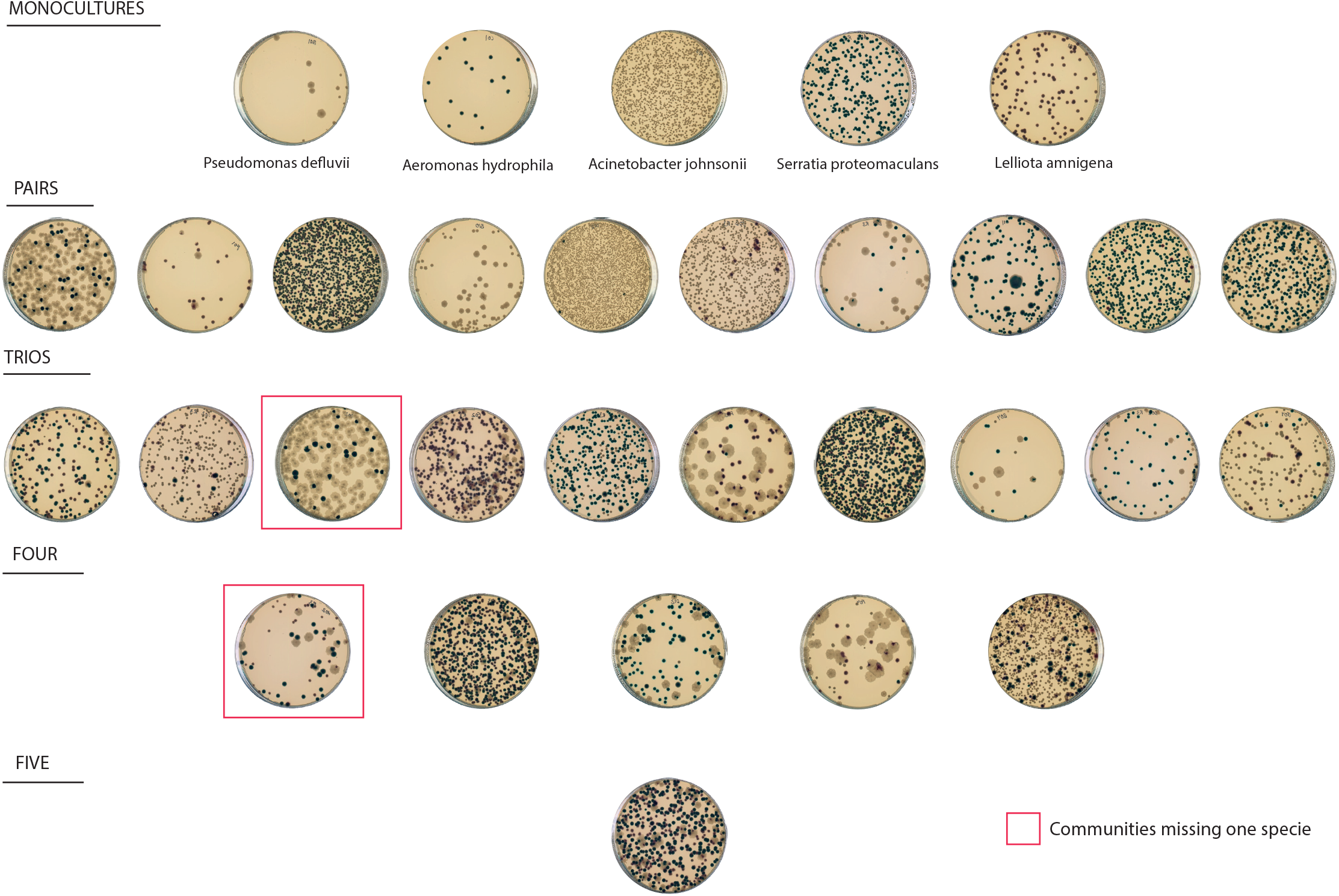
Agar plate images of all five-species combinations after the 10-day experiment. The 31 agar plates each contain a different species combination, including monocultures, pairs, trios, four-species, and five-species communities. Red squares indicate the only two final communities in which one species was missing after 10 days.

**Figure S19:**
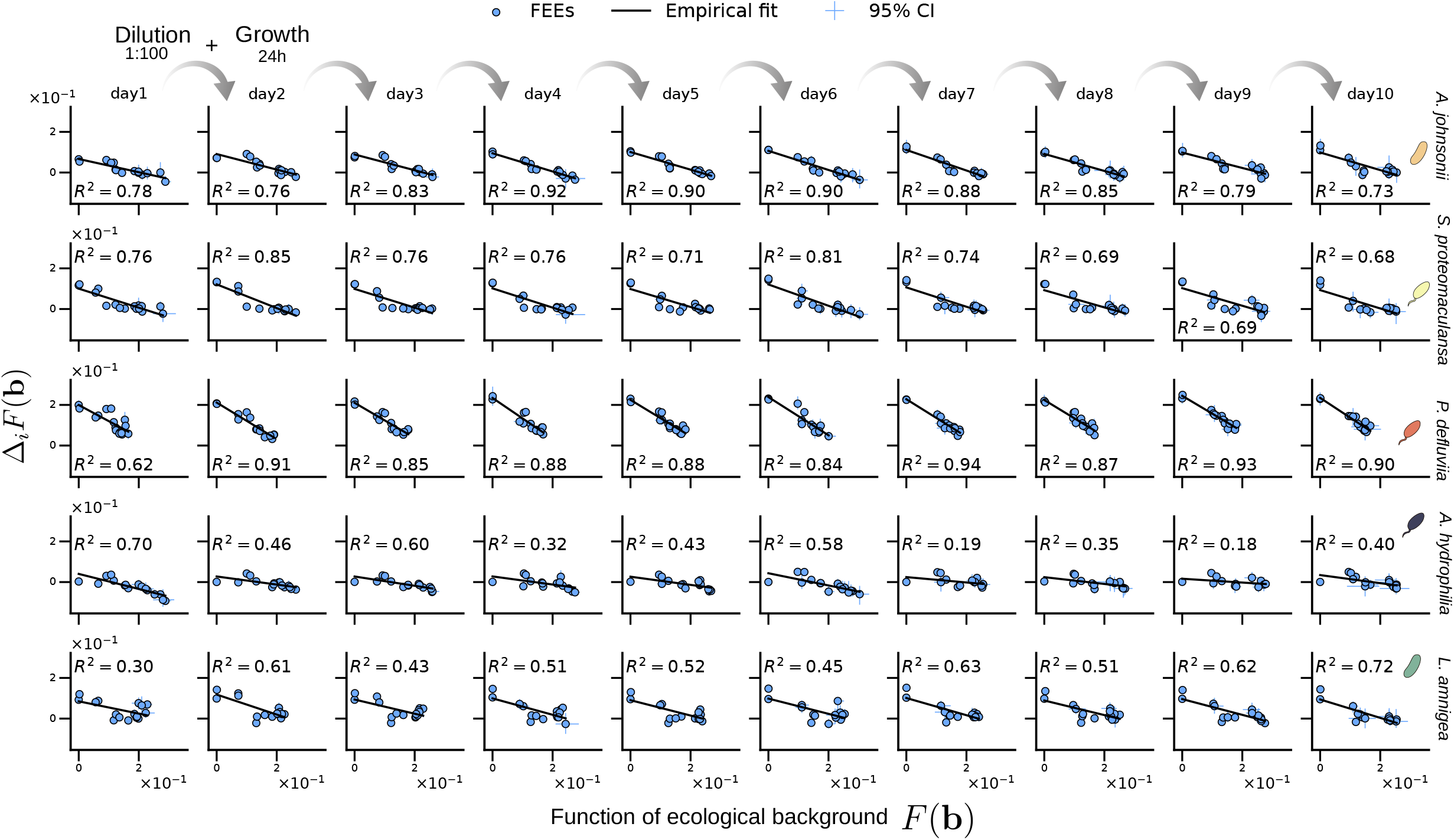
Functional Effect Equations remain predictable during serial dilution. Functional Effect Equations (FEEs) are shown for each of the ten daily passages of the serial-dilution experiment. Columns correspond to passages, and rows to focal species. In each panel, the functional effect of adding species *i*, Δ_*i*_*F* (**x**), is plotted against the function of the corresponding background community, *F* (**x**). Points show measured functional effects, error bars denote 95% bootstrap confidence intervals, and solid black lines show empirical linear fits. The coefficient of determination, 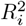, is reported in each panel. Species names are indicated on the right.

**Figure S20:**
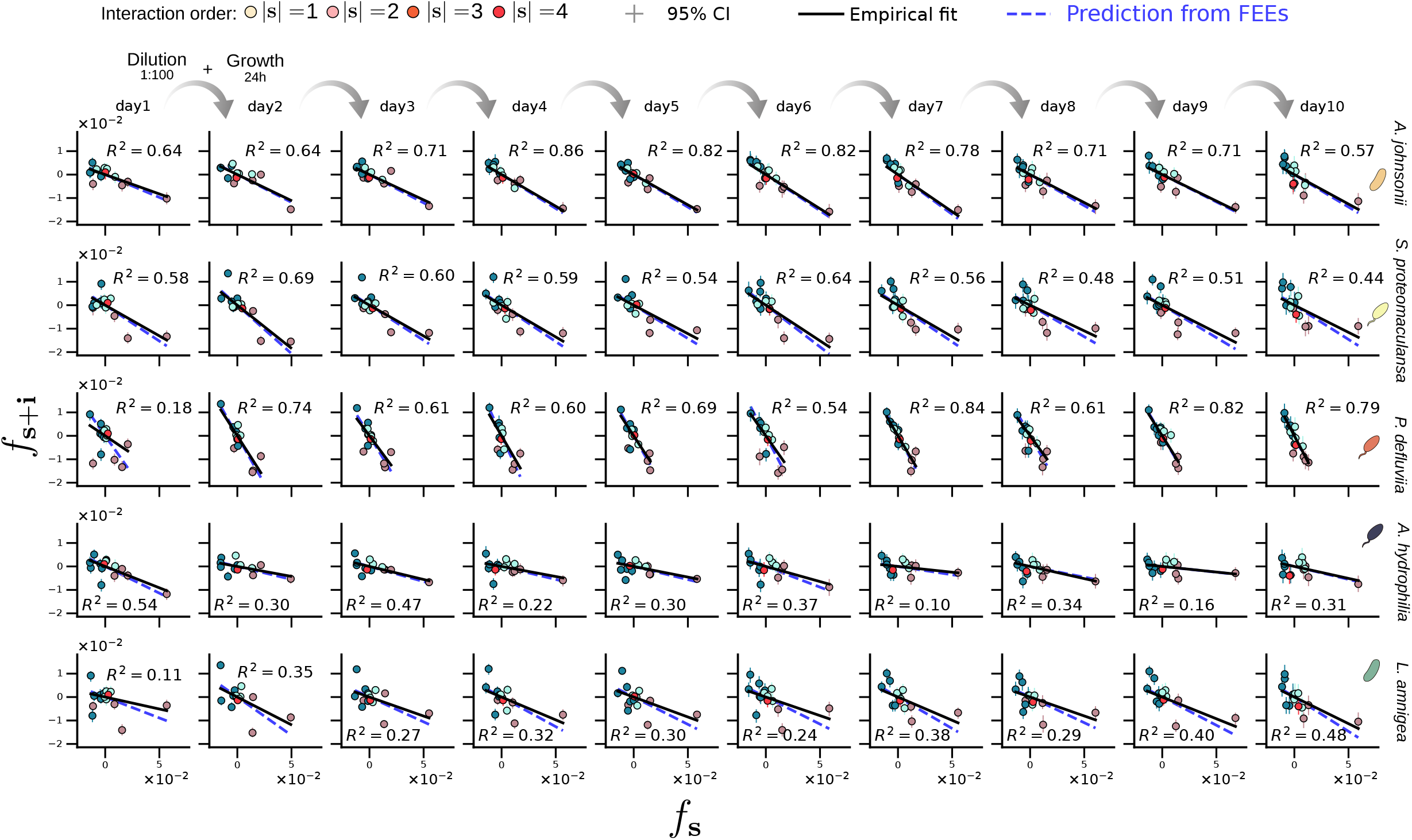
WH interaction coefficients retain simple cross-order relationships during serial dilution. Cross-order relationships among WH interaction coefficients are shown for each of the ten daily passages of the serial-dilution experiment. Columns correspond to passages, and rows to focal species. In each panel, WH coefficients involving focal species *i, f*_**s**+**i**_, are plotted against the corresponding coefficients one order below, *f*_**s**_, for subsets **s** not containing species *i*. Points are colored by interaction order, and error bars denote 95% bootstrap confidence intervals. Solid black lines show empirical linear fits, whereas dashed blue lines indicate the slopes predicted from the corresponding fitted FEE slope *b*_*i*_, using *c*_*i*_ = *b*_*i*_*/*(2 + *b*_*i*_) as given by Eq. (15). The coefficient of determination of the interaction-space fit, 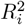, is reported in each panel. Species names are indicated on the right.

**Figure S21:**
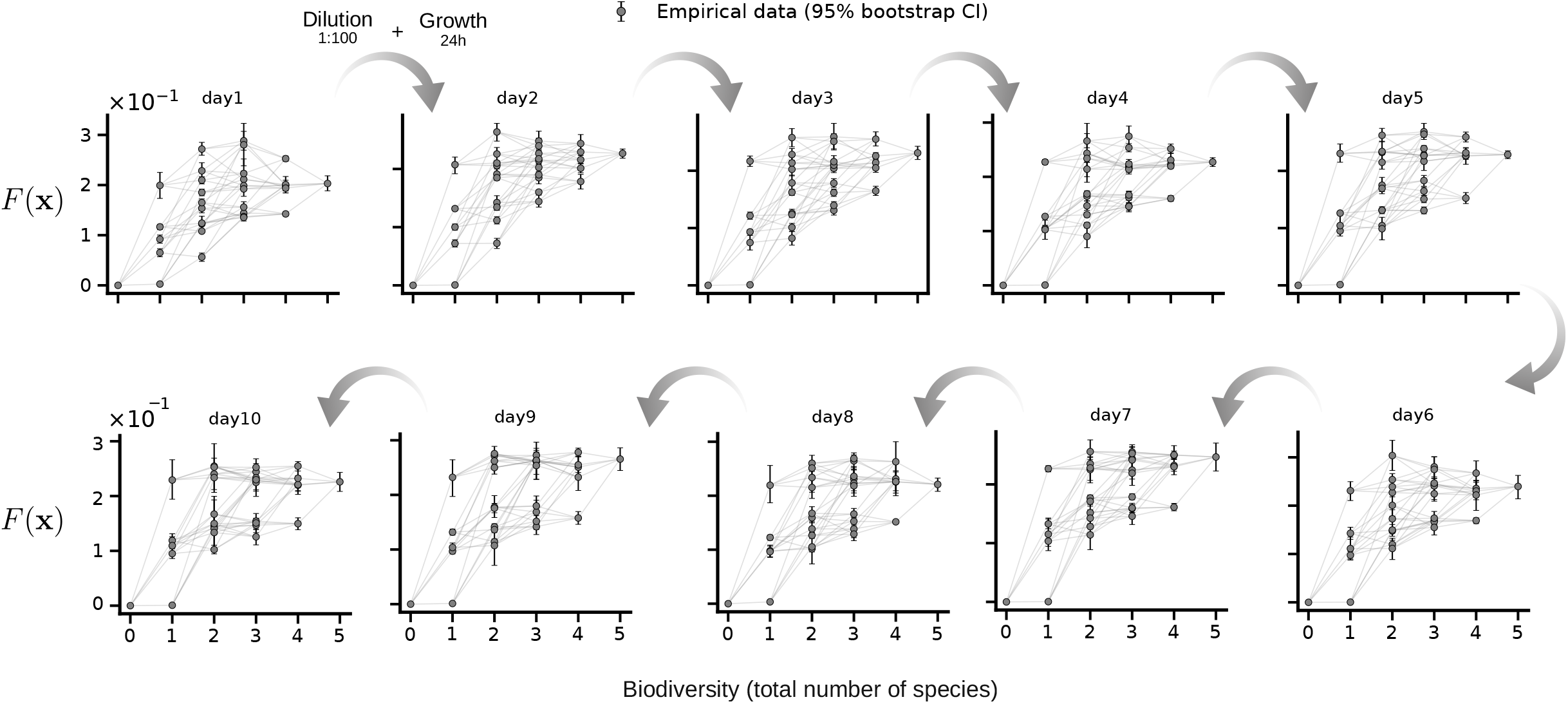
Biomass landscapes projected onto community biodiversity during serial dilution. For each of the ten daily passages, the biomass community-function landscape was projected onto community biodiversity, defined as the number of species present in the initial assemblage. Points show measured biomass values, *F* (**x**), and error bars denote 95% bootstrap confidence intervals. Light grey lines connect communities that differ by the addition or removal of a single species, preserving the adjacency structure of the community-composition hypercube. The relationship between biomass and biodiversity remains broadly conserved across passages, indicating that the coarse organization of the biomass landscape persists during population dynamics.

**Figure S22:**
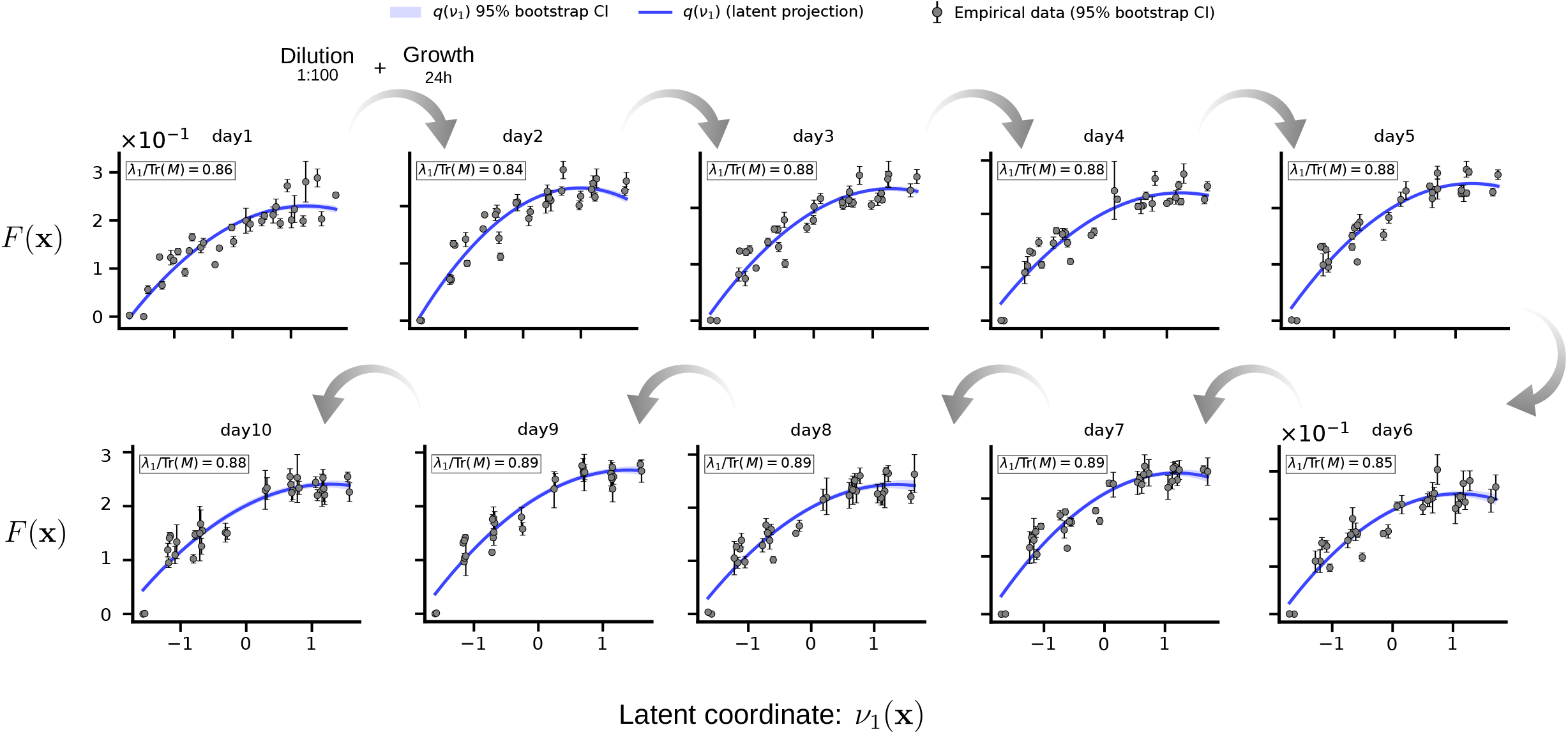
Biomass landscapes retain a stable projection onto the leading collective coordinate during serial dilution. For each of the ten daily passages, the biomass community-function landscape was projected onto its leading latent coordinate,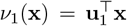. Points show measured biomass values, *F* (**x**), and error bars denote 95% bootstrap confidence intervals. Solid blue lines show the latent-coordinate prediction, *q*(*ν*_1_), and shaded blue regions denote the corresponding 95% bootstrap confidence intervals. The relationship between biomass and *ν*_1_(**x**) remains stable across passages, indicating that the dominant collective axis organizing biomass production is preserved during population dynamics.

**Figure S23:**
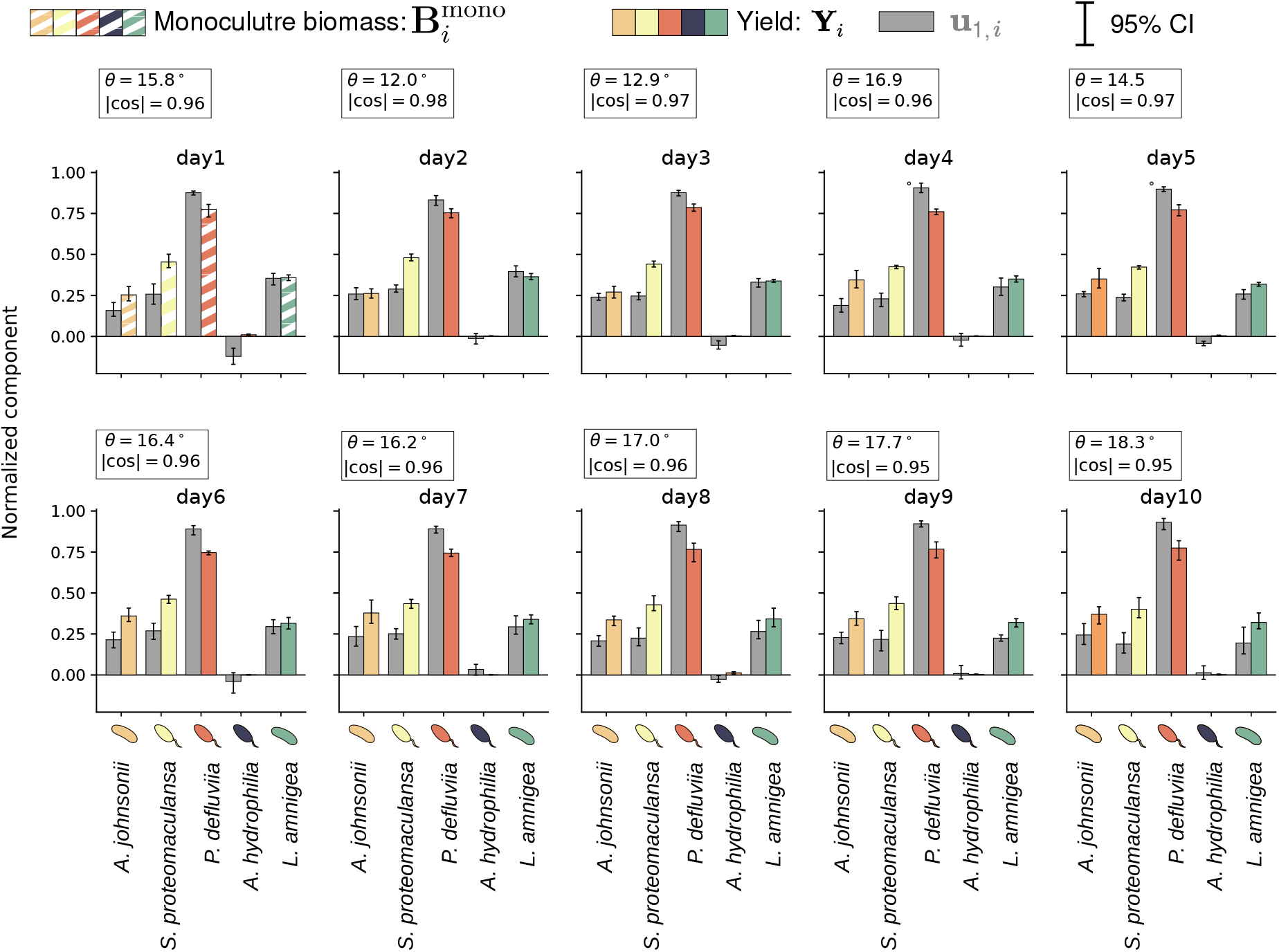
The leading productivity mode remains aligned with species-level yield estimates during serial dilution. For each passage, bars compare the normalized components of the leading collective mode, **u**_1_, with the corresponding normalized species-level productivity vector. On day 1, species productivity is approximated by monoculture biomass, **Y** ≈ **B**_mono_, because the yield estimator requires measurements from two consecutive passages. For days 2–10, species yields are estimated from monoculture biomass measured in consecutive passages using Eq. (1). Error bars denote 95% bootstrap confidence intervals. Insets report the sign-invariant angle, 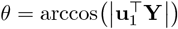, and the corresponding absolute cosine similarity, 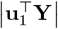, between the leading collective mode and the productivity vector used for each passage. The consistently high similarity indicates that the dominant collective mode remains aligned with species-specific productivity throughout the serial-dilution experiment.

**Figure S24:**
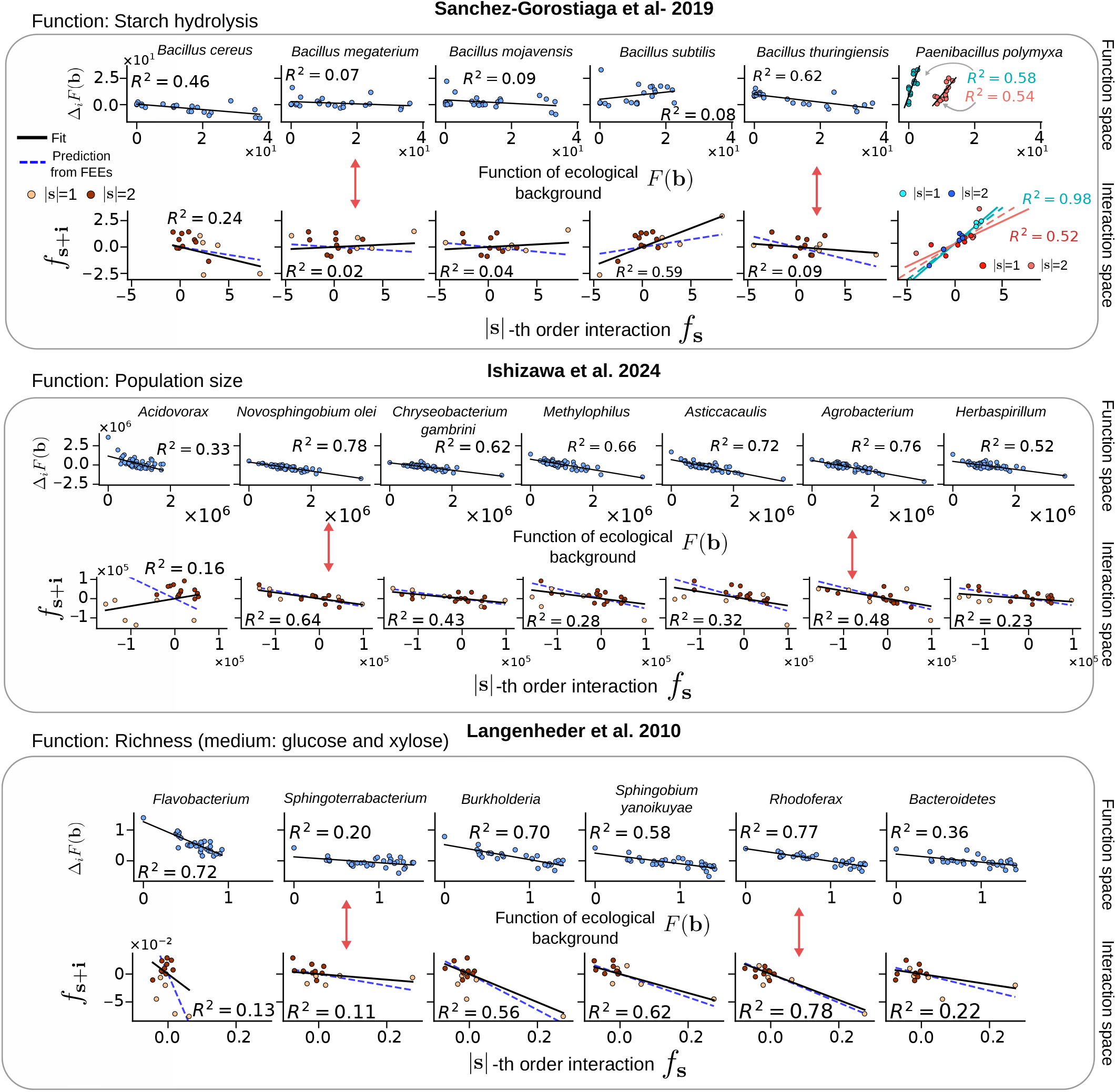
Functional Effect Equations predict cross-order dependencies across published community-function landscapes. Top, biomass data from Sanchez-Gorostiaga et al. (2019) [20]. Middle, population-size data from Ishizawa et al. (2024) [15] Bottom, richness data from Langenheder et al. (2010) [17]. Each column corresponds to one focal taxon. Upper rows show the functional effect, Δ_*i*_*F* (**b**), as a function of background function, *F* (**b**). Lower rows show *f*_**s**+**i**_ against the corresponding lower-order coefficient *f*_**s**_. Point colors indicate interaction order |**s** |. Black lines show empirical linear fits. Blue dashed lines show FEE predictions. Values report *R*^2^. For *Paenibacillus polymyxa* in Sanchez-Gorostiaga et al. (2019), the FEE separates into two branches depending on whether *Bacillus thuringiensis* is present or absent in the ecological background. Only first- and second-order relationships are shown because higher-order coefficients are difficult to estimate reliably without replication [3].

**Figure S25:**
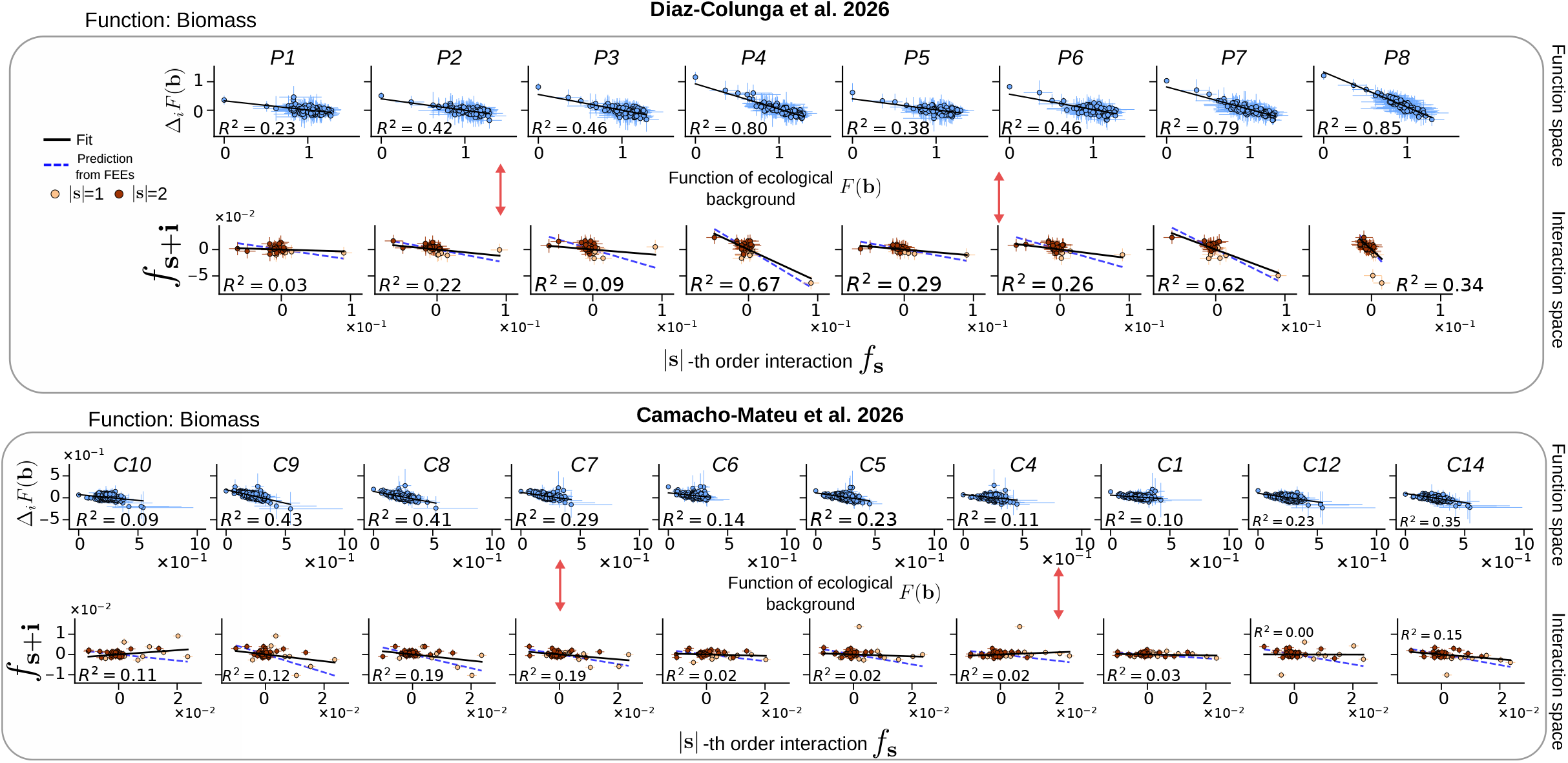
Functional Effect Equations predict cross-order dependencies among ecological interactions in published biomass landscapes. Top, Diaz-Colunga et al. (2026) [8]. Bottom, Camacho-Mateu et al. (2026) [3]. Each column corresponds to one focal species. Upper rows show the functional effect, Δ_*i*_*F* (**b**), as a function of background biomass, *F* (**b**). Lower rows show interaction coefficients involving the focal species, *f*_**s**+**i**_, against the corresponding lower-order coefficients, *f*_**s**_. Point colors indicate interaction order |**s** |. Black lines show empirical linear fits. Blue dashed lines show the slopes predicted from the corresponding FEEs. Values report the coefficient of determination, *R*^2^. Red arrows highlight representative correspondences between function space and interaction space. Stronger FEEs are generally associated with more pronounced cross-order organization of interaction coefficients.

**Figure S26:**
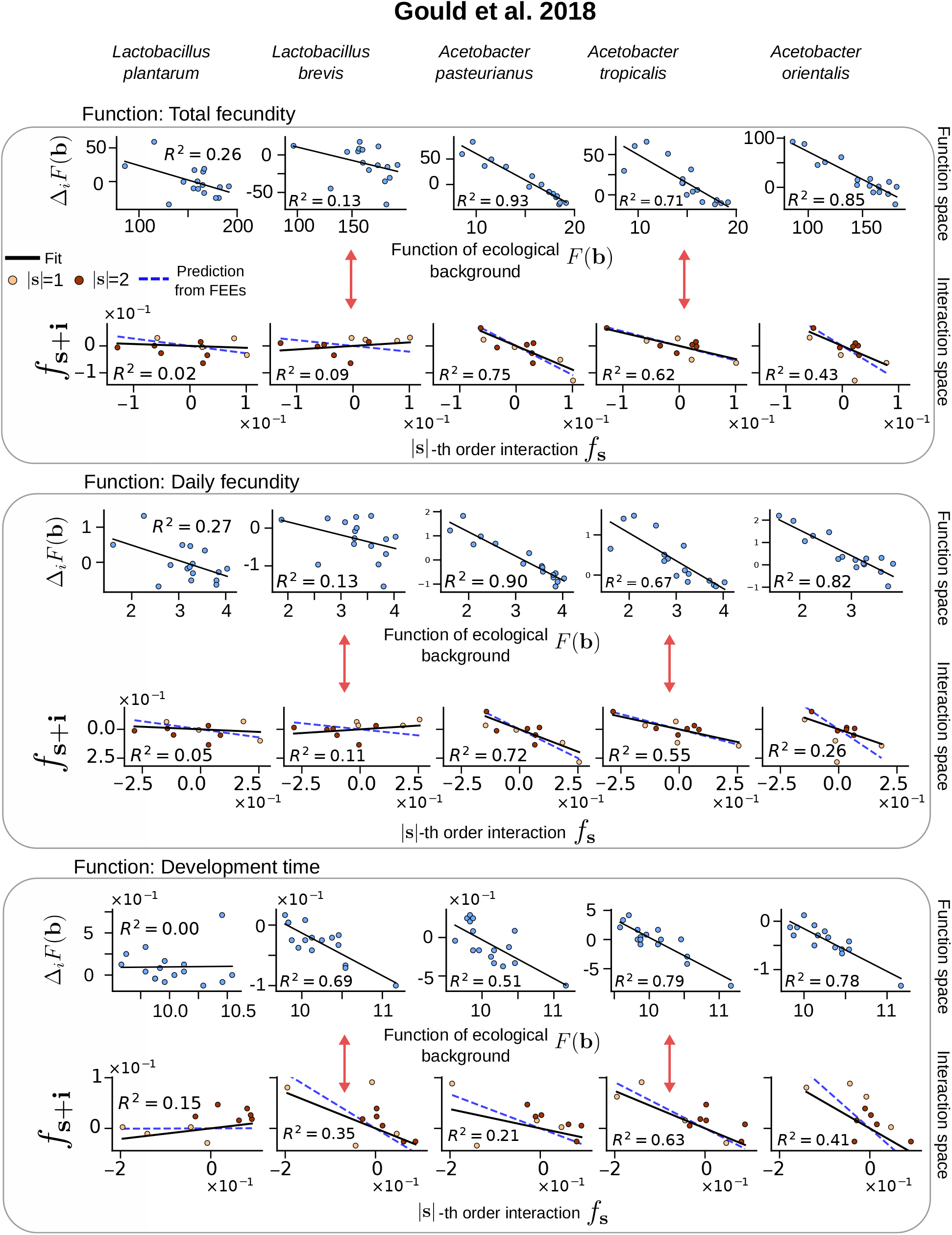
Functional Effect Equations and cross-order relationships among WH coefficients in the *Drosophila* microbiome landscape of Gould et al. (2018). Data are from Gould et al. [11]. Columns correspond to the five bacterial species and rows to total fecundity, daily fecundity, and development time. Upper panels show the functional effect of focal species *i*, Δ_*i*_*F* (**b**), as a function of the community function in ecological background **b.**Lower panels compare WH coefficients involving the focal species, *f*_**s**+**i**_, with the corresponding lower-order coefficients, *f*_**s**_, where |**s**| denotes interaction order. Solid black lines show empirical fits, dashed blue lines show slopes predicted from the FEEs, and *R*^2^ values quantify linear fit quality.

**Figure S27:**
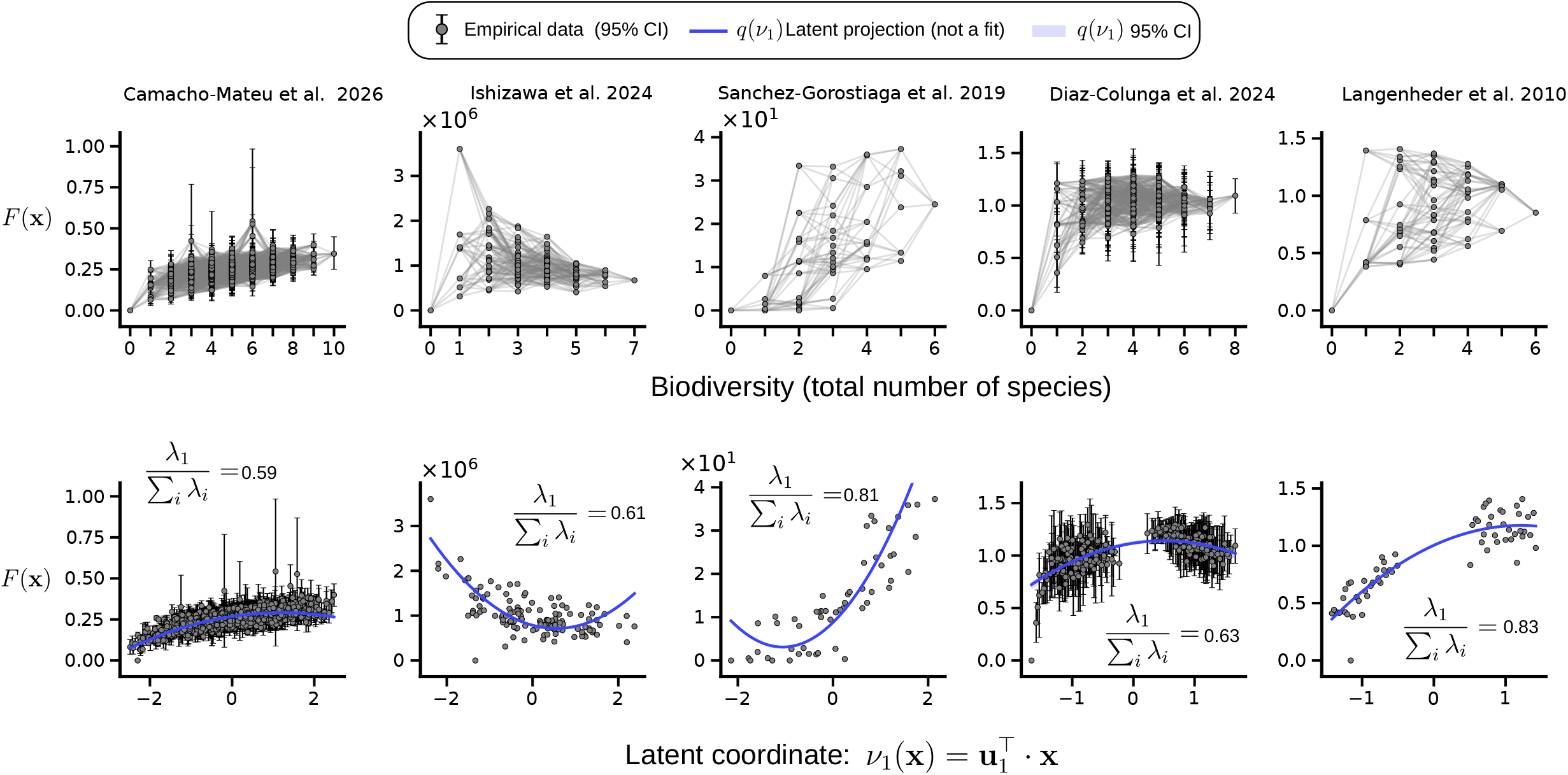
One-dimensional latent-coordinate projections of previously published community-function landscapes. Five fully sampled community-function landscapes from previously published datasets were analyzed using the same framework applied to our experimental data (Table T2). Each row corresponds to one dataset. Left panels show the community function, *F* (**x**), as a function of biodiversity, whereas right panels show *F* (**x**) projected onto the leading latent coordinate,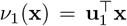. Black points show empirical measurements, and vertical error bars denote 95% bootstrap confidence intervals when replicate measurements are available. Solid blue lines show the predicted response, *q*(*ν*_1_), and shaded blue regions denote the corresponding 95% bootstrap confidence intervals. Labels report the fraction of total spectral weight captured by the leading mode, *λ*_1_*/*Tr(**M**).

**Figure S28:**
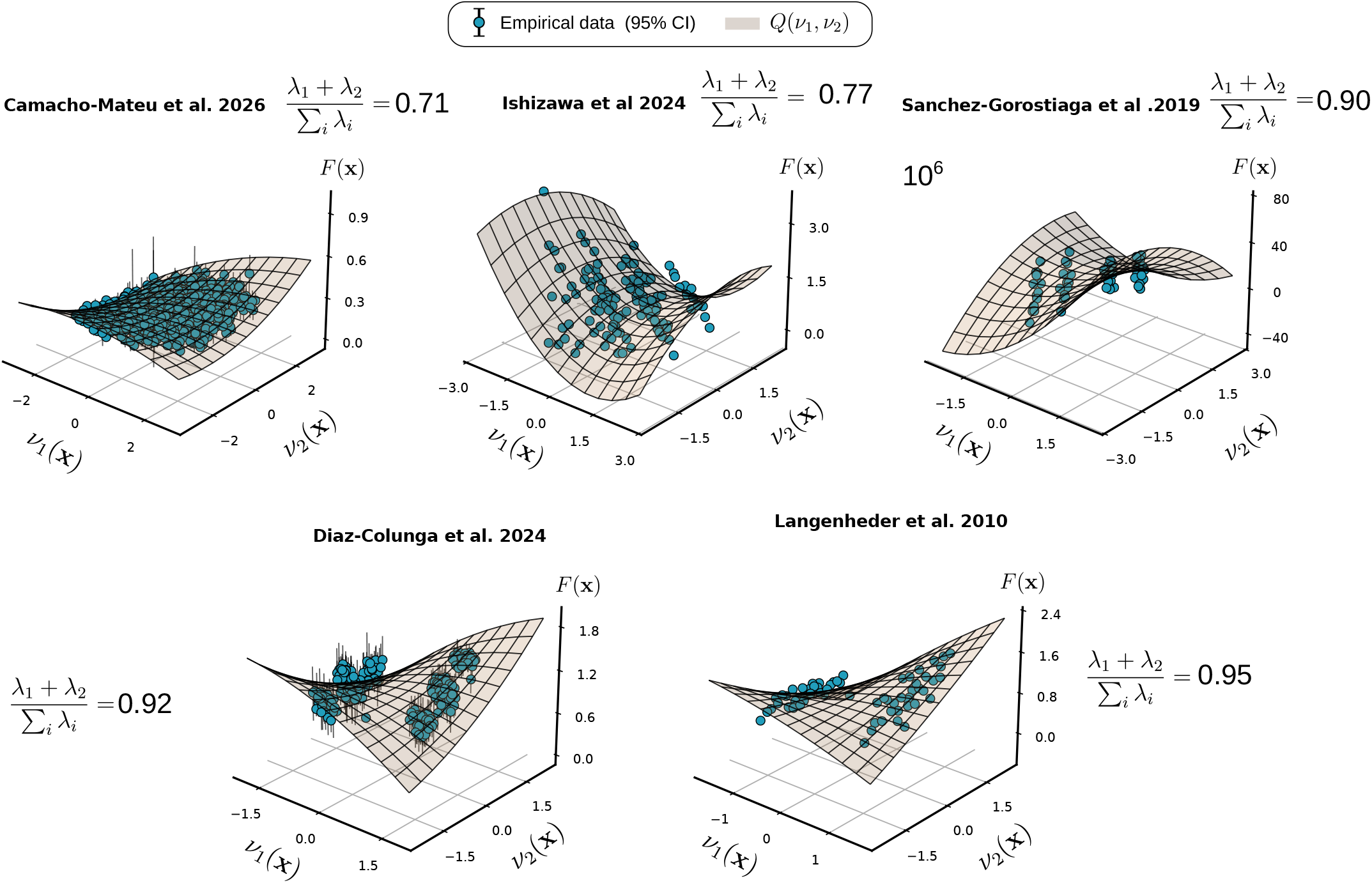
Two-dimensional latent-coordinate manifolds of previously published community-function landscapes. The same five datasets shown in Fig. S27 were projected onto their first two latent coordinates, 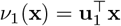 and 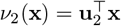. Points show empirical measurements, and vertical error bars denote 95% bootstrap confidence intervals when replicate measurements are available. Surfaces show the predicted quadratic latent manifold, *Q*(*ν*_1_, *ν*_2_). Labels report the fraction of total spectral weight captured by the first two modes, (*λ*_1_ + *λ*_2_)*/*Tr(**M**).

**Figure S29:**
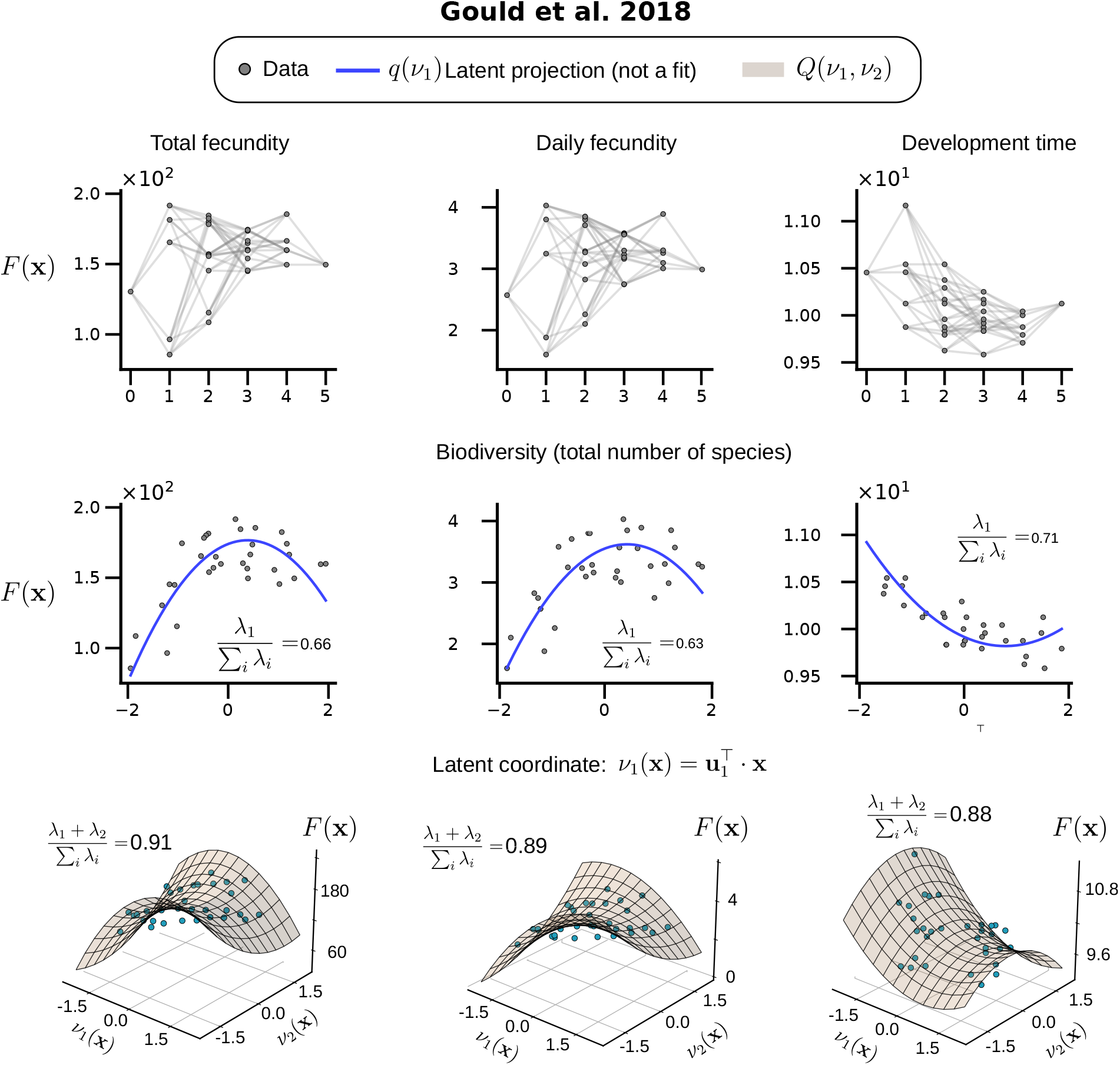
Low-dimensional latent projections of host fitness landscapes in the *Drosophila* microbiome dataset of Gould et al. (2018). Data are from Gould et al. [11]. Columns correspond to total fecundity, daily fecundity, and development time. Top row, measured community function *F* (**x**) plotted against bacterial biodiversity, defined as the total number of species in each community. Grey points denote observed community functions, and gray lines connect communities that differ by the addition or removal of a single bacterial species. Middle row, the same landscapes projected onto the leading latent coordinate, 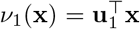, where **u**_1_ is the dominant eigenvector of the second-moment matrix of marginal effects. Blue curves show the one-dimensional latent projection *q*(*ν*_1_) obtained from the second-order WH representation; these curves are determined by the spectral projection and are not fitted directly to the plotted data. Insets report the fraction of total spectral weight captured by the leading mode, *λ*_1_*/*Tr(**M**). Bottom row, two-dimensional representations of each landscape in the coordinates (*ν*_1_, *ν*_2_). Cyan points show the measured functions, and the surfaces show the quadratic latent projection *Q*(*ν*_1_, *ν*_2_). The reported values (*λ*_1_ + *λ*_2_)*/*Tr(**M**) indicate the fraction of spectral weight captured by the first two collective modes. Despite the weak organization of the landscapes by species biodiversity, one or two ecological collective modes capture most of the coordinated variation in marginal effects and provide a compact representation of all three host traits.

**Figure S30:**
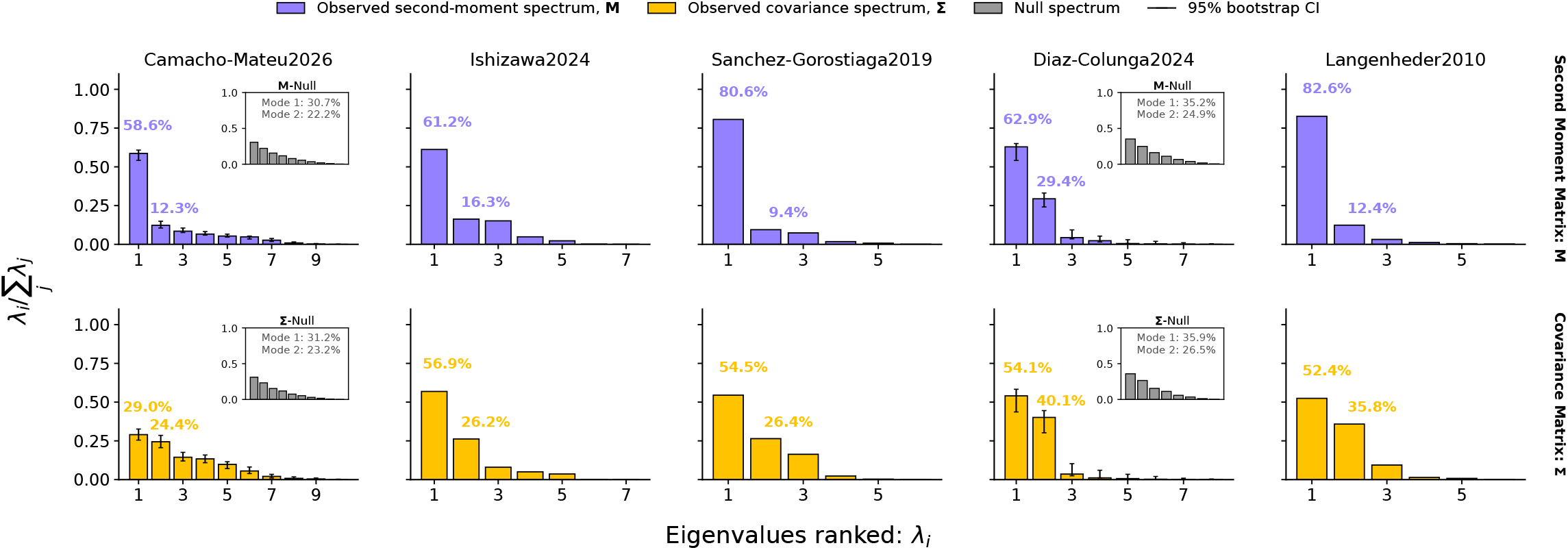
Second-moment and covariance eigenspectra of previously published community-function landscapes. Five fully sampled community-function landscapes from previously published datasets were analyzed using the same spectral framework applied to our experimental data (Table T2). Columns correspond to datasets. Top panels show the normalized eigenspectrum of the second-moment matrix of marginal effects, **M**, whereas bottom panels show the normalized eigenspectrum of the centered covariance matrix, **Σ.** Bars report *λ*_*i*_*/* ∑_*j*_ *λ*_*j*_, with eigenvalues ranked in decreasing order; error bars denote 95% bootstrap confidence intervals when replicate measurements are available. Labels above the first two bars indicate the fractions of total spectral weight captured by the leading two modes. Insets show the corresponding mean null eigenspectra and report the spectral weight of the first two null modes. Across datasets, the spectra of **M** are strongly concentrated in one or two leading modes. The spectra of **Σ** are also concentrated relative to the null expectation, indicating that the background-dependent variation of marginal effects is itself organized along a small number of collective directions.

**Figure S31:**
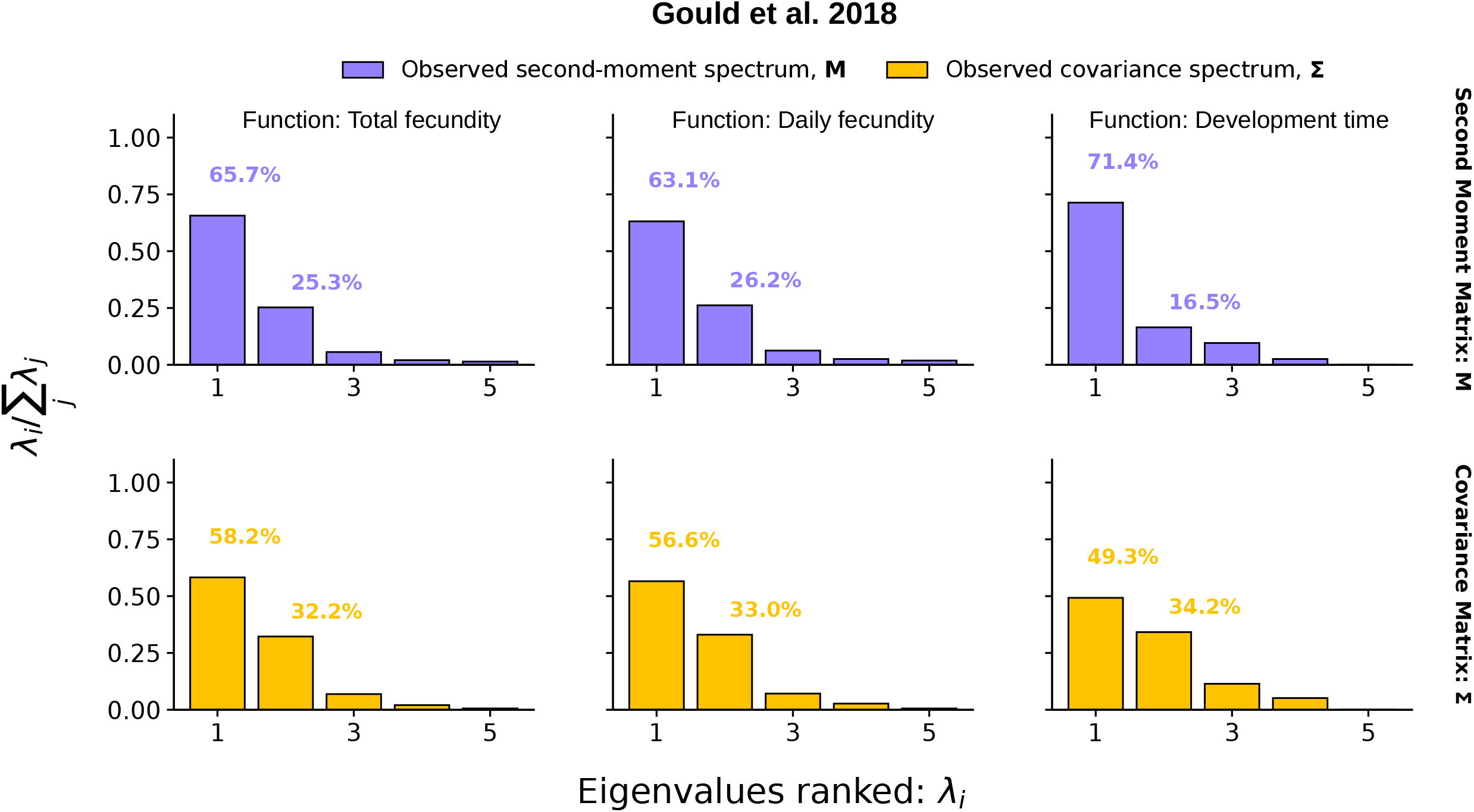
Second-moment and covariance eigenspectra of the *Drosophila* microbiome landscape of Gould et al. (2018). Data are from Gould et al. [11].

